# Accessory genes define species-specific pathways to antibiotic resistance

**DOI:** 10.1101/2023.08.09.552647

**Authors:** L. Dillon, NJ. Dimonaco, CJ. Creevey

## Abstract

**Background:** The rise of antimicrobial resistance (AMR) is a growing concern globally and a deeper understanding of AMR gene carriage vs usage is vital for future responses to reduce the spread of AMR. Identification of AMR phenotype by laboratory-based assays are often hindered by difficulties in establishing cultures. This issue could be resolved by rapid computational assessment of an organism’s genome, however, AMR gene finder tools are not intended to infer AMR phenotype which is likely to be a product of multiple gene interactions.

**Methods:** To understand the importance of multi-gene interactions to the relationship between AMR genotype and AMR phenotype, we applied machine learning approaches to 16,950 genomes from microbial isolates representing 28 different genera with 1.2 million corresponding laboratory-determined MICs for 23 different antibiotics. We then elucidated the genomic paths to phenotypic antimicrobial resistance with the aim of allowing for the development of rapid determination of AMR phenotype from genomes or even whole microbiomes.

**Results:** The application of machine learning models resulted in a >1.5-fold increase in average prediction accuracy of AMR phenotype across the 23 antibiotic models. Interpretation of these models revealed 528 distinct genomic pathways to phenotypic resistance, many of which were species-specific and involved genes which have not previously been associated with AMR phenotype. This is the first study to demonstrate the utility of machine learning models in the prediction of AMR phenotype for a wide range of clinically relevant organisms and antibiotics. This could be applied as a rapid and affordable alternative to culture-based techniques, estimating taxonomy in addition to AMR phenotype, and providing real-time monitoring of multi-drug resistant pathogens.

**Availability and implementation:** *Contact:*

*View supplementary information at this link:* https://osf.io/cj4bq/?view_only=c0ee87b7609543b688953089be4c376f See Code Availability for scripts used.

## INTRODUCTION

The overuse and misuse of antibiotics has escalated the rate at which many bacteria have evolved resistance to multiple antibiotics [9, 37], including last-resort treatments [2]. This has led to a growing prevalence of antimicrobial-resistant infections worldwide [27], which can be challenging to treat [7]. This has caused antimicrobial resistance (AMR) to become an increasing burden on society from a global health, agricultural and financial perspective [1, 20, 42]. If the rate of AMR continues as projected, it is estimated that by the year 2050, there will be *>* 10 million deaths annually as a result of AMR-related infections [26].

AMR phenotype is typically distinguished through laboratory-based approaches such as broth microdilution, E-tests or disk diffusion assays [4]. However, it typically takes 2-4 days to allow time to culture the bacteria and then complete the test [5]. More rapid testing is available through automated instruments for antibiotic susceptibility testing, such as commercial automated antimicrobial susceptibility tests (i.e. Vitek 2 system and Microscan WalkAway) [4, 24] or isothermal microcalorimetry accurately determine MIC values (i.e. Symcel) [41], which is often used in hospital environments. However, while these assays are usually a good estimate of AMR phenotype in culture, this does not always translate to clinical settings. This is further complicated by the difficulty in culturing many organisms, especially when assessing species directly from microbiome samples [36]. Besides the importance of understanding the role of AMR phenotype in microbiomes from an AMR reservoir perspective [37], it also has the potential to reveal the mechanisms underpinning AMR-driven dysbiosis [31] within humans and animals and potentially aid us to prevent disease whilst concomitantly slowing the spread of AMR.

Recently, computational methods to identify AMR-causing genes in genomic data have become widely available [6, 30, 19] and are often used to assess the potential antibiotic resistance phenotype of an organism [37] or even entire microbiomes [46]. These AMR gene finder tools run relatively quickly, especially compared to laboratory-based assays. Still, different tools can provide varying results [16], likely driven by differences in the databases that they use to detect AMR genes and the varying methods to extract the AMR genes [14]. AMR gene finder tools are also prone to error, for example, when closely related genes may be predicted as resistance genes incorrectly [5]. Very often microbiomes harbour AMR genes even when antibiotic usage is absent [28, 48, 18]. Why these bacteria harbour AMR genes within the microbiome is unclear. Several AMR genes have been reported to have alternative functions, such as transporters [11], yet this is not the case for all.

Most importantly, AMR gene finder tools predictions represent what is likely in many cases to be a simplified concept of the mechanisms underpinning the presentation of AMR phenotype. There is the assumption that a single gene or mutation is responsible for the phenotypic expression of AMR and gene finder tools do not take into account other genes which may be required to confer resistance to an organism [25]. We refer to these non-classical AMR genes that are important to the presentation of AMR phenotype as “accessory” genes.

Some previous studies have attempted to use machine learning as a way of predicting the AMR phenotype from genotype [33, 34]. These studies have had several limitations such as only studying a specific species, and/or using a single antibiotic [32, 29, 45, 44] or using non-interpretable methods such as a neural network [3], thereby limiting our ability to understand the biological processes involved. Using a more interpretable method such as decision trees, applied to a wide range of taxa across multiple antibiotics has the potential to provide a unique biological understanding of antibiotic resistance and allow the identification of accessory genes associated with alternative “paths” to phenotypic resistance.

To address this, we determined the role of “accessory” genes in the presentation of an AMR phenotype. Our hypothesis is that focusing solely on classic AMR genes misses vital information needed to evaluate AMR phenotypes accurately. We address this through the application of multiple Machine Learning (ML) models to a dataset of 16,950 genomes from microbial isolates representing 28 different genera with 1.9 million corresponding laboratory-determined MICs for 79 different antibiotics. This data was filtered by matching to EUCAST breakpoints and to ensure more balanced datasets according to AMR phenotype (see Methods and Materials: Data for Analysis for further details). We then elucidate the genomic paths (combinations of genes presence and absence to reach a phenotype) to phenotypic antimicrobial resistance that are shared or unique to species or antibiotics with the aim of allowing for the development of rapid determination of AMR phenotype from genomes or even whole microbiomes.

## METHODS AND MATERIALS

All scripts and files mentioned in the text can be found at https://github.com/LucyDillon/AMR_ML_paper/tree/main. This includes all bash scripts to analyse data using tools and details of how gene counts for RGI and Eggnog gene families were calculated.

Supplementary files and additional data can be found at: https://osf.io/cj4bq/?view_only= c0ee87b7609543b688953089be4c376f.

### Data for Analysis

Using the PATRIC command line interface (version 1.034 - now known as BV-BRC) [13, 35], 16,950 bacterial genomes from isolates of known taxonomy with 1,249,188 corresponding laboratory-determined MIC values were sourced. The genomes used in this study can be found using a wget command called: PATRIC_genomes.sh using the input: genome_ids.txt. For each genome, the AMR genotype was determined using the Resistance Gene Identifier (RGI) tool v5.1.1 [23] with the CARD database v3.1.1 [30] using the default parameters and the whole genome sequence from the genome as input. The CARD database includes acquired resistance and resistance due to mutations.

Each predicted AMR gene in each genome was then associated with the specific antibiotic(s) to which it was listed as conferring resistance to using the information in the CARD database. The MIC values were categorised into ‘Susceptible’ or ‘Resistant’ using EUCAST breakpoints (Jan 2021 release) [15] which are taxonomic-specific MIC values that can differ between species. The MICs were categorised into the respective EUCAST breakpoints using custom Python scripts (OG_RGI_analysis.py, Logistic_regression_RGI.py, RGI_specific_analysis.py, RGI_all_analysis.py, and Eggnog_analysis.py). Any MIC values that fell outside the EUCAST definition of “susceptible” or “resistant” for any specific species were removed from the analysis. In the case that a genome had >1 MIC values for the same antibiotic, the average was calculated and then compared to the EUCAST breakpoints. This resulted in 5,990 genomes, 19 Genera, with 47,711 EUCAST classified MICs for subsequent analysis (28,480 resistant and 19,231 susceptible MICs). Details of the number of each Genus for each antibiotic model can be found in Supplementary Table 1.

### Analysis of AMR genotype to phenotype relationship

In this study, we used several techniques to further understand the relationship between AMR genotype and AMR phenotype. The models used are binary classifiers (either classifying as susceptible or resistant) which although makes for a simpler model, excludes the use of intermediate resistance or more complex conditions such as persistence or tolerance. To predict the AMR genes present within each genome, we used RGI, a commonly used AMR gene finder tool. We evaluated four phenotype prediction approaches using linked laboratory-determined resistance/susceptibility profiles against a range of antibiotics. We first tested a naive prediction of AMR phenotype using the presence/absence of AMR genes and the antibiotics to which the genes were listed as conferring resistance in the RGI database. Secondly, we tested the application of a basic logistic regression model to the AMR gene presence-absence data. Finally, we tested the application of four machine-learning approaches to predict AMR phenotype using gene counts of known AMR genes with and without gene counts of all other functionally annotated genes in the genomes (eggNOG gene families). Each of these approaches (further outlined below) was independently applied to the prediction of resistance to 23 different antibiotics for which relevant MICs were available.

### Naive prediction of AMR phenotype

Although RGI and other AMR gene finder tools do not claim to be able to infer AMR phenotype, the presence of an AMR gene is often used to designate whether a genome is susceptible or resistant [40, 6]. Therefore, the presence of an RGI-annotated AMR gene was used as an indicator of resistance to the antibiotic(s) to which the gene was labelled as resistant in the CARD database. Precision, recall and accuracy for both susceptible and resistant phenotypes were calculated for this naive model using a custom Python script (OG_RGI_analysis.py) as a baseline to compare the subsequent models.

### Logistic regression prediction of AMR phenotype

To evaluate the relationship between the AMR genotype and AMR phenotype, a logistic regression model was used for each antibiotic (Fig.1) with a split of 3:1 between training and test datasets respectively, using a custom python script (Logistic_regression_RGI.py). This model evaluated how the presence or absence of specific AMR genes was related to the AMR phenotype. Model precision, recall and accuracy for both susceptible and resistant phenotypes were calculated to evaluate the model efficacy and potential bias. The ratio of susceptible organisms to resistant organisms can help determine the likelihood of bias in the training data (Fig.S1).

**Figure 1.**
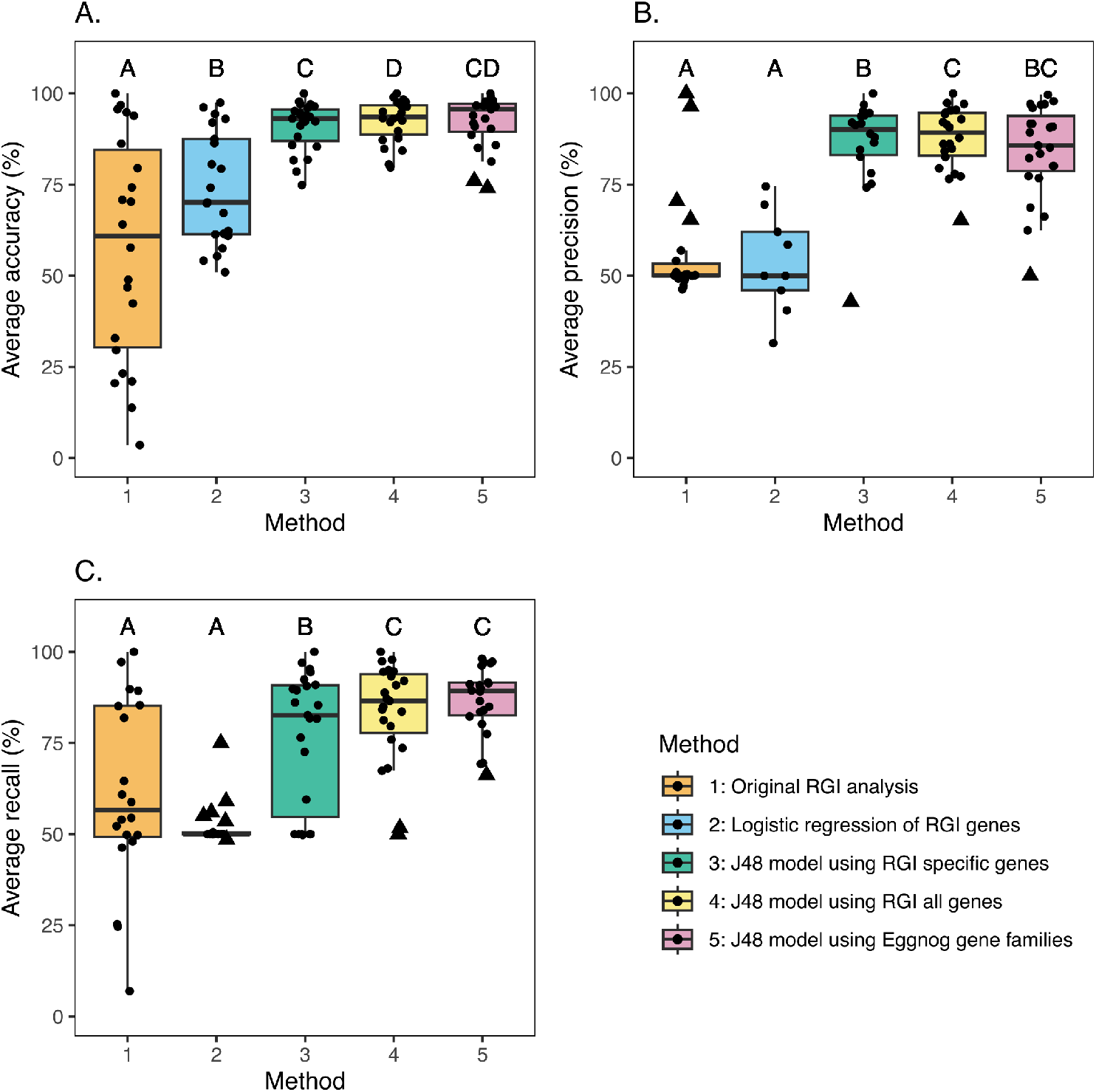
Model average accuracy (A), average precision (B) and average recall (C). The boxplots represent the following methods used in this study to predict AMR phenotype in the following order: Naive RGI analyses (orange), logistic regressions using the RGI data (blue), J48 decision trees using RGI genes specific to the antibiotic (green), J48 decision trees using all RGI genes regardless of the antibiotic model (yellow), J48 decision trees using Eggnog gene families (pink). The statistical significances are the result of a pairwise Wilcoxon signed-rank test adjusted for multiple testing using the Benjamini-Hochberg method (q*<*0.05). No significant difference between distributions is indicated by a shared letter above their respective boxplot (see Supplementary Table 11 for more details). Outliers are represented with a triangle-shaped point.

### Decision tree prediction of AMR phenotype using only AMR genes

To understand how specific AMR genes may drive the relationship between the AMR phenotype and AMR genotype, 4 machine-learning approaches were used. A custom Python script was used to convert the RGI gene counts into an Attribute-Relation File Format (ARFF) file (RGI_specific_analysis.py) and using the csv2arff tool found at https://github.com/LucyDillon/CSV_2_arff. The J48 decision tree models were built as implemented in the WEKA machine learning platform (version 3.8.5) [12]. The J48 model is written in Java and is an adaptation of the landmark C4.5 algorithm. In this analysis, the model takes into account the number of copies or absence of an AMR gene in relation to the AMR phenotype. J48 decision trees are used to classify each ‘instance’, or genome, based on the provided labels (AMR gene count). The model evaluates the data overall and then splits the genomes based on their labels (one label-based decision for each split). Next, it repeats this process on the subsets of genomes until the model has reached a preset limit based on either model parameters, such as the minimum number of genomes per split, or a consensus split of the correct categorical variable, in this case, AMR phenotype (further details below).

This analysis was then repeated using the Random Forest, Support Vector Machine (SVM - WEKA package: libsvm 3.25) and Logistic Model Trees (LMT) models in WEKA to compare the efficacy of each machine learning approach (Supplementary Table 2).

Models for 23 different antibiotics were selected with respect to various data constraints. Each model is trained specific to a single antibiotic and the genomes present in the model must have corresponding MIC values. For a model to be able to learn from the data and thus predict the correct AMR phenotype, the models had to have both susceptible and resistant organisms (Supplementary Table 3). The proportion of organisms with a susceptible to resistant MIC value can be seen in Fig.S1.

The J48 model was chosen for further analysis due to the interpretability of its decisions. Hence, providing the biological reasoning behind the predictions it made. The output of the J48 model is a human-readable tree of the decisions to partition the genomes (as resistant or susceptible) (Fig.S3-S5). The default parameters were used for the WEKA J48 model, however, the parameters were first evaluated by a matrix comparing M (Minimum number of instances per leaf) and C values (Confidence value: the lower value indicates more pruning) (Supplementary Table 4). There was no difference in eight of the antibiotic models using the different parameters and the rest of the models had minor differences. The most accurate C value could be found by using 0.25 or 0.5 for 15 out of 23 antibiotic models. The C value of 0.25 was selected as this level of tree pruning is recommended to not overfit the model or prune the tree too much and miss important information. The M value of 0.2 was selected as this is the default of the model and the other M values had very similar accuracy. The model accuracy was evaluated by 10-fold cross-validation. The individual fold results allowed the standard error of the models to be calculated (Supplementary Table 3).

To evaluate what factors may impact the models or improve model accuracy, the composition of AMR genes used to train the models was analysed. The models were originally trained using specific AMR genes for the antibiotic the model represented. For example, Ampicillin-specific AMR genes to train the Ampicillin model. The antibiotic target is defined in the CARD database in which the genes are annotated to correspond to specific antibiotics. The models were then trained with all AMR genes present in the genomes regardless of which antibiotic model they were training (Supplementary Table 3, Supplementary Table 5, Fig.S2). A custom Python script used to make the .arff files for this analysis (RGI all analysis.py).

### Investigating the role of taxonomy on decision tree model accuracy

To investigate the role of taxonomy on model accuracy, for each antibiotic model, one genus was excluded from the training data. The excluded genus was then used to test the model. This included each genus available for each antibiotic model (see Supplementary Table 6 for details). This way we can evaluate how the models may perform on a species that was not in the training set. We used a custom python script to develop the .arff files (taxa test train files.py).

To process CSV files into the format required for weka (.arff) we created a simple tool to translate a .csv file into a .arff file. This code is freely available at https://github.com/LucyDillon/CSV_2_arff.

### Analysis of accessory gene involvement in AMR phenotype

To investigate the role of accessory genes in AMR phenotype, the genomes from BV-BRC were analysed using Prodigal v2.6.3 [21], Diamond v0.9.24.125 [8], and eggNOG-mapper version 2.1.6 to predict gene families [10]. All tools were used with default parameters. The least specific level of the eggNOG gene family (i.e. COG or NOG) was taken to get the most general result so that the gene families could be compared across different taxa. The number of genes present in a gene family, including their absence, was compared with the genome AMR phenotype using the same J48 model and parameters in Weka, using a custom python script to make the .arff files (Eggnog_analysis.py). A 10-fold cross-validation was used to evaluate the model’s accuracy in predicting AMR phenotype, from which the standard error was calculated.

We mapped the RGI AMR genes onto the eggNOG decision tree models by analysing the CARD database with or eggNOG-mapper. The eggNOG gene families reported were matched to the AMR genes and were then labelled as having a known AMR gene function in the models. Finally, “pathways to resistance” were identified in all of the resulting decision tree models by identifying all possible paths through the resulting trees that lead to a “resistance” outcome, using Apply_Decision_Tree available at https://github.com/ChrisCreevey/apply_decision_tree. All gene families traversed to reach each resistance outcome on the decision trees were considered important to that resistance path (regardless if it needed to be present or absent) and included in subsequent analyses of different paths to resistance.

### Investigating protein-protein interactions

Protein-protein interactions of the gene families within individual decision trees were investigated using the STRING protein-protein interaction database (version 11.5) [39]. One protein sequence from each gene family was selected (the first sequence in the fasta file downloaded from eggNOG for each gene family) to represent that gene family (723 unique gene families in total) using the “Protein families “COGs”” and “multiple sequences” options for each individual antibiotic model in STRING. This analysis was used to highlight if gene families within the same model or pathway to resistance were predicted to interact and therefore, may have a role together in AMR phenotype.

To find associations with the gene families across multiple models, STRING and Cytoscape (version 3.9.1) [38] were used to analyse the data using the same options as above, but including all gene families from all antibiotic models. To reduce the network to directly link to the decision trees, edges in the network were only retained if both gene families were present in the STRING protein-protein interactions (and therefore predicted to interact) and the pair was also present in at least one decision tree model. Details of this analysis can be found in the Cytoscape_analysis.md file.

The predicted protein-protein interaction network of each path to resistance was also produced using STRING, but including only those gene families predicted for each individual path (allowing connections based on low confidence to provide further evidence that these putative connections which may not be well documented in the database may have a role in AMR phenotype). To investigate if pathways to AMR phenotype within the decision trees are taxonomically related, we traversed the decision trees to investigate which route each genome took for each model. This was performed using Apply_Decision_Tree using the same input genome .arff and dot files. DOT is a graph description language to visualise information, such as decision trees.

For all models, accuracy is defined by the sum of the true positives and true negatives divided by the sum of the total number of genomes (instances). Precision and recall of the models are calculated for both susceptible organisms and resistant organisms separately. This highlights whether a model is better at predicting one phenotype over another.

## RESULTS

### Machine Learning approaches accurately predict AMR Phenotype from AMR Genotype

Within this study, we analysed several techniques for predicting AMR phenotype from genomic data, including logistic regression of AMR genes, J48 decision tree models, Random Forest, Support Vector Machine (SVM), and Logistic Model Trees (LMT).

Even though AMR gene finder tools are designed to identify the presence of AMR genes in genomic data, their results are frequently used to directly infer AMR phenotype in literature [40, 6, 17, 43]. We examined the accuracy of predicting AMR phenotype solely based on the presence/absence of AMR genes for 23 antibiotics and 16,950 genomes, from organisms with laboratory-derived MIC data. This naive model assumed an antibiotic-resistant phenotype when an AMR gene which targeted a particular antibiotic as defined in the CARD database was found in a genome.

The average prediction accuracy of this model (as defined by the number of genomes correctly predicted to be susceptible or resistant to an antibiotic divided by the total number of genomes tested) was 57.6% and ranged from 3.5% (Clindamycin) to 100% (Moxifloxacin) (Fig.1). Clindamycin had quite a poor ratio of susceptible to resistant genomes (273:10) in comparison to moxifloxacin which was better proportioned to make a more accurate model (4:10). The precision and recall were calculated using the confusion matrix (Supplementary Table 5, Supplementary Table 8). The average prediction precision was 56.2% and ranged from 46.3% (Fosfomycin) to 100.0% (Moxifloxacin) (Supplementary Table 8, Supplementary Table 6). The average prediction recall for all 23 antibiotics was 61.2% and ranged from 24.6% (Ertapenem) to 100.0% (Moxifloxacin). Logistic regression models of the RGI genes had an average accuracy was 73.9% and ranged from 50.96% (Erythromycin) to 97.44% (Amoxicillin) (Fig.S2), however, >50% of the models only predicted one phenotype, resulting in an average recall of 52.3% (ranging from 48.5% (Doripenem) to 75.0% (Amoxicillin)) and the average precision of 53.6% (ranging 31.5% (Doripenem) to 74.5% (Erythromycin)) (Supplementary Table 3, Supplementary Table 8).

As logistic regression did not result in good precision or recall for most models, we applied a decision tree approach (using the WEKA J48 model). The resulting decision tree models were highly accurate in predicting the correct AMR phenotype, when using 10-fold cross-validation the average accuracy was 91.1% and ranged from 74.85% (Tigecycline) to 100% (Moxifloxacin)(Fig.1, Supplementary Table 3, Supplementary Table 10). The average recall of the RGI-specific decision tree models was 76.8% (ranging from 50.0% for Amoxicillin, Aztreonam, Clindamycin, Colistin, Fosfomycin, and Nitrofurantoin to 100.0% for Moxifloxacin). The average precision was 86.2% (ranging from 43.0% Colistin to 100.0% Moxifloxacin). Furthermore, traversal of the resulting decision trees indicated different genomic routes to resistance and susceptibility (see (Fig.2), highlighting the importance of both the presence and absence of multiple genes to predicting AMR phenotype from genomic data).

**Figure 2.**
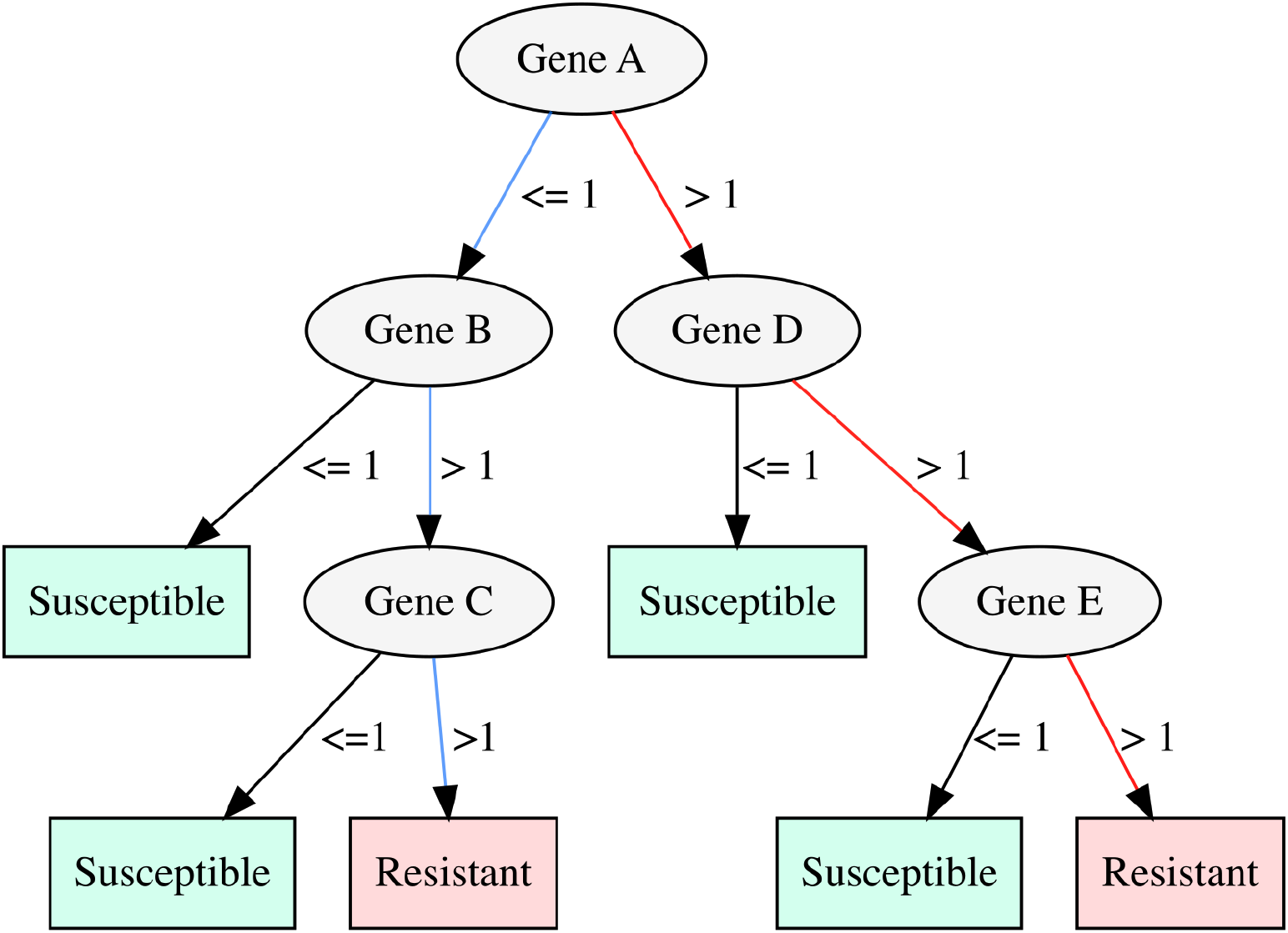
An example of a decision tree with two routes to resistance, indicated by the red and blue lines. For example, in the red pathway, if more than one copy of Gene A, D and E are present the genome will be resistant but if one of those genes is not present (i.e. Gene E) the organism will be susceptible.

The J48 model’s average accuracy of 91.0% was comparable to Random Forests 92.0%, SVMs 86.3% and LMT 92.2% (Supplementary Table 2, Fig.S7) and had the advantage over the other models of allowing biological interpretation of the genes driving the AMR phenotype/genotype relationship. For this reason, we focussed further analysis on the decision tree models.

### Model accuracy is not reliant on specific taxonomy

To investigate the ability of the decision-tree models to predict AMR phenotype for groups of organisms that were not included in the training data, for each antibiotic, we generated multiple sub-datasets where for each we excluded all genomes (and MIC data) from a selected genus from the training data and re-generated the model. The excluded genus and associated MIC data were then used to test the accuracy of the regenerated model for predicting AMR phenotype across taxonomic groups. The AMR phenotypes were predicted accurately given that both phenotypes were distributed evenly in the training data. Despite that the genomes are dominated by the Pseudomonadota phylum, the models were able to predict 100% of *Streptococcus* phenotype (Bacillota phylum). However, in the Ampicillin model, *Salmonella* phenotypes were not predicted well (34%). This may be due to a severe imbalance in phenotypes in the training data, meaning it incorrectly predicted *Salmonella* made from a biased model. Nevertheless, the average accuracy was 80.3% (ranging from 0% to 100 %) when trained on a different genus (Supplementary Table 6). *Klebsiella* had the most genomes for each antibiotic model and had relatively good accuracy (average accuracy 84.4%). However, in some cases, the genus with the second-largest number of genomes occasionally performed poorly for example, for Ampicillin *Salmonella* scored 34.3% but for Ciprofloxacin scored 97.7%. This may be due to some genera being more genetically similar than others, such as *Kelbsiella* and *Escherichia* compared to *Nessieria*. To further investigate *Salmonella*, the results of the taxonomic analysis were compared to the tree traversals showing which different routes to AMR phenotype, this shed light on why *Salmonella* had high accuracy for Ceftriaxone, Ciprofloxacin, and Gentamicin but not Ampicillin. The tree traversal showed that in the models for Ceftriaxone, Ciprofloxacin, and Gentamicin the genera were very diverse and are not dominated by a singular genus. Yet, in the model for Ampicillin the tree traversal showed that the majority of the genomes were from *Salmonella* and every pathway contained *Salmonella* genomes. Therefore, when excluded from the training data the pathways for *Salmonella* will have also been excluded when building the model, hence when testing the accuracy is low.

### ML models identify putative additional antibiotic targets of AMR genes

To investigate the role of AMR genes in antibiotic resistance to which they are not indicated in the CARD database, we generated decision trees which included all AMR genes regardless of the antibiotic target listed in the CARD database. This resulted in 17 antibiotic models improving in accuracy and overall significantly better compared to the models using only the AMR genes specific to the antibiotic which is listed in CARD (Wilcoxon signed rank test (q = 8.27E-04) (Supplementary Table 11)). The average accuracy was 92.5% (ranging from 79.7% (Tigecycline) to 100.0% (Moxifloxacin)) and the average recall and precision were 83.5% (ranging from 50.0% (Fosfomycin) to 100.0% (Moxifloxacin)) and 87.5% (ranging from 65.0% (Nitrofurantoin) to 100.0% (Moxifloxacin)), respectively (Supplementary Table 8). A significant increase in average recall and precision was also observed (recall q = 4.39E-04 and precision q = 0.04) (Supplementary Table 11). This suggests that AMR genes may have additional antibiotic targets not annotated in the databases. One example of this can be seen in the Gentamicin RGI-all model, which shows the presence of > 1 TEM-185 genes confer resistance to Gentamicin (Fig.S4). This particular gene is not labelled to confer resistance to aminoglycoside antibiotics in the CARD database. We investigated if the models were better at predicting one phenotype than the other, this can be inferred from Supplementary Tables 5, 10, and 12 confusion matrices.

### Accessory genes have a key role in AMR phenotype

To see if this observation extended to non-classic AMR genes, decision trees were generated for the 23 antibiotics using eggNOG gene family functional profiles generated for all 16,950 genomes. The average accuracy for these models was 92.2% (ranging from 74.0% (Tigecycline) - 100.0% (Moxifloxacin)). In the comparison of the eggNOG models to the RGI models, the mean value was the highest for RGI all analysis (92.5%) (Supplementary Table 3, Supplementary Table 12). The difference between the RGI decision tree models and eggNOG gene families was not significant overall (RGI specific genes vs Eggnog q = 3.66E-01, RGI all genes models vs Eggnog q = 2.49E-01) (Supplementary Table 11). Overall, the eggNOG models were not significantly worse than the AMR gene-based decision trees, this highlights that the decision trees are able to extract key genes involved in AMR phenotype.

The average precision of the eggNOG-based decision tree was 84.3% (ranging from 50.0% (Fosfomycin) to 100.0% (Moxifloxacin)) (Supplementary Table 8). This was significantly better than the logistic regression of RGI genes. This suggests that the models based on the eggNOG genes are less biased than the logistic regression and the RGI-specific analysis.

The average recall of the eggNOG-based decision trees was 86.6% (ranging from 66.0% (Colistin)-100.0% (Moxifloxacin)) (Supplementary Table 8). This was significantly better than the naive RGI, the logistic regression and RGI-specific decision tree models.

We can gain further biological insight by using these accessory genes which may be involved in resistance pathways. This could provide novel information about pathways to resistance to particular antibiotics. Using the eggNOG decision trees we found an additional 675 gene families across all 23 models which are not in the RGI database but are linked to the AMR phenotype.

### Decision trees show biological pathways to resistance

The use of decision trees allowed biological interpretation of (428 susceptible routes and 528 resistant routes in eggNOG models) paths to resistance and susceptibility (Fig.3 and Fig.S3-S5, Supplementary Table 13). The models showed the importance of the absence or number of copies of genes which could influence the AMR phenotype of an organism. The co-occurrence of genes was another important factor when determining the AMR phenotype. This highlights key genes involved in AMR phenotype that may not be classic AMR genes (Fig.S7). RGI genes were matched to the gene families in the decision tree models, we can see that the majority of models contained RGI gene families. In the eggNOG-based Amikacin model, COG0050 is matched to a multidrug-resistant gene (*Escherichia coli* acrA) but this is not involving aminoglycoside suggesting that this gene may have additional targets. The Tetracycline decision tree model using eggNOG gene families shows there are 6 routes to resistance. The decision trees have values for each phenotype in the tree so the number of genomes reaching that route can be distinguished, this way the main routes to resistance are revealed (in the case in which there are two numbers, the first is the total number of genomes and the second is the number of incorrectly classified genomes). The most common route to resistance involves COG0480 and COG0765. The gene family COG0480 is a key gene family involved in Tetracycline resistance (tet(44)), Figure 3A shows that COG0765 does not have to be present for an organism to be resistant.

**Figure 3.**
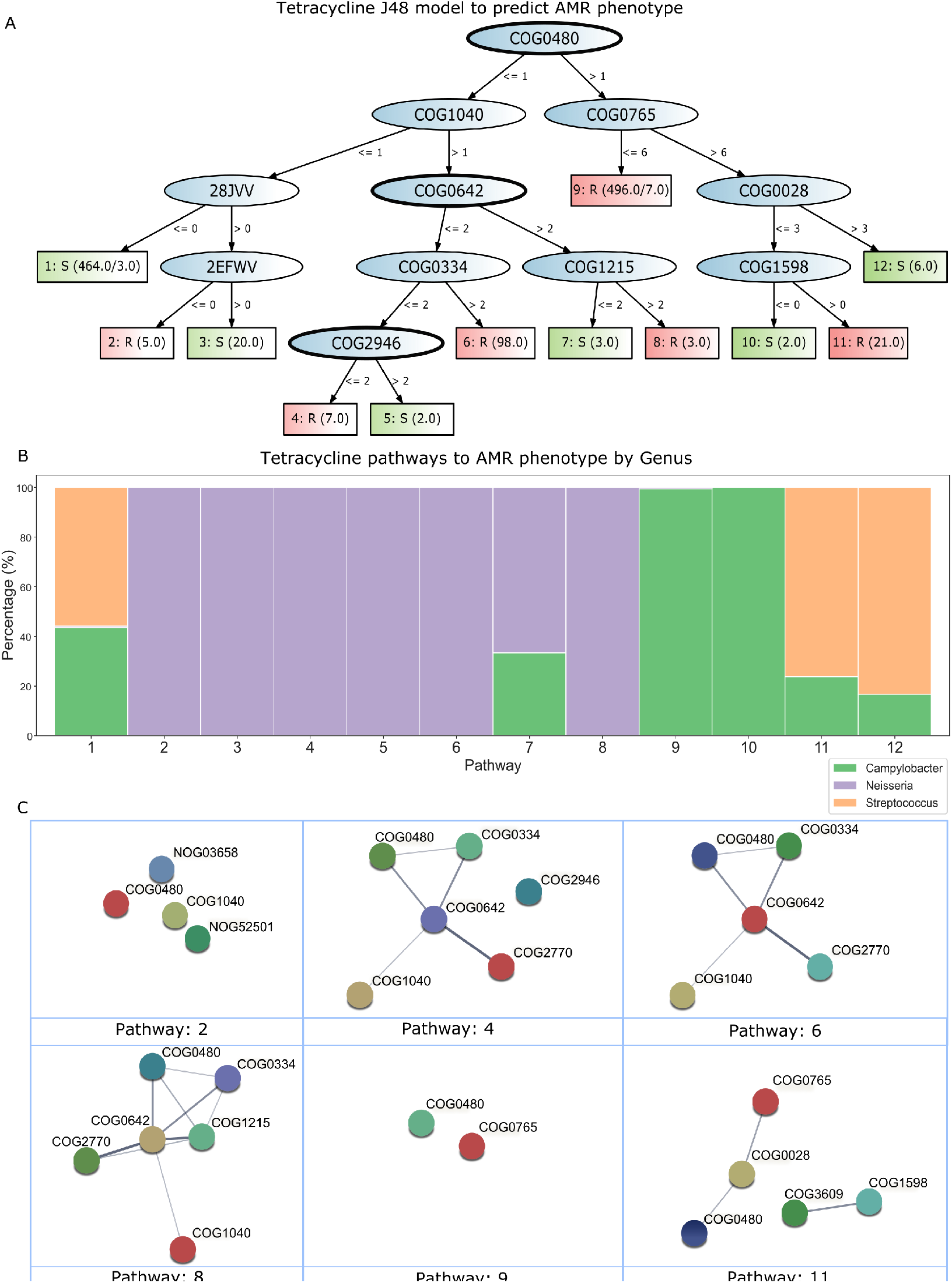
Predicting Tetracycline resistance using Eggnog gene families copy number or absence. **A**. A J48 decision tree model to predict Tetracycline AMR phenotype. RGI-associated gene families have been highlighted with a thick black outline. COG0480 relates to gene tet(44), COG0642 relates to gene adeS, and COG2946 relates to gene tetU. The decision trees have numbers in the phenotype boxes to represent the number of genomes. This may include two numbers in some cases, the first number indicates the total number of genomes and the second number are incorrectly classified genomes.* **B**. Stacked bar chart showing the routes to susceptibility and resistance for Tetracycline. This is a genus level analysis, the species, family, order, class, and phylum analysis can be found in Fig.S8. The pathway numbers relate to the numbers on the decision tree (Part A). Note: pathway 9 is not 100% *Campylobacter*, 0.4% are *Nesseria*. **C**. Protein-protein interactions between gene families for each pathway to resistance. The lines (edges) represent the protein-protein interactions from STRING and the thicker the line, the higher the confidence (see Supplementary Table 13 for details). See part A for details of each pathway (the pathway numbers correspond to the numbers on the phenotype boxes in part A).* Note*: COG 28JVV and 2EFWV are recognised as NOG03658 and NOG52501 in the STRING database, respectively.

The analysis to link eggNOG gene families and RGI AMR genes revealed that most decision tree models had several gene families linked to RGI genes. However, the RGI gene families did not dominate the trees suggesting that accessory genes have an important role in AMR phenotype. The eggNOG-based decision tree using the Tetracycline phenotypic data had three gene families (tet(44), tetU, and adeS) associated with known AMR genes, yet their presence did not always guarantee resistance to tetracycline (Fig.3A). Therefore, the presence or absence of certain accessory genes was relied on to confer the resistant phenotype.

STRING was used to identify putative protein-protein interactions between genes within each decision tree model. Of the 23 models, 18 models contained COGs which were predicted to have protein-protein interactions, for example, co-occurrence and co-expression. The Tetracycline decision tree using eggNOG gene families showed that 63.6% of the gene families had protein connections in STRING (Fig.S9). Each pathway to resistance was analysed in STRING to calculate a network of the protein-protein interactions of how likely they are to interact based on confidence values. Across all models, each path to resistance had an average of 1.2 connections ranging from 0 to 7.6 (Supplementary Table 13). After investigating the pathways to AMR phenotypes within the decision trees, we found that many of the routes are taxonomically dependent. This is shown in all versions of the decision tree models (using AMR and accessory genes). In Figure 3A we can see 12 distinct routes to AMR phenotype, pathways 4-6 are all classified as the *Neisseria* genus. Pathway 9 is classified as *Campylobacter* (99.4%) (Fig.3B). While this is not the case for every pathway in the trees (i.e. pathway 1 is very mixed) many of the branches in the trees could predict the taxonomy as precise as the species as well as the AMR phenotype. Additional antibiotic model pathways for species - phylum can be found in Fig.S8. Each pathway to resistance was investigated using the confidence values in STRING (including texmining, Neighborhood, Gene Fusion, Experiments, Co-occurrence, Databases, and Co-expression), we can see in Figure3C the majority of pathways have multiple connections. Details on all other pathways to resistance can be found in Supplementary Table 13.

### Resistance pathways to different antibiotics are distinct from each other

Using the decision trees we can work out which combinations of genes are involved in resistance (see example decision tree (Fig.2) for reference). Understanding which genes are key to resistance in particular antibiotics or shared across different antibiotics could help provide insight into novel approaches to combat AMR in the future. To investigate these key genes, we analysed every gene family present in every decision tree. The COG distribution across the different models was analysed and there were 723 unique COGs in total for all the models. Of these unique COGs, 48 were linked to RGI AMR genes (Supplementary Table 14). The distribution of different COG functional categories was varied across all models, suggesting that resistance to different antibiotics has distinct mechanisms (Fig.S6).

To find connections between all the antibiotic models we found protein-protein interactions in STRING for all gene families from all decision tree models. The initial STRING network had 450 nodes and 10,786 edges. This included all evidence types in STRING at a medium level of confidence (0.4). This was reduced to relate directly to the decision tree models by only including node pairs that had predicted protein-protein interactions in STRING and the same pair was also present in at least one decision tree.

The reduced network had 247 nodes and 1,300 edges, which showed clear clusters linking gene families to the specific antibiotic models (Fig.4). The network showed several gene families which connected to many different antibiotic gene clusters. This suggests these genes have an important role in AMR phenotype across multiple antibiotics. The most common gene family, COG2367 appeared in seven models. This is within the defence mechanisms group of eggNOG gene families, labelled as beta-lactamase. Interestingly, six of the models were built using the MICs of the beta-lactam antibiotics but COG2367 was also part of the Nitrofurantoin model which is distinct from the beta-lactam drug class. Within the Nitrofurantoin model, the COG2367 can have 4 copies of the gene family and still be susceptible (which according to the labels accounts for the majority of genomes on this pathway), if over 4 copies were present then there is a chance that the organism can be resistance. In the gene network, we can also see how different antibiotics have distinct genes which are only associated with that particular antibiotic (Fig.4, Fig.S10). This suggests that antibiotics have unique pathways to resistance. The network was clustered by antibiotic drug class, this showed that several drug classes were highly connected, including carbapenems and aminoglycosides (Fig.S11).

**Figure 4.**
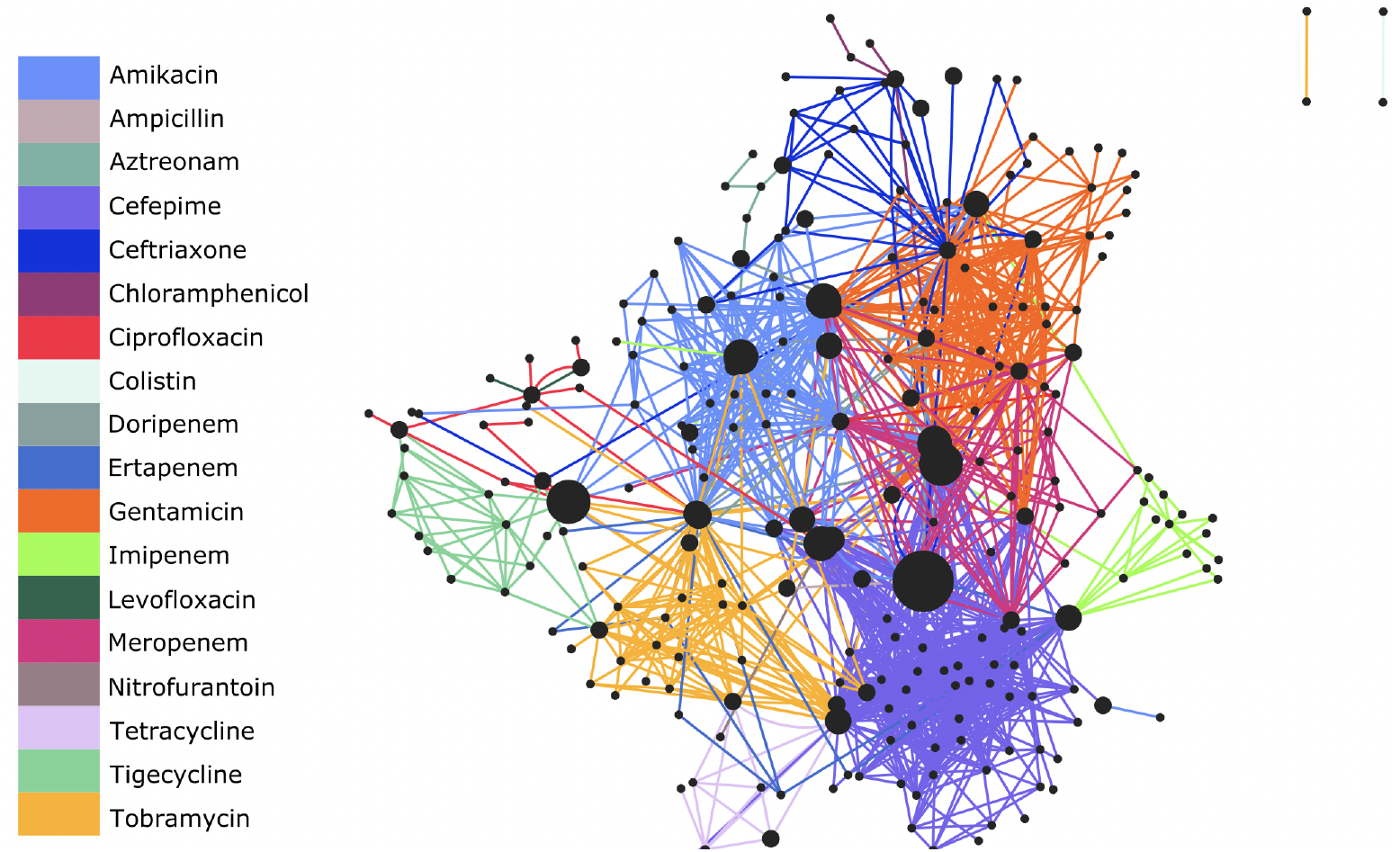
Gene network of all unique COGs across all antibiotic Eggnog models. Nodes are the different COGs, edges are protein-protein interactions between COGs. The edges are coloured regarding the antibiotic model that the COG pair is present in. Node size is proportional to the number of models the COG is present in.

## DISCUSSION

Within this study, we have shown that machine learning can vastly improve the prediction of AMR phenotype from genomic data. The consideration of accessory genes with AMR genes in these analyses provides valuable biological insights into the paths to AMR resistance and susceptibility.

The naive RGI model analysis shows that the presence of an AMR gene does not necessarily indicate the correct AMR phenotype. The average accuracy of RGI to predict AMR phenotype was 57.58%, which is comparable to a game of chance. The average precision was 56.2% and the average recall was 61.2%, which highlights key flaws in using the tool to predict AMR phenotype. While RGI is not designed to identify AMR phenotype but rather the AMR genotype, its results are often inferred as phenotypic resistance for genomes [40, 6, 17, 43] and metagenomes [22, 47].

Overall, while results from the logistic regression analysis was significantly better than those using RGI-only (Fig.1), it still underperformed which could suggest there is not enough data to make an accurate model, or there is not a strong enough link between the phenotype and the genes to be able to classify accurately using this approach. However, decision trees, show over 17% increase (statistically significant q= 6.03E-05, Supplementary Table 11) in accuracy compared to logistic regression suggesting that the poor performance of logistic regression may be due to the use of presence/absence information rather than the number of copies of genes, which the decision trees are capable of utilising.

The decision trees have shown that it is both the presence (including the number of copies) and absence of different gene families that are key in the accurate prediction of AMR phenotype. Biologically this makes sense as we know that genes perform their function most often as an ensemble with other genes. The decision trees show that even when a known AMR gene is present, it does not necessarily mean that the organism is resistant (Fig.S3-S5). Interestingly the decision tree models which included AMR genes not thought to be involved in resistance to the antibiotics (RGI all models) being examined showed an increase in accuracy compared to the models which were generated using only those AMR genes known to provide resistance to the specific antibiotic. This suggests that many of the AMR genes within the CARD database may be involved in providing resistance to a broader range of antibiotics than what is annotated in the database. However, AMR genes may need additional genes present to confer resistance to particular antibiotics which is not identified in any commonly used AMR gene finder tool.

The use of eggNOG gene families has shown the importance of accessory genes in the role of AMR phenotype. Accessory genes are generally ignored when determining the AMR phenotype of an organism when using computational techniques to predict AMR phenotype. Therefore, studies that rely on AMR gene finder tools to determine resistance could be misleading as the full picture is not described. All the eggNOG decision tree models are dominated by non-AMR genes (Fig.S5). The eggNOG models alongside the RGI models show that the presence of an AMR gene does not guarantee resistance to a particular antibiotic. Almost 30,000 gene families were used to train the eggNOG-based J48 models, in comparison, while 1,424 RGI AMR genes were used to train the RGI all-gene J48 models. The accuracy, precision and recall did not differ significantly between these models, this suggests that the J48 model is sufficient at extracting the most important factors involved in AMR phenotype. This is especially interesting as the eggNOG gene families are mostly not associated with AMR, unlike the RGI AMR genes which are defined in the CARD database to target particular antibiotic(s). Therefore, a lack of difference between the two datasets’ accuracy, precision and recall indicates the importance of accessory genes.

The precision and recall scores can be used to evaluate the model bias, using the values specific to the AMR phenotype. Therefore, we can evaluate if a model only predicts one phenotype well. The data shows that the average precision and recall for the logistic regression of the RGI genes were significantly worse than the majority of the decision tree models. The recall of the eggNOG models was not significantly different to the RGI all-gene analysis. The precision of the eggNOG models was not significantly different to both RGI decision trees. This suggests that the eggNOG models and the RGI models have a similar level of model bias despite the varying data input of the models (Supplementary Table 8).

Analysing the gene family networks demonstrates that different antibiotics have clusters of genes relating to that specific antibiotic, suggesting that different antibiotics have distinct genes involved in pathways to resistance. Yet, the models are still connected to various gene families from other clusters, suggesting there are key non-AMR annotated genes involved across many antibiotic resistance mechanisms (Fig.4). Creating gene networks based on specific antibiotic models in addition to the overall network, highlighted the links between specific gene families in mechanisms to resistance for particular antibiotics. The network analysis for each path to resistance showed that many pathways to resistance have multiple connections. This shows that genes could be dependent on other genes to help confer a resistance phenotype.

Identifying taxonomically dependent pathways to resistance within the decision trees highlights key genes to target for particular pathogens. Conversely, pathways to resistance with multiple taxa involved could suggest that the route is more transmissible resistance, which may provide opportunities for an easier target of resistance. These pathways also provide further biological insight into AMR mechanisms, many of which are understudied. Pathways dominated by one particular species but with other species present in small numbers could be indicators of horizontal gene transfer between species to confer resistance. Predicting the taxonomy in addition to the AMR phenotype will provide an additional function of the tool that these models are helping to create. More data and improvements would be needed to provide accurate predictions for greater taxonomic diversity. AMR phenotype cannot be clearly explained by the use of AMR gene finder tools alone. AMR gene finder tools may still provide a reasonable estimate for the AMR genotype of samples, yet to define specific AMR phenotypes a more detailed approach is required.

The J48 models provide more biological insight than other machine learning techniques tested such as random forest, SVM, and LMT. For example, the Tetracycline model generated from eggNOG gene families identified 6 genotypic routes to phenotypic resistance. The main pathway to resistance involves the presence and absence of two gene families, COG0480 and COG0765 (Fig.3). COG0480 is linked to an RGI gene (tet(44)), and COG0765 is in COG category P and involved in amino acid transport. Further work needs to be done in this area to investigate the role of the presence and absence of accessory genes and AMR phenotype.

## Summary

We have shown that AMR phenotype can be accurately predicted using interpretable machine learning models such as decision trees that utilise both known AMR and accessory genes (eggNOG gene families) for multiple taxonomies, across multiple antibiotics. The use of AMR gene finder tools has repeatedly been shown to have limitations in their ability to predict AMR phenotype based solely on the presence of AMR genes. The use of machine learning techniques in this study has shown the benefit of analysing different factors, such as gene counts and absence, as key factors together when predicting AMR phenotype. Equally, we have also highlighted that the role of accessory genes in AMR phenotype is understudied in relation to AMR. Building models with the near-complete functional capacity of a genome showed accessory genes are fundamental to resistance. Finally, this study demonstrates the complexity of the AMR phenotype in relation to its genome but it has also highlighted that there are routes to resistance that are taxonomically dependent.

These machine learning approaches have the potential to transform laboratory-based diagnostics, providing a rapid and affordable alternative to culture-based techniques, estimating taxonomy in addition to AMR phenotype, and providing real-time monitoring of multi-drug resistant pathogens.

A call for data: If you would like to be involved in improving these models by contributing genomes with corresponding MIC (micro broth dilution) data please contact us at.

## Supporting information

All supplementary files, incase link in the paper does not work for readers.

## ACKNOWLEDGMENTS

We acknowledge funding from the Department for Economy Northern Ireland for PhD funding for L.D. C.J.C. wishes to acknowledge funding from the European Commission via Horizon 2020 (818368, MASTER and 101000213 HoloRuminant). N.J.D. wishes to acknowledge the Farncombe Digestive Health Disease Institute (McMaster University) and a grant from the Weston Family Microbiome Initiative. This work was undertaken on Kelvin2, an EPSRC-funded tier-2 High-Performance Computing facility at Queen’s University Belfast, UK.

## AUTHOR CONTRIBUTIONS

**LD** Carried out the data analysis, writing of the code on the AMR_ML_paper and CSV_2_arff GitHub.

**NJD** Advised on using ML and review of manuscript and methods.

**CJC** Tree traversal code and direction of scientific discovery and reporting.

All authors contributed to the scientific direction and writing of the manuscript.

## DATA AVAILABILITY

All supplementary data and additional files can be found here: https://osf.io/cj4bq/?view_only=c0ee87b7609543b688953089be4c376f. For specific files please.

## CODE AVAILABILITY

All code used in this study can be found at the following links: https://github.com/LucyDillon/AMR_ML_paper/tree/main https://github.com/LucyDillon/CSV_2_arff https://github.com/ChrisCreevey/apply_decision_tree/tree/master

