## Supplementary material for "Accessory genes define species-specific pathways to antibiotic resistance": All supplementary files, incase link in the paper does not work for readers.: Supplementary figure and table legends.docx

Supplementary Figure 1: Antibiotic resistance phenotype ratio. Each antibiotic model is shown and the ratio of the susceptible strains to resistant strains.

Supplementary Figure 2: A schematic of methods of build decision trees using RGI AMR genes.

Supplementary Figure 3: RGI specific decision trees. Parts A-W represent each antibiotic model.

Supplementary Figure 4: RGI all-gene decision trees. Parts A-W represent each antibiotic model.

Supplementary Figure 5: Eggnog gene family decision trees. Parts A-W represent each antibiotic model.

Supplementary Figure 6: Distribution of COG categories across each model.

Supplementary Figure 7: Model accuracy for different ML techniques. Including J48, LMT, RF, and SVM (left to right).

Supplementary Figure 8: Tree traversal pathways for each antibiotic at the following taxonomic levels: Species, Genus, Family, Order, Class, Phylum. The pathways can be matched to the trees (Supplementary Figure 7) by following the routes down the tree left to right (starting at the top node- use the numbers in Figure 3A for clarity).

Supplementary Figure 9: Diagrams of STRING evidence based for each antibiotic model.

Supplementary Figure 10: Gene network of protein-protein interactions by individual antibiotics.

Supplementary Figure 11: Gene network of protein-protein interactions by drug class.

Supplementary Table 1: Number of genomes by genus and antibiotic.

Supplementary Table 2: Comparison of different Machine Learning techniques, with the addition of the original RGI analysis for reference (NOT A ML technique). The model accuracy is shown for each individual antibiotic model (%).

Supplementary Table 3: Model accuracy and standard error values.

Supplementary Table 4: J48 model accuracy using different parameters.

Supplementary Table 5: J48 model confusion matrix using RGI data using all AMR genes regardless of the model.

Supplementary Table 6: Taxonomy analysis for each antibiotic model. The genus shown is the taxa which was tested on the model (excluded from the training set). The model accuracy is shown in percentage (%) and the number of strains is also shown.

Supplementary Table 7: Original RGI analysis confusion matrix.

Supplementary Table 8: The precision and recall values for each kind of model: logistic regression of RGI data, J48 model using RGI specific genes, J48 model using all RGI genes, and J49 model using Eggnog gene families. The precision and recall have been worked out for both resistant (res) and susceptible (sus) phenotypes to see if the model is better at predicting one than the other.

Supplementary Table 9: Confusion matrix of the logistic regression.

Supplementary Table 10: Confusion matrix of RGI specific J48 model

Supplementary Table 11: Wilcoxon signed rank tests of datasets.

Supplementary Table 12: Confusion matrix of J48 model based on Eggnog gene families.

Supplementary Table 13: Each pathway to resistant phenotype for each model. Average degree is the number of nodes connected by confidence score in the STRING database (low confidence). Number of nodes is the number of gene families involved in each pathway.

Supplementary Table 14: Each gene family by model and if linked to an RGI AMR gene family (1 represents it is linked, 0 represents it is not linked).
