## Supplementary figures and images for "Accessory genes define species-specific pathways to antibiotic resistance"

### Part_A_Amikacin_RGI_all.png

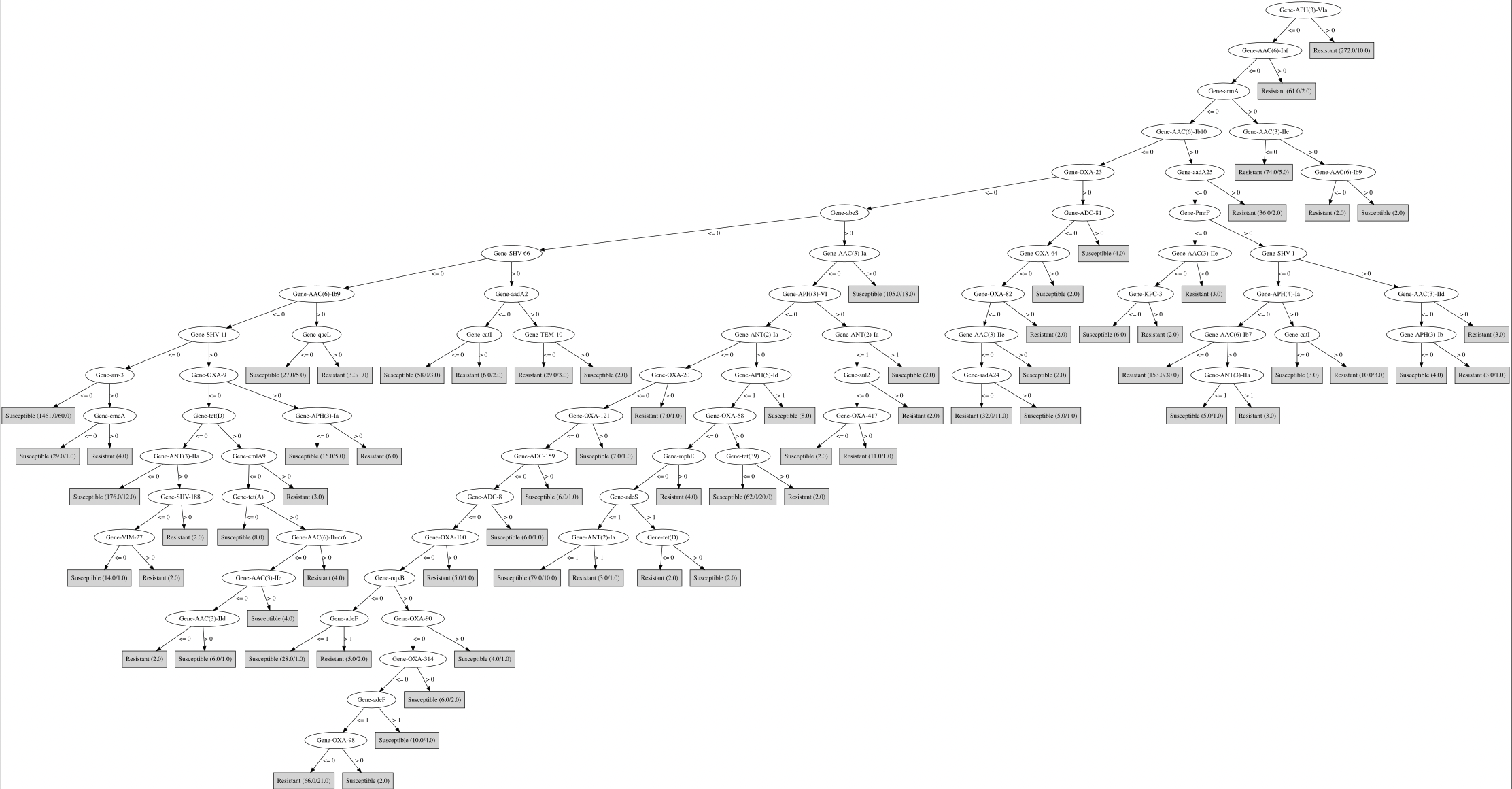

### Part_B_Amoxicillin_RGI_all.png

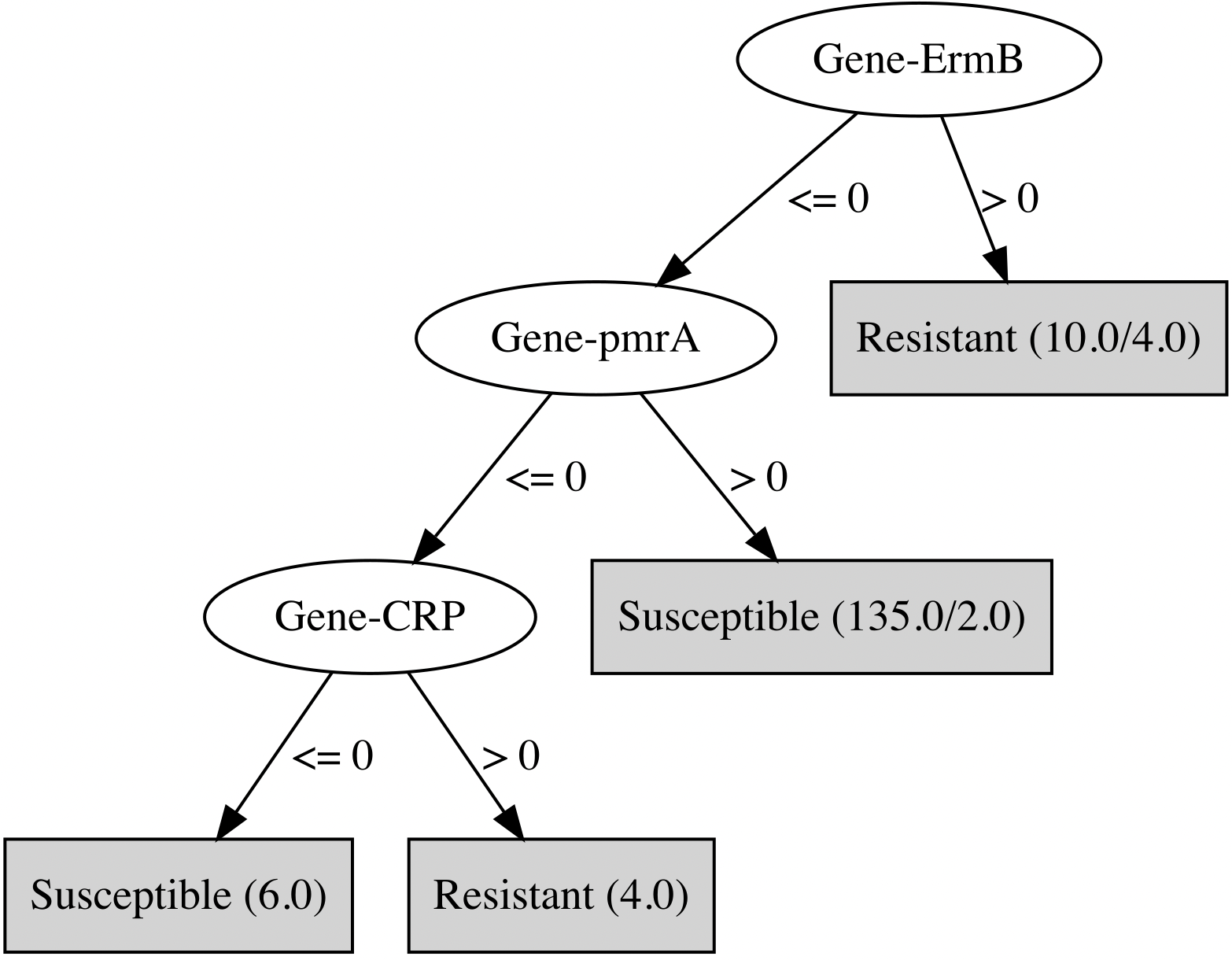

### Part_C_Ampicillin_RGI_all.png

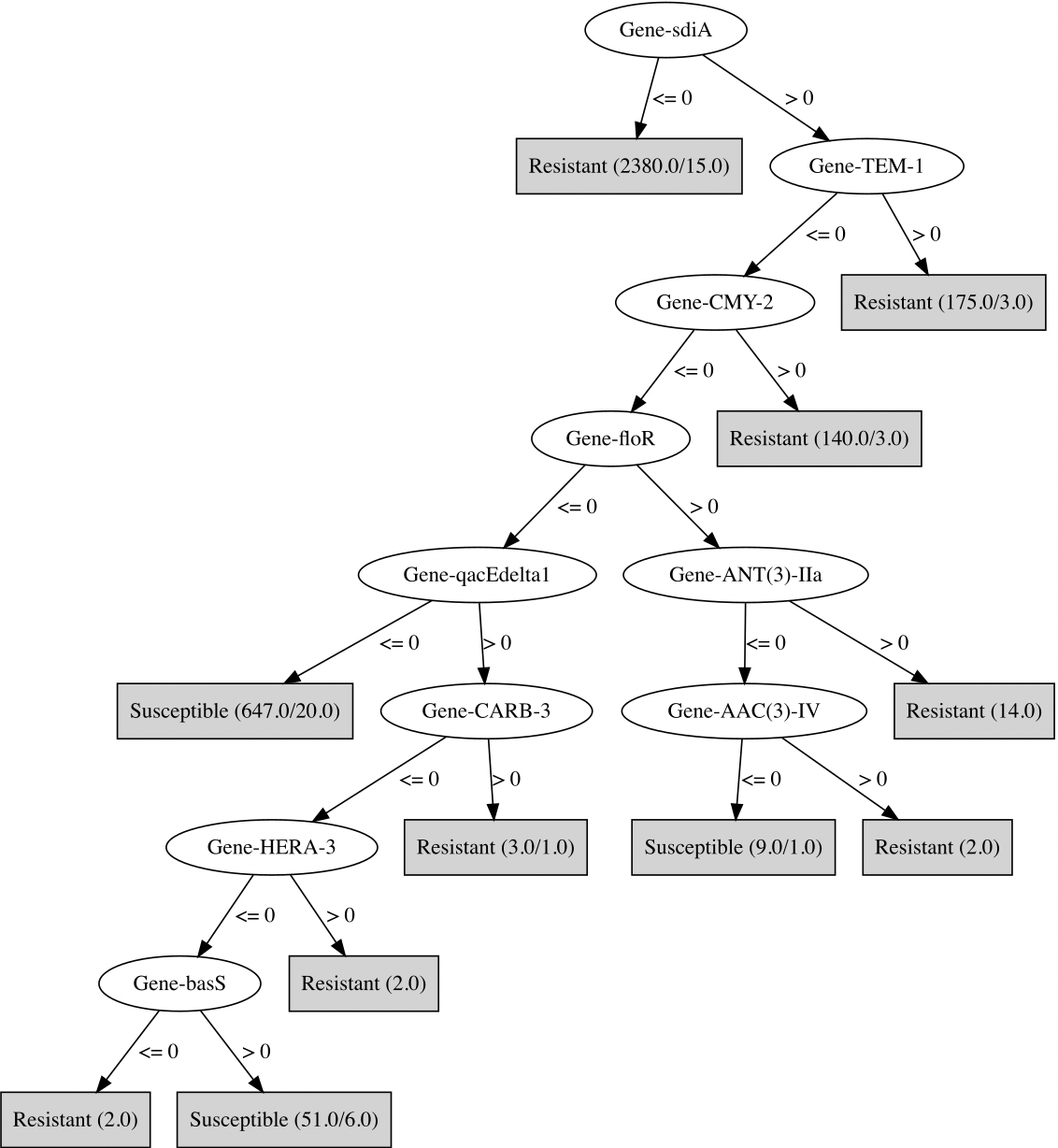

### Part_D_Aztreonam_RGI_all.png

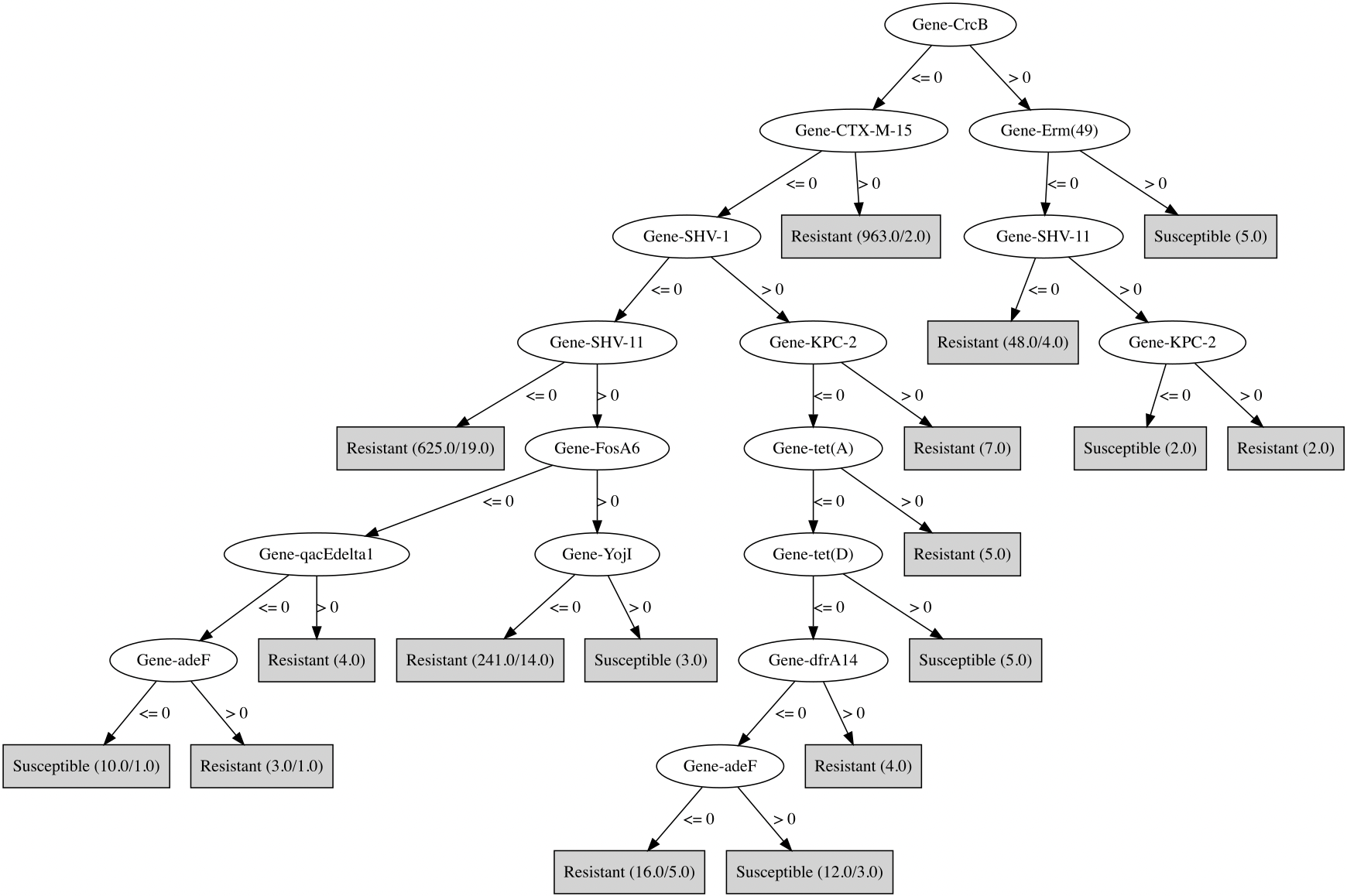

### Part_E_Cefepime_RGI_all.png

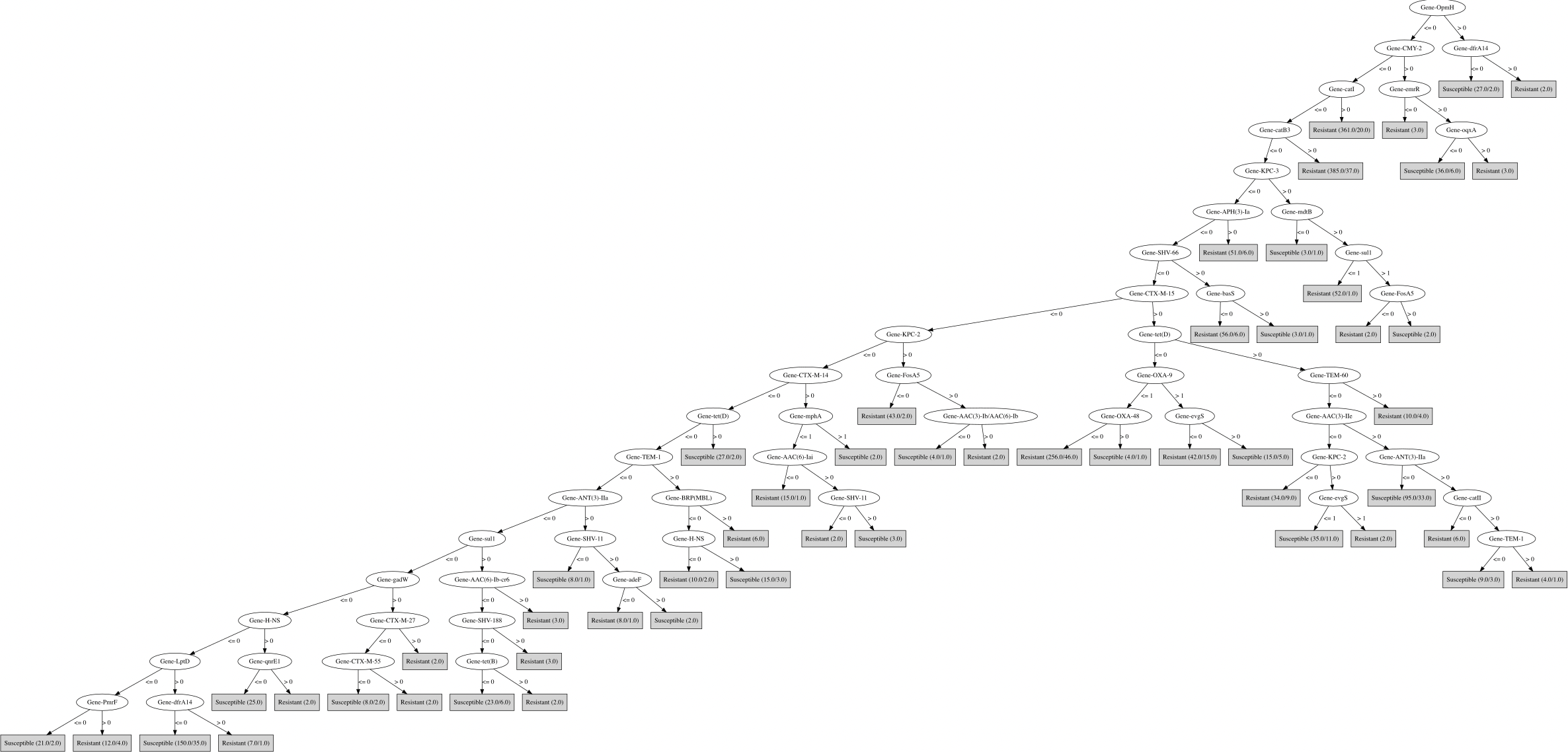

### Part_F_Ceftriaxone_RGI_all.png

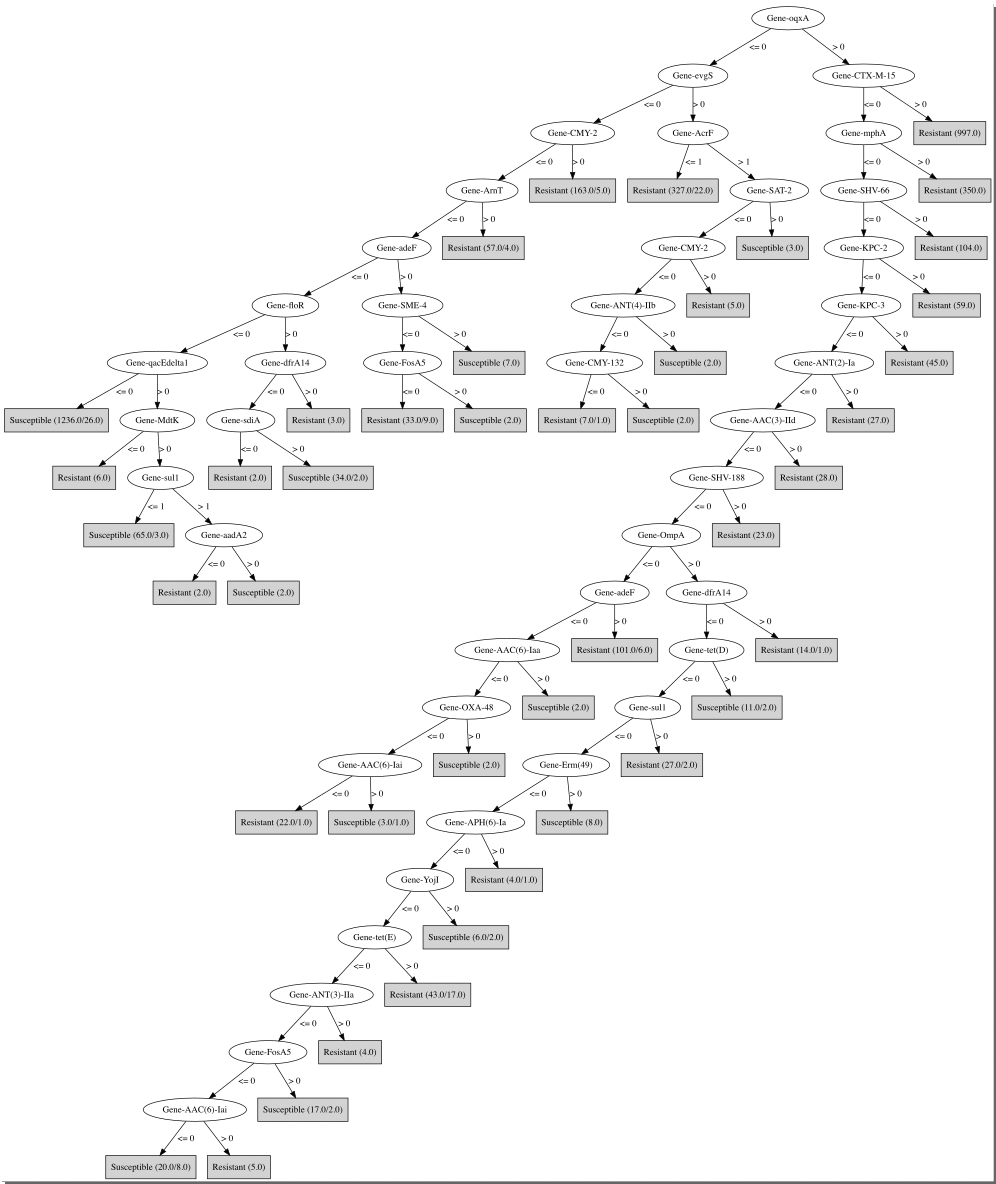

### Part_G_Chloramphenicol_RGI_all.png

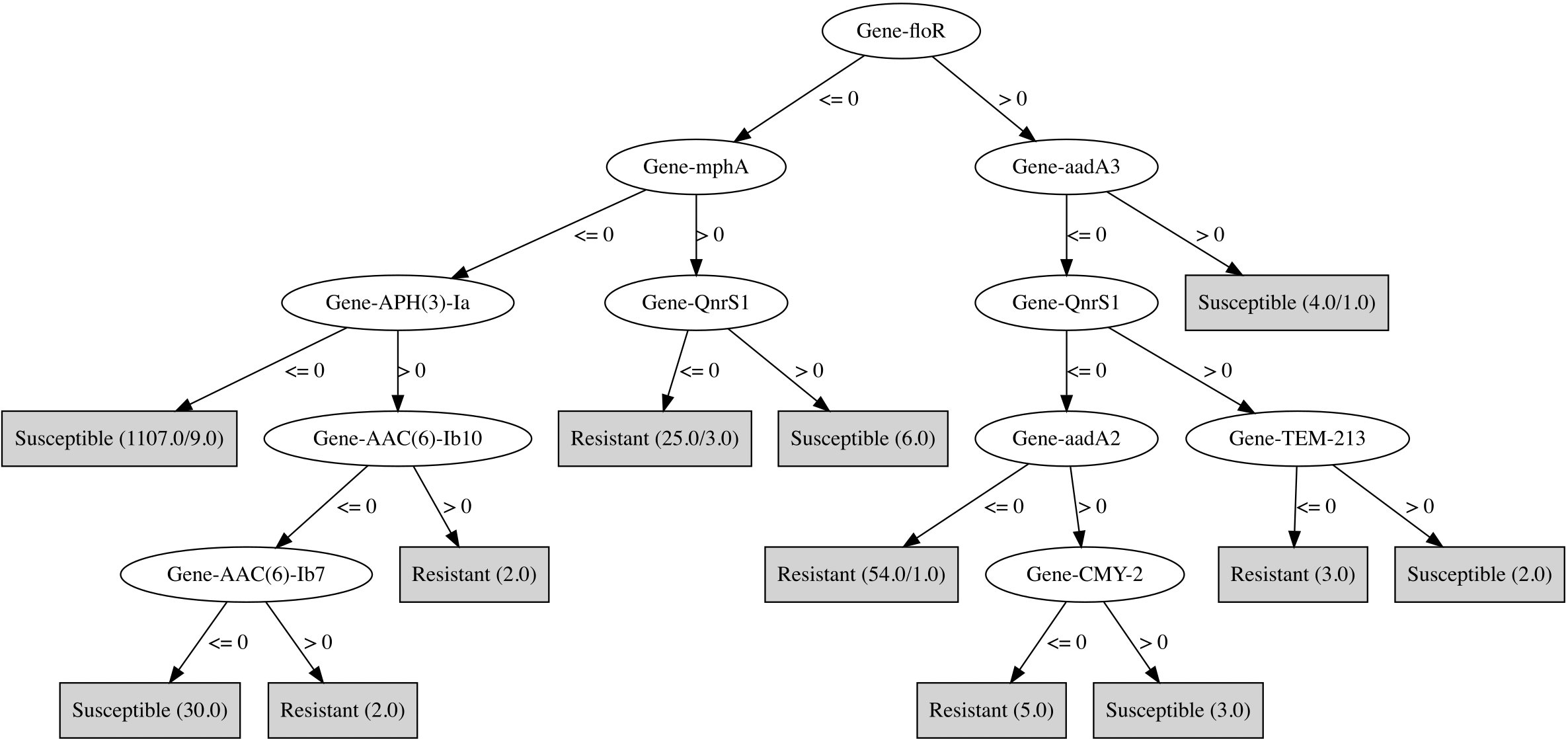

### Part_H_Ciprofloxacin_RGI_all.png

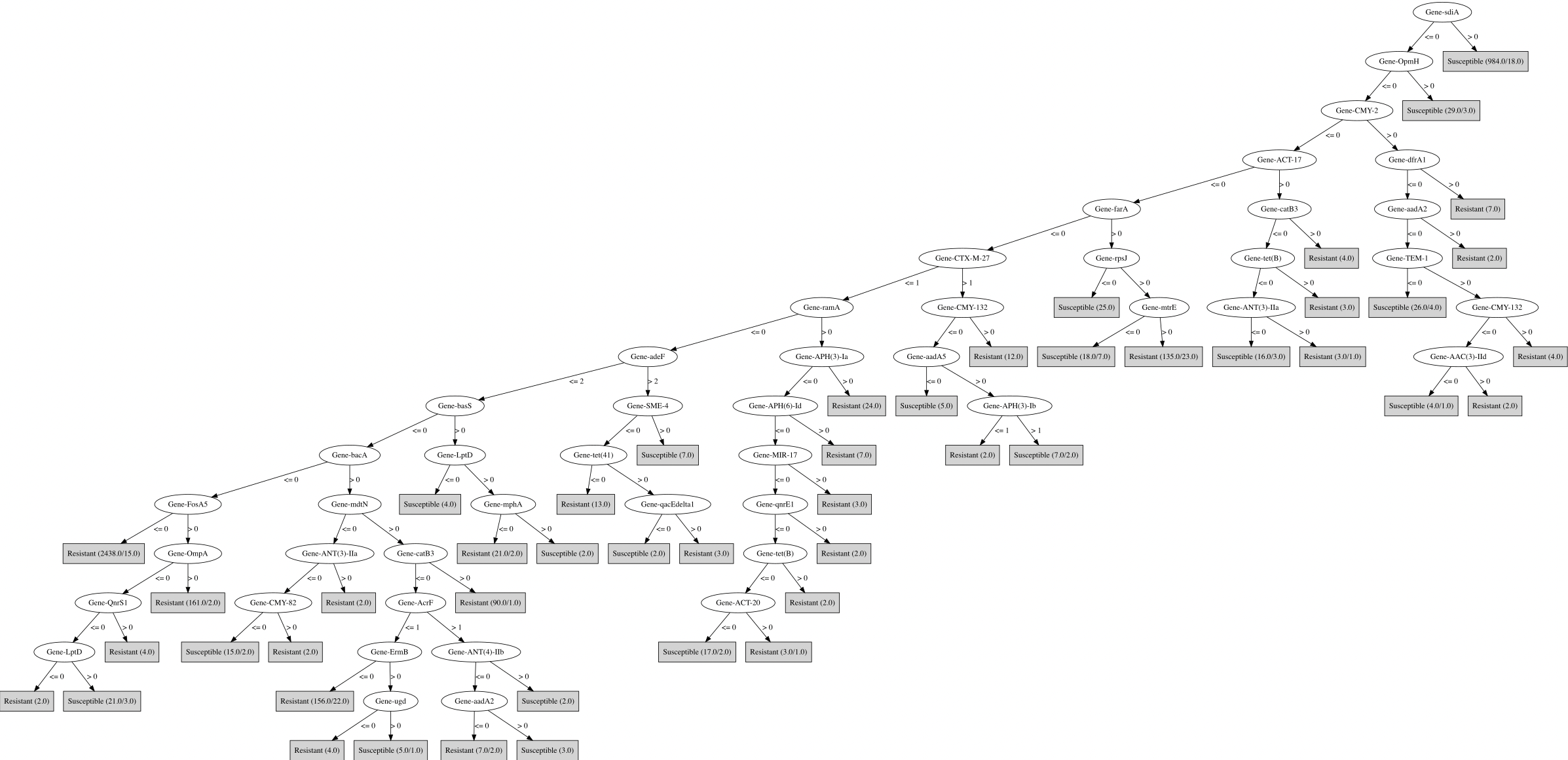

### Part_I_Clindamycin_RGI_all.png

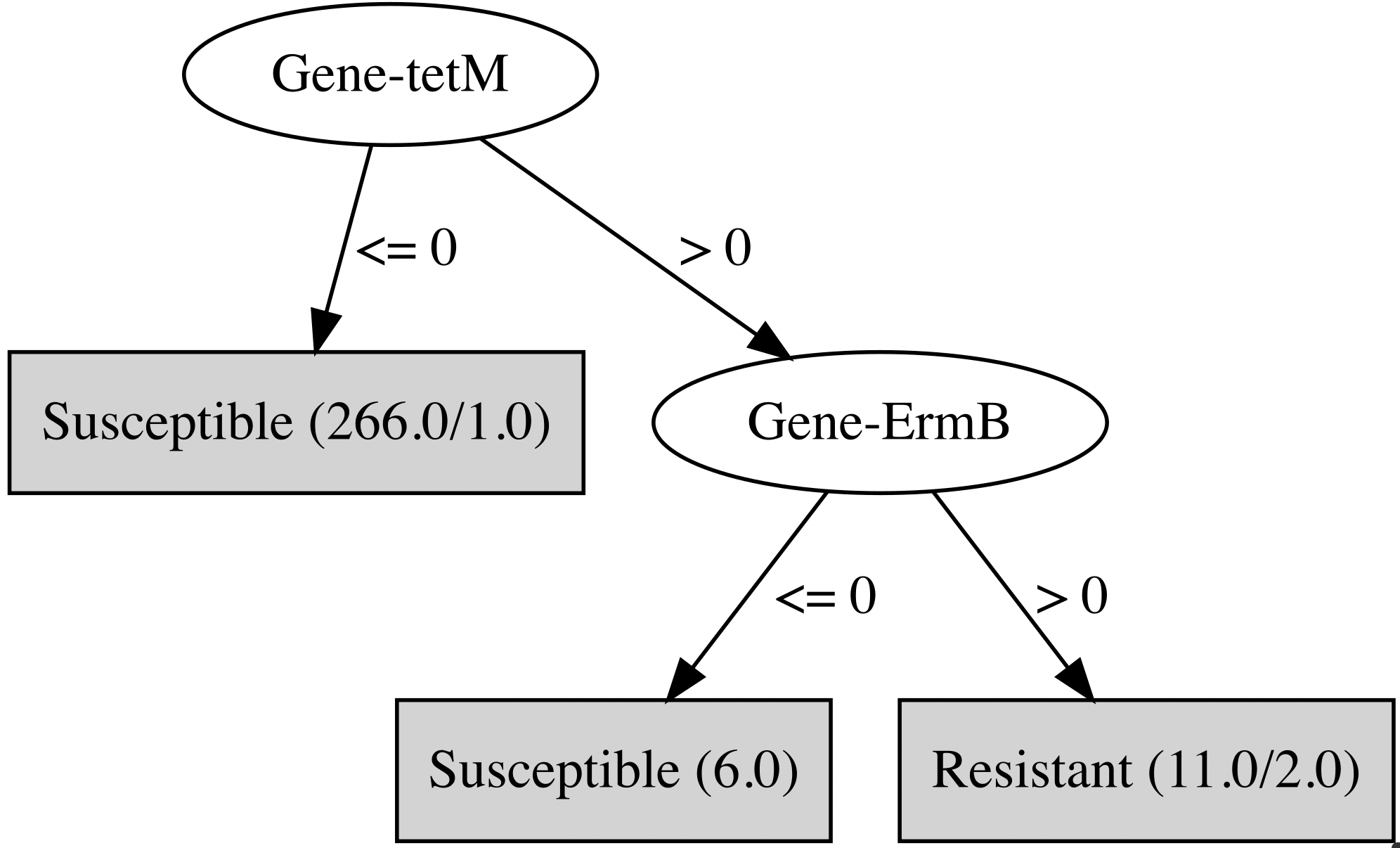

### Part_J_Colistin_RGI_all.png

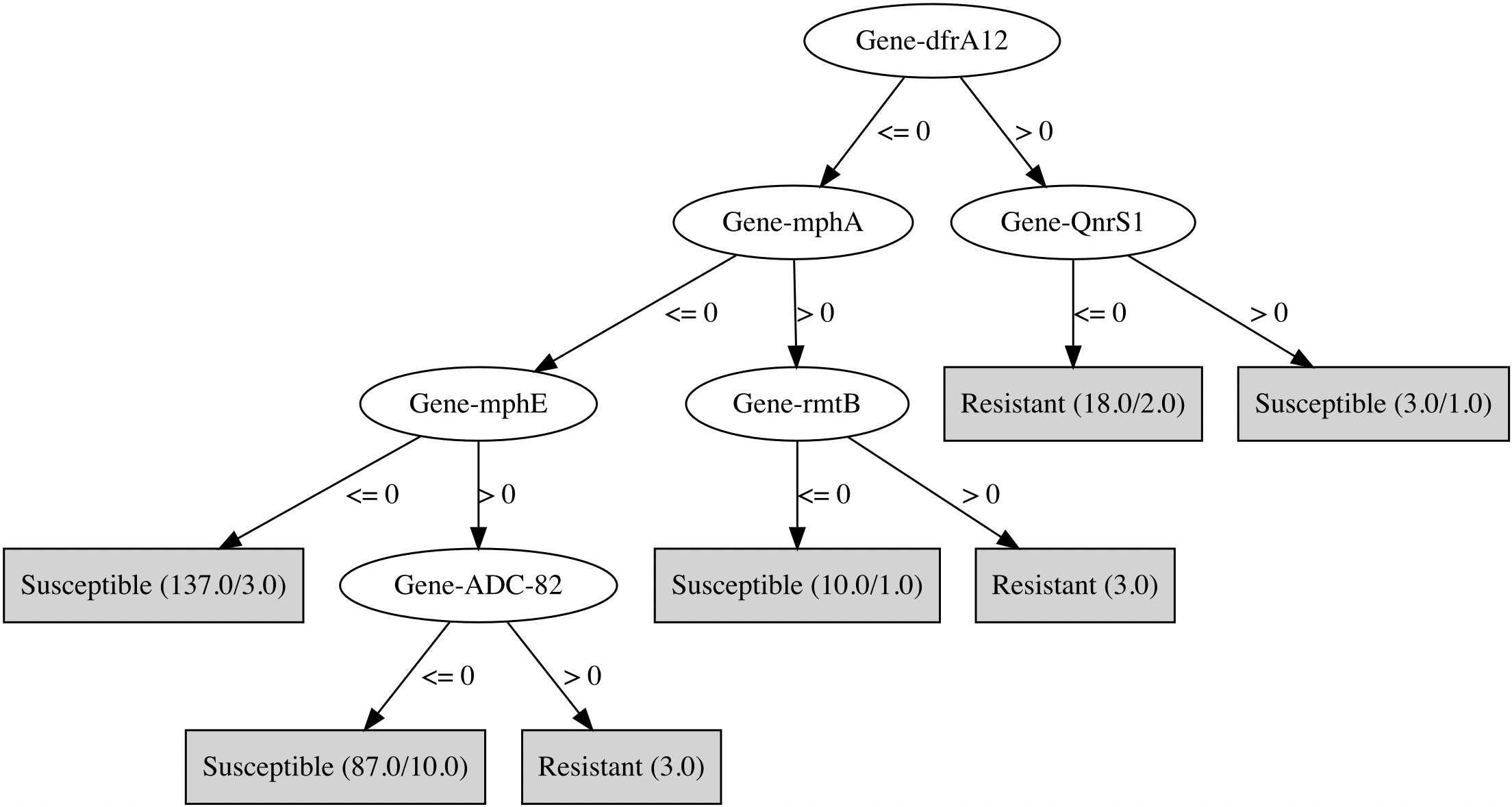

### Part_K_Doripenem_RGI_all.png

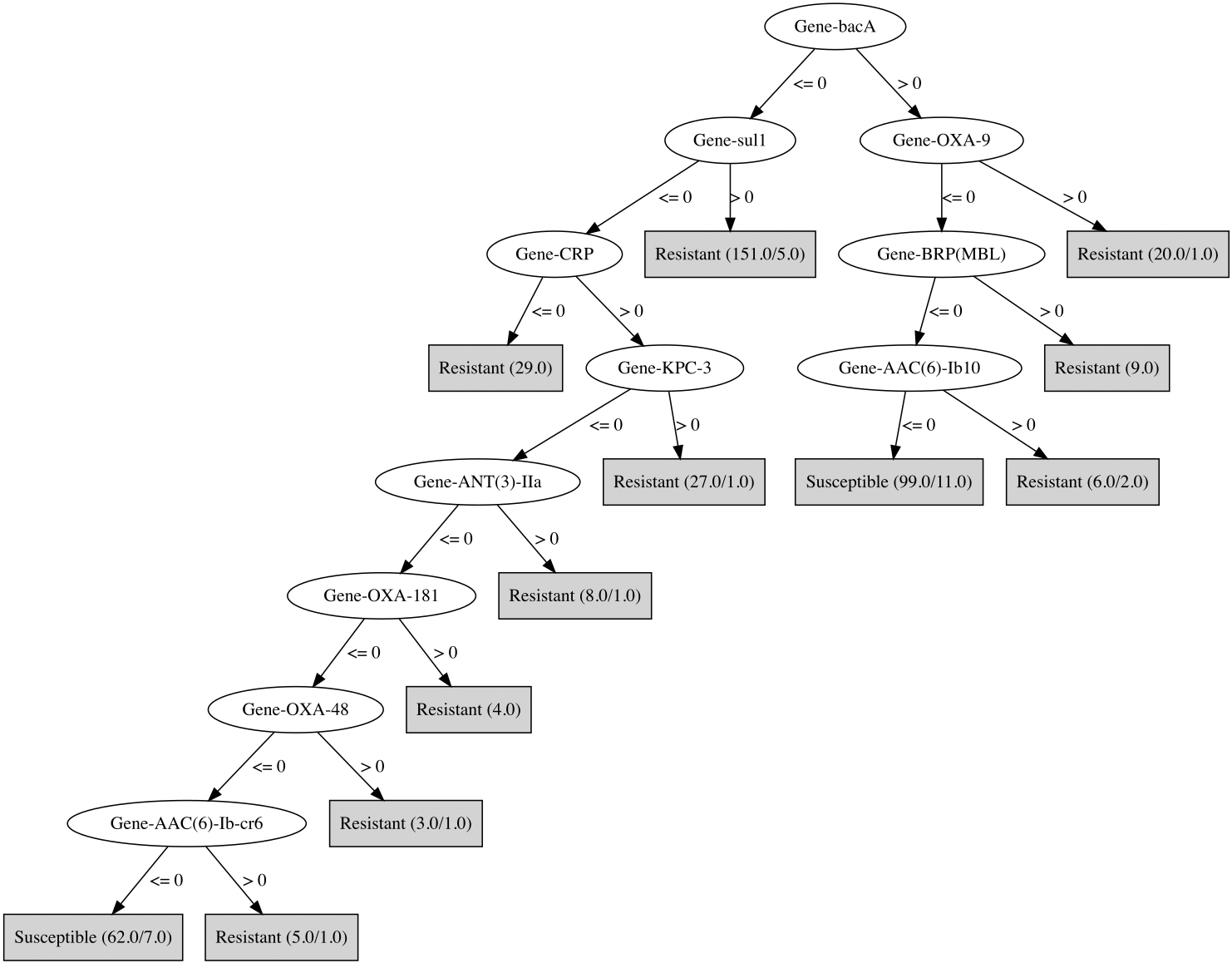

### Part_L_Ertapenem_RGI_all.png

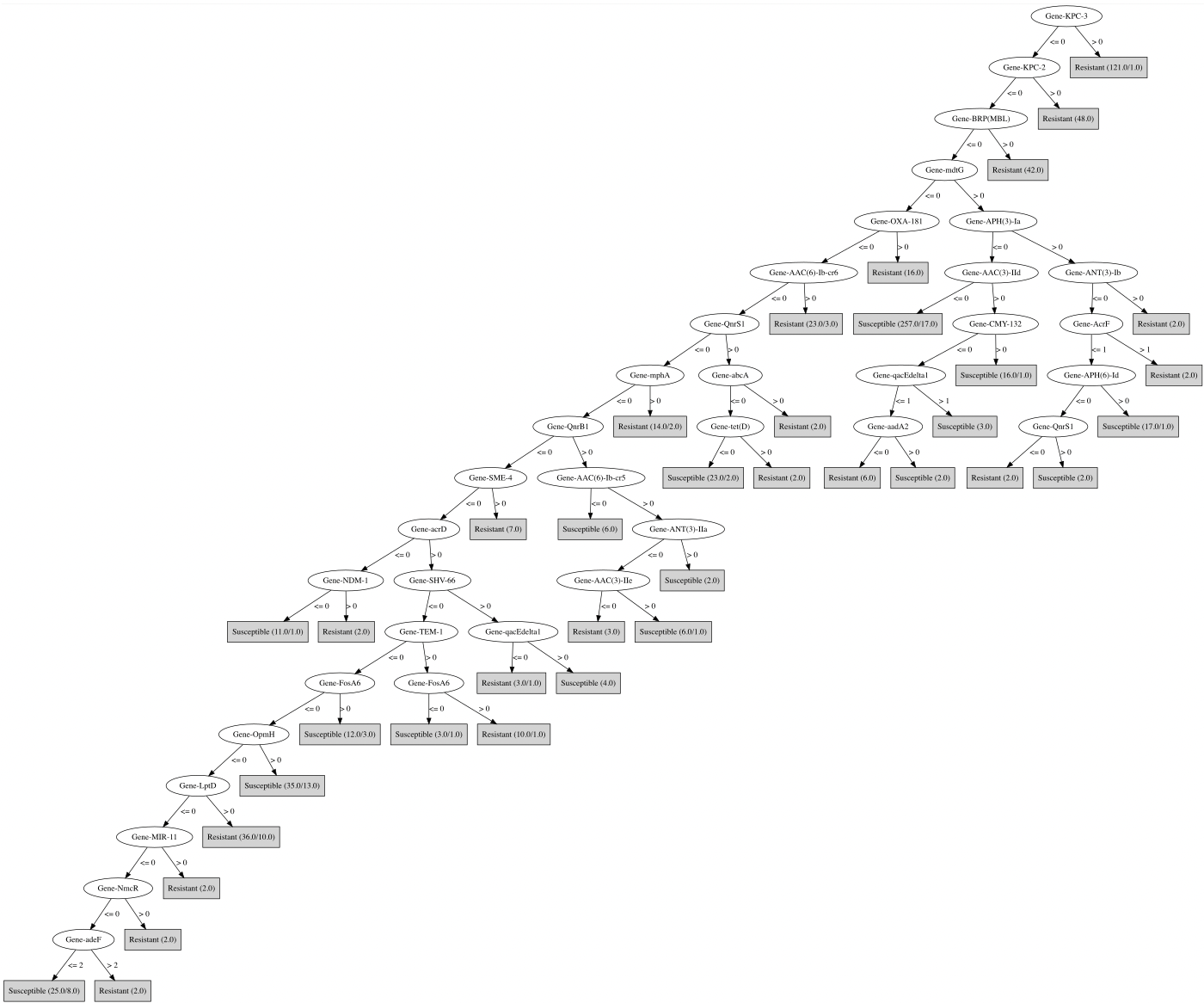

### Part_M_Erythromycin_RGI_all.png

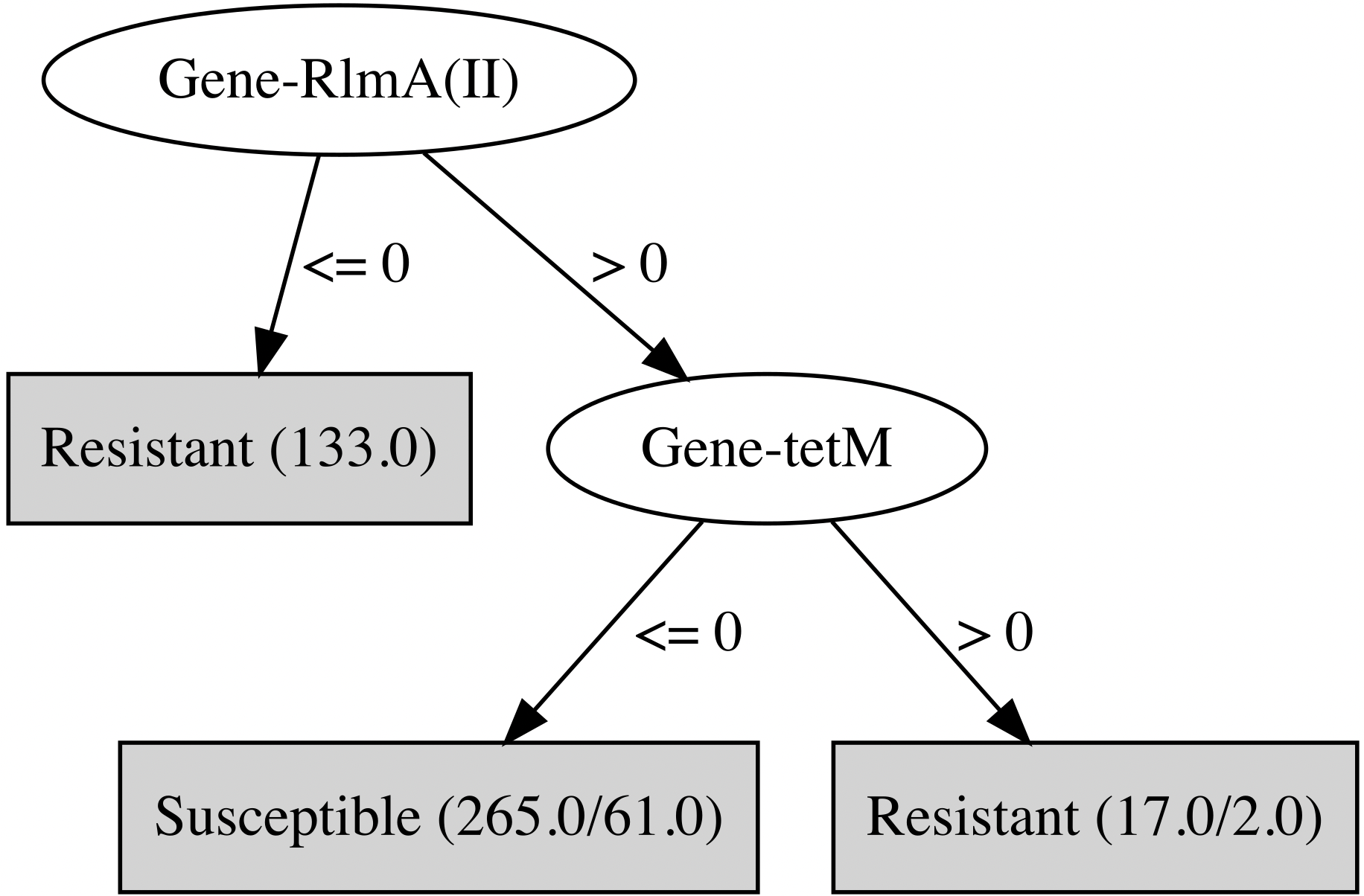

### Part_N_Fosfomycin_RGI_all.png

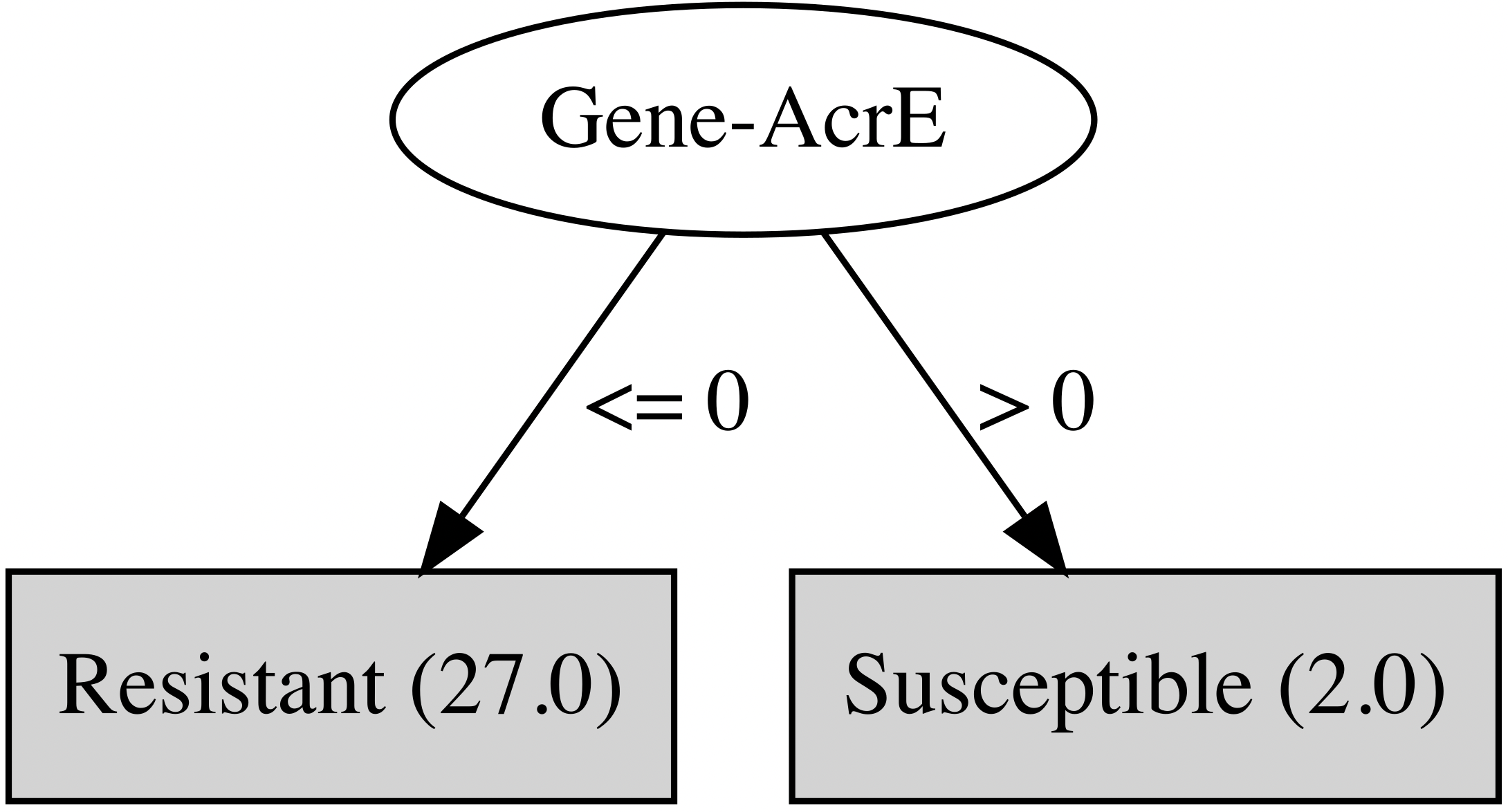

### Part_O_Gentamicin_RGI_all.png

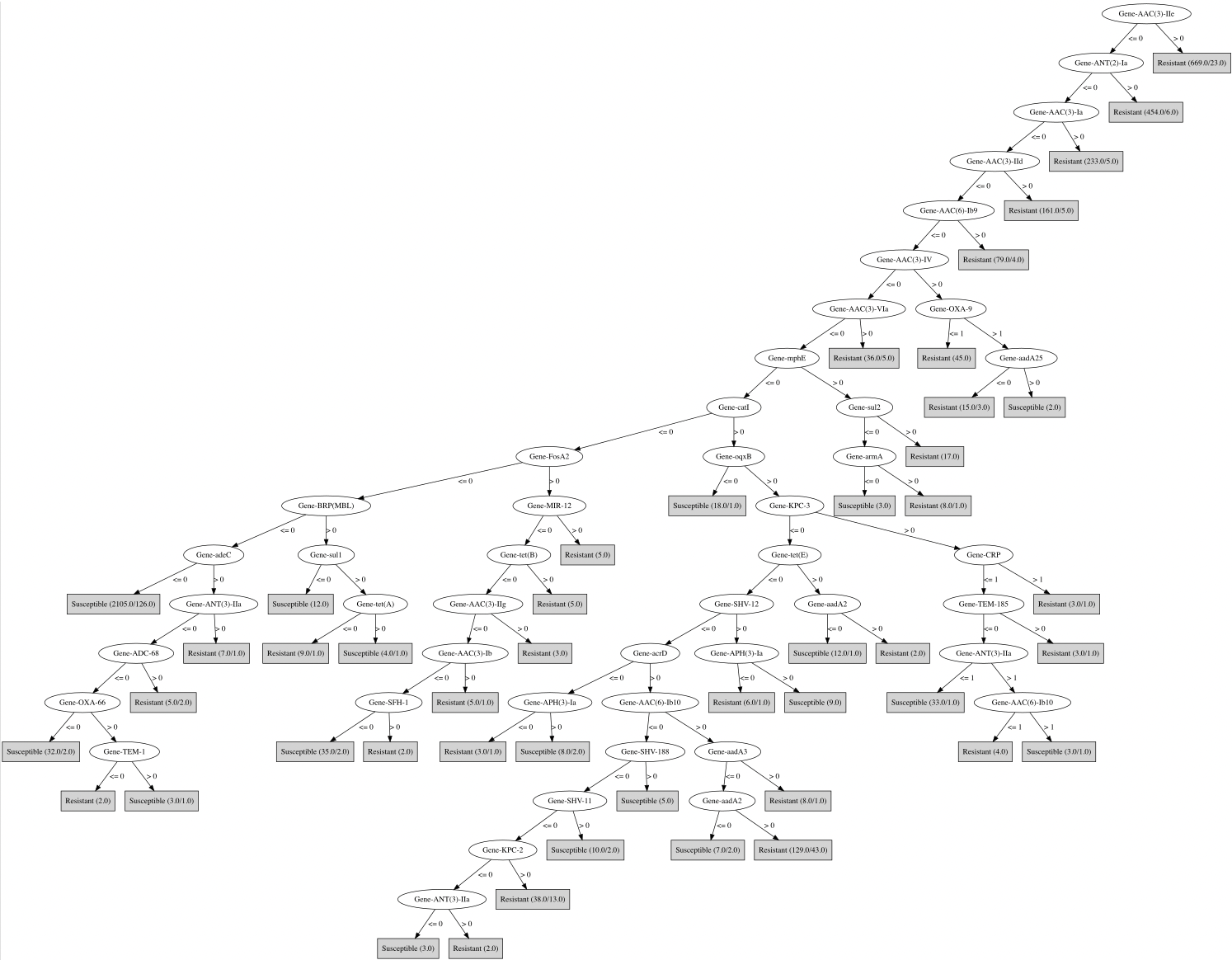

### Part_P_Imipenem_RGI_all.png

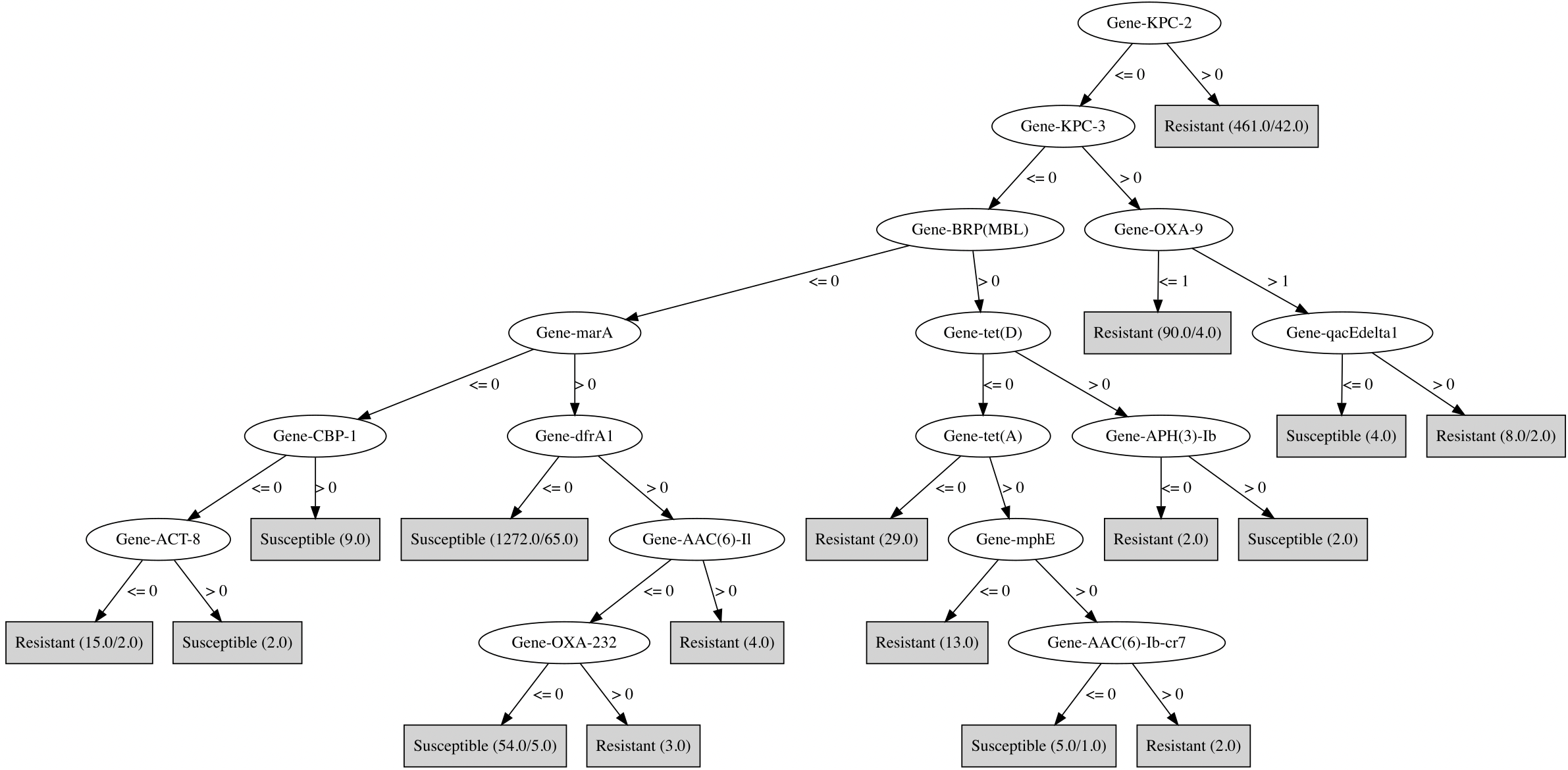

### Part_Q_Levofloxacin_RGI_all.png

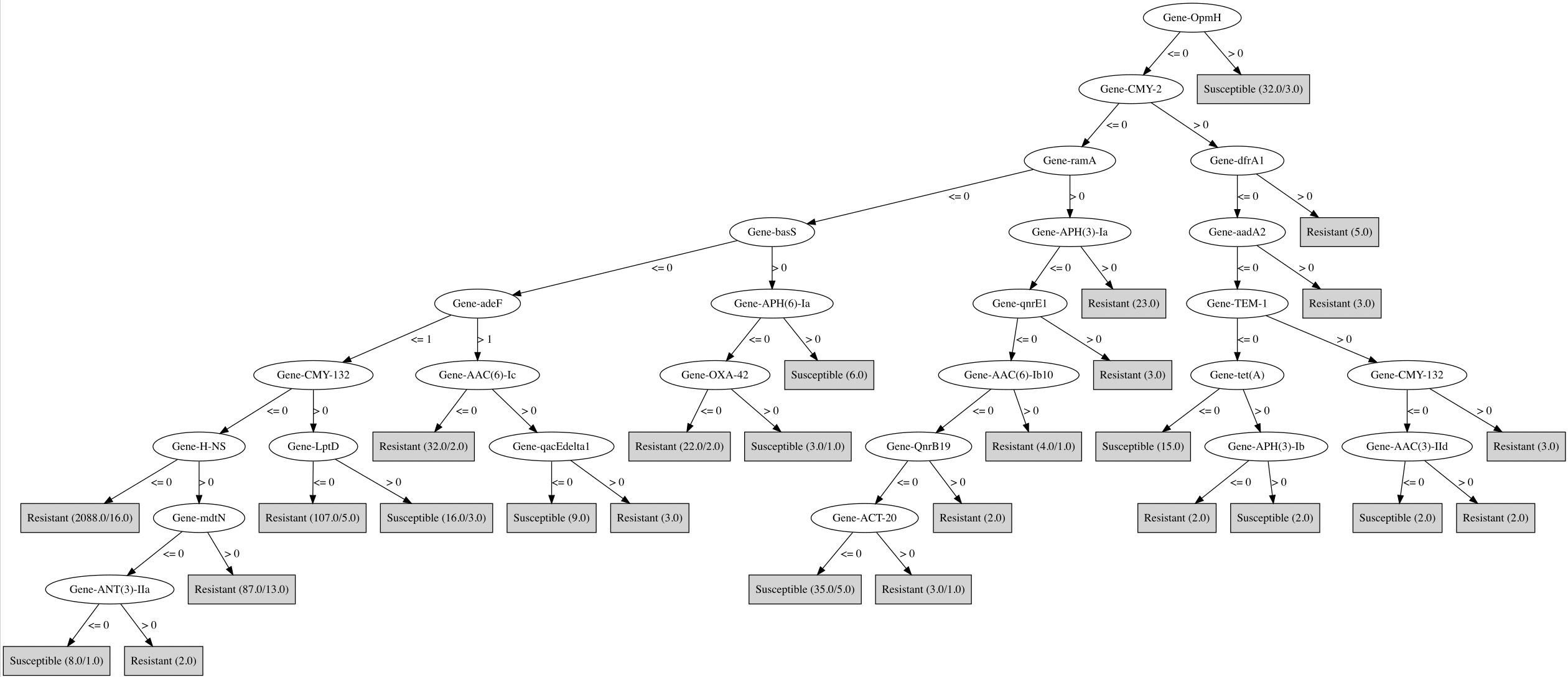

### Part_R_Meropenem_RGI_all.png

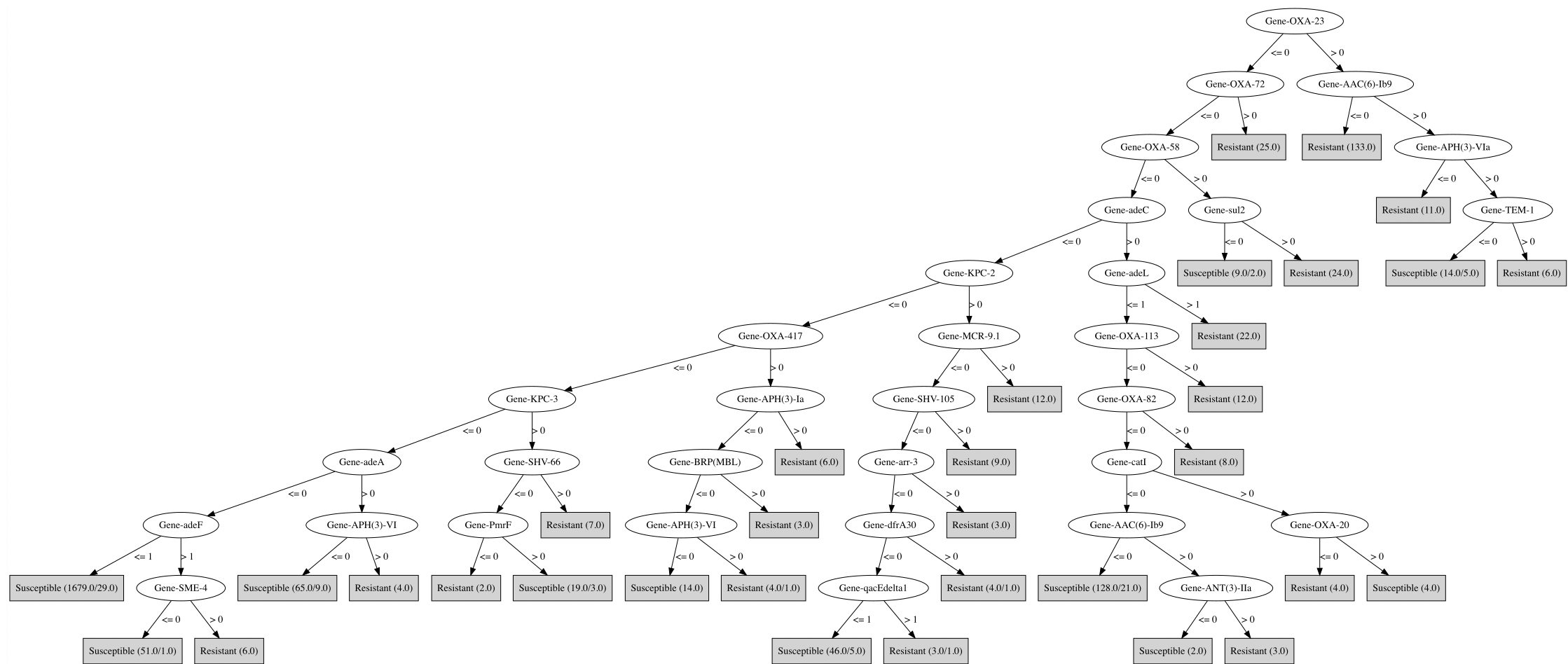

### Part_S_Moxifloxacin_RGI_all.png

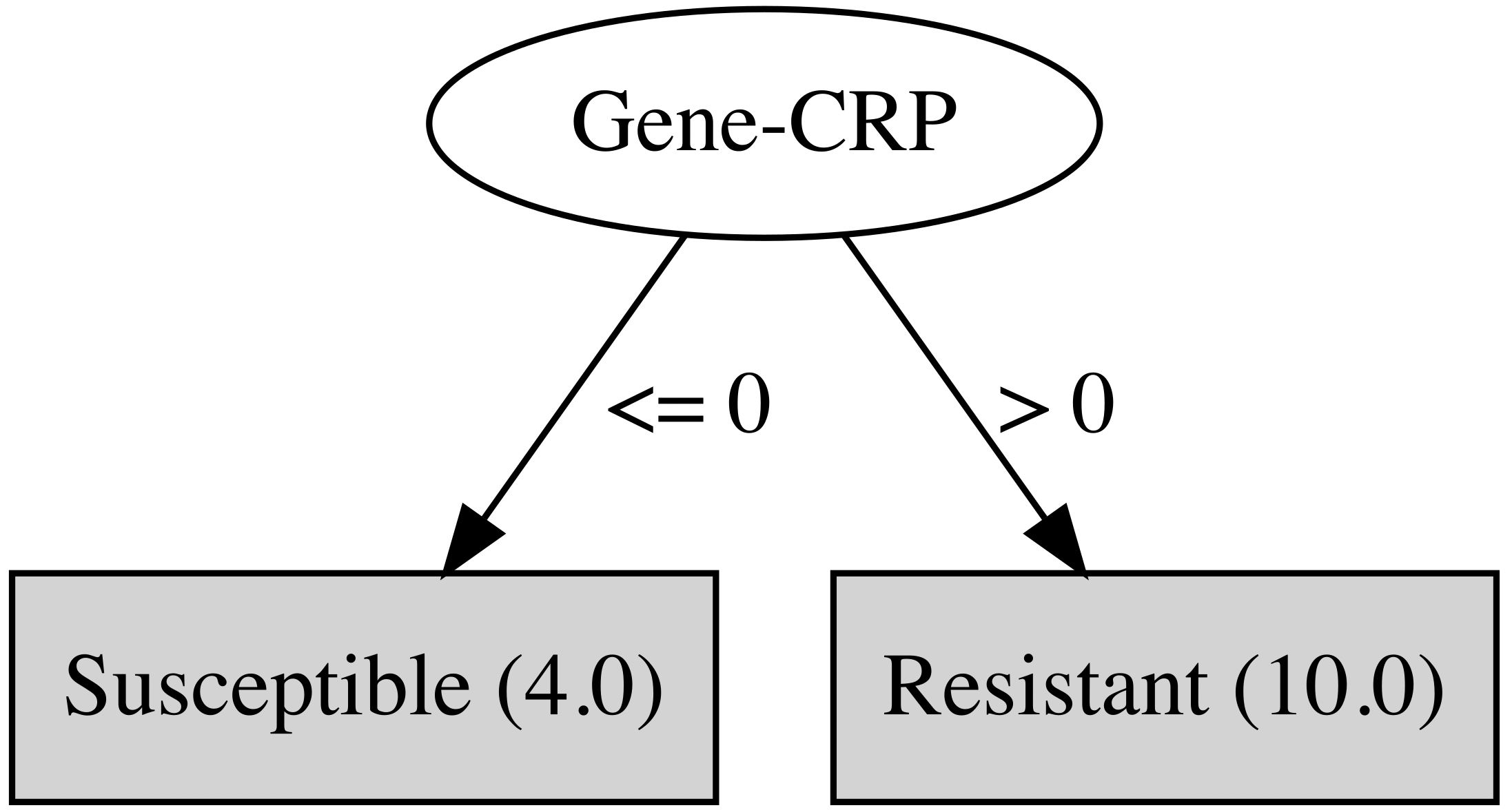

### Part_T_Nitrofurantoin_RGI_all.png

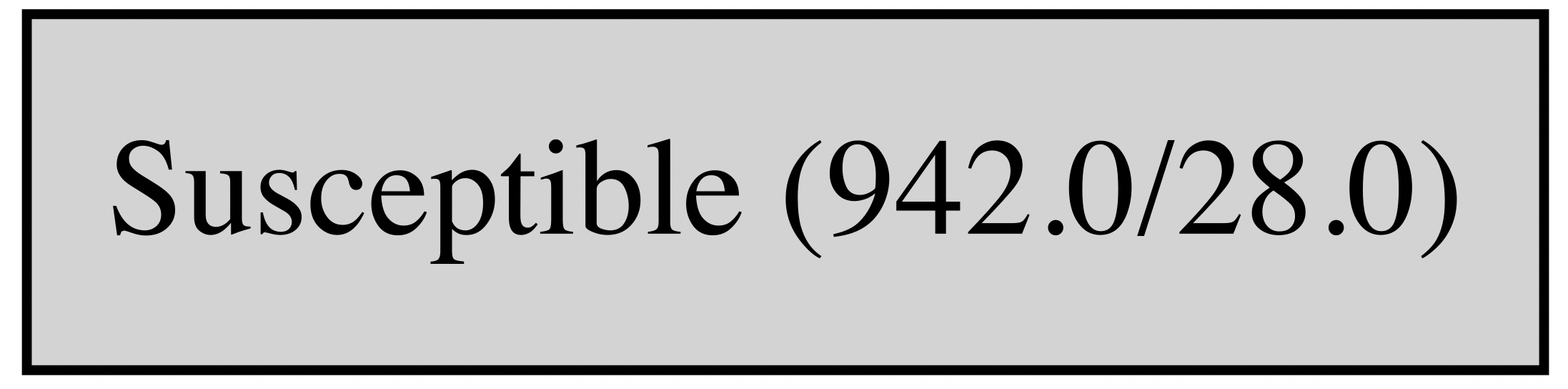

### Part_U_Tetracycline_RGI_all.png

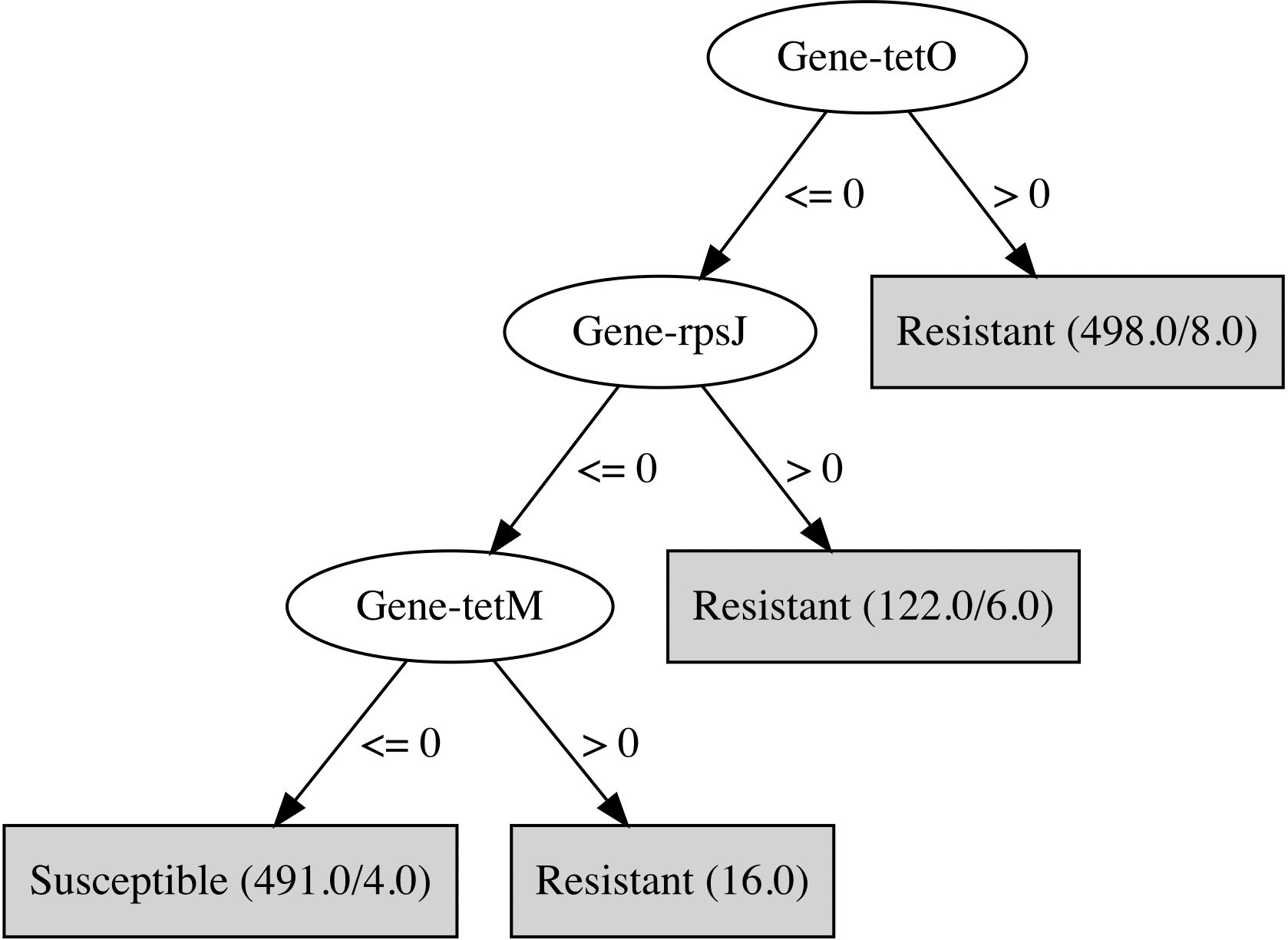

### Part_V_Tigecycline_RGI_all.png

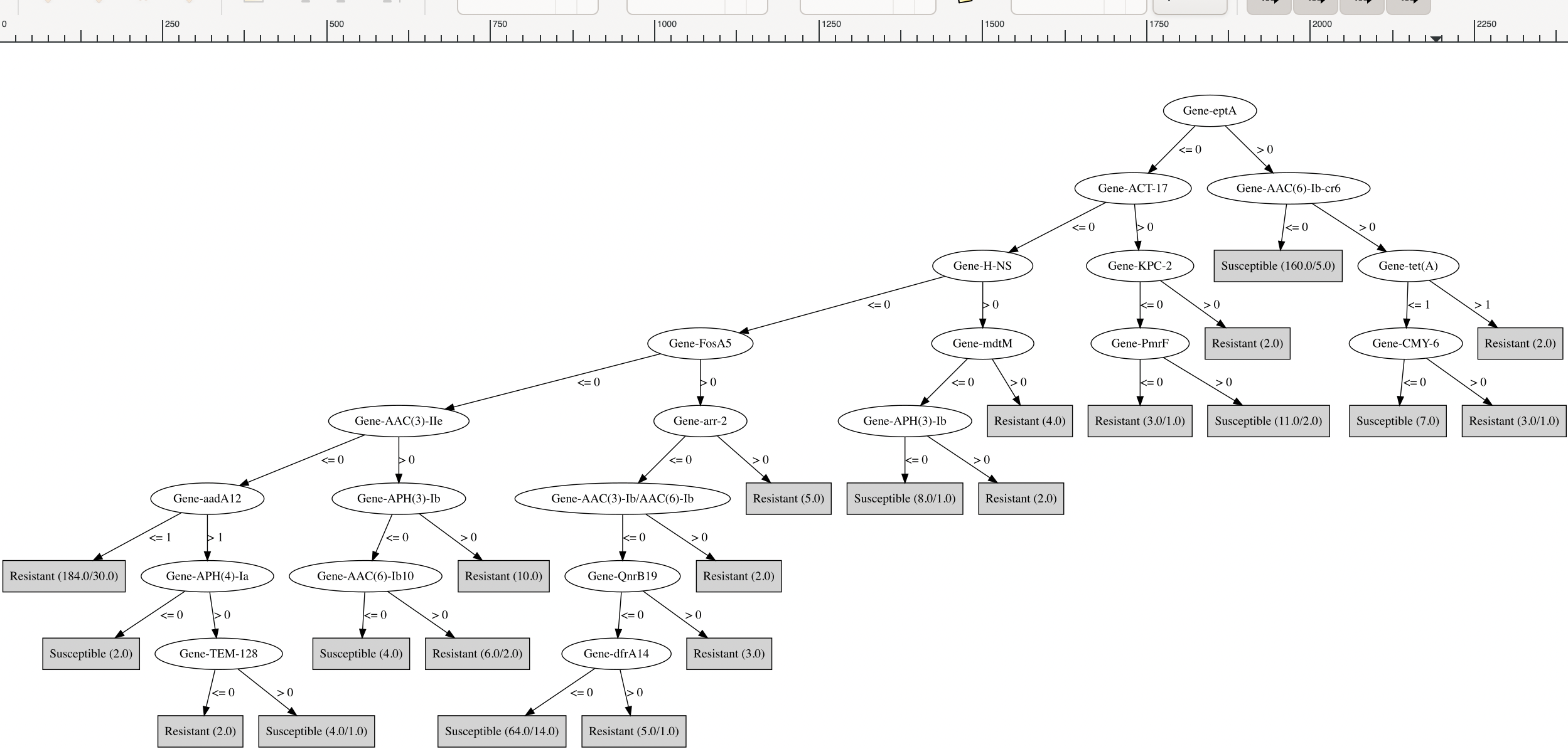

### Part_W_Tobramycin_RGI_all.png

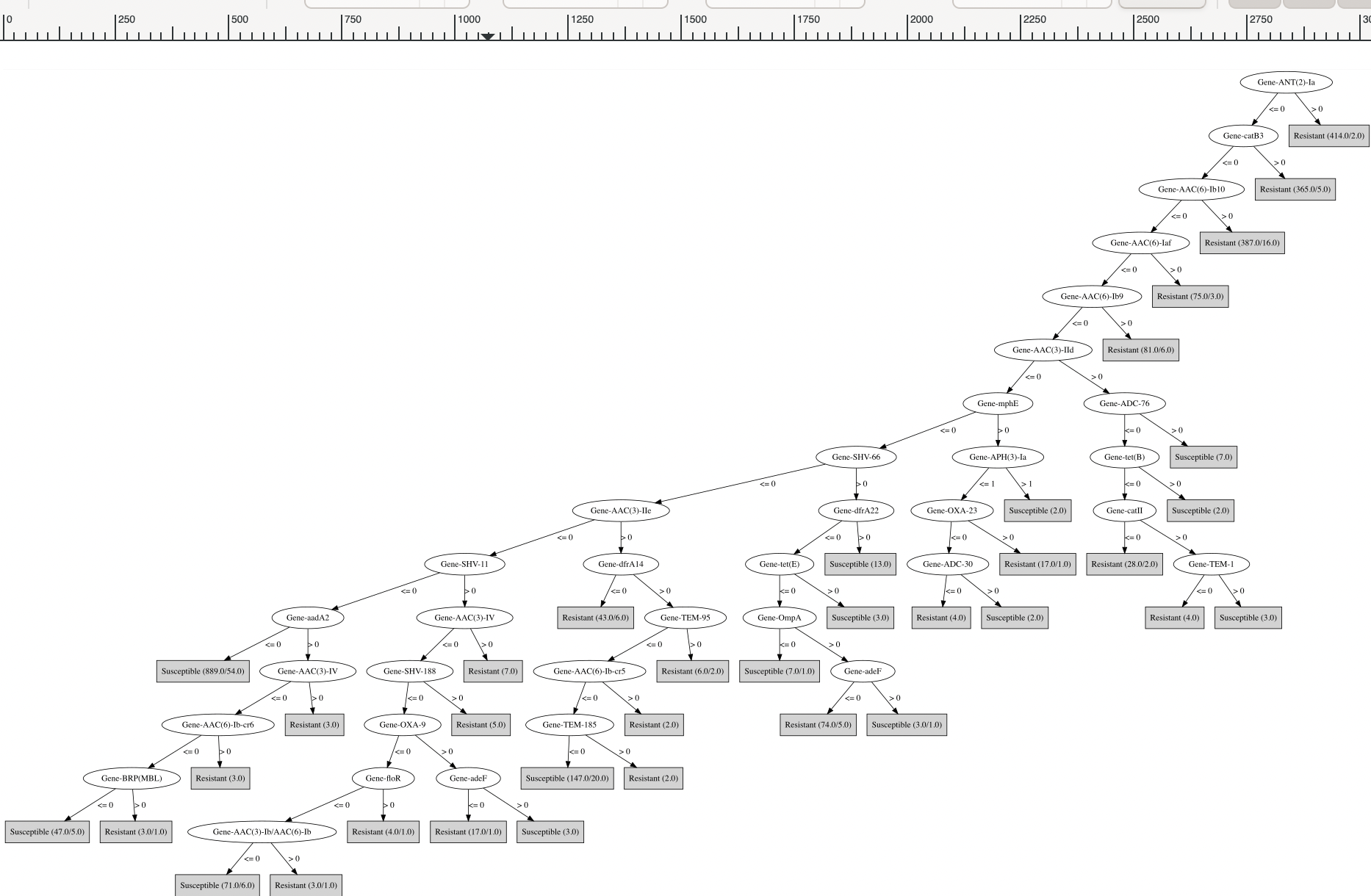

### PartA_Amikacin_RGI_specific.png

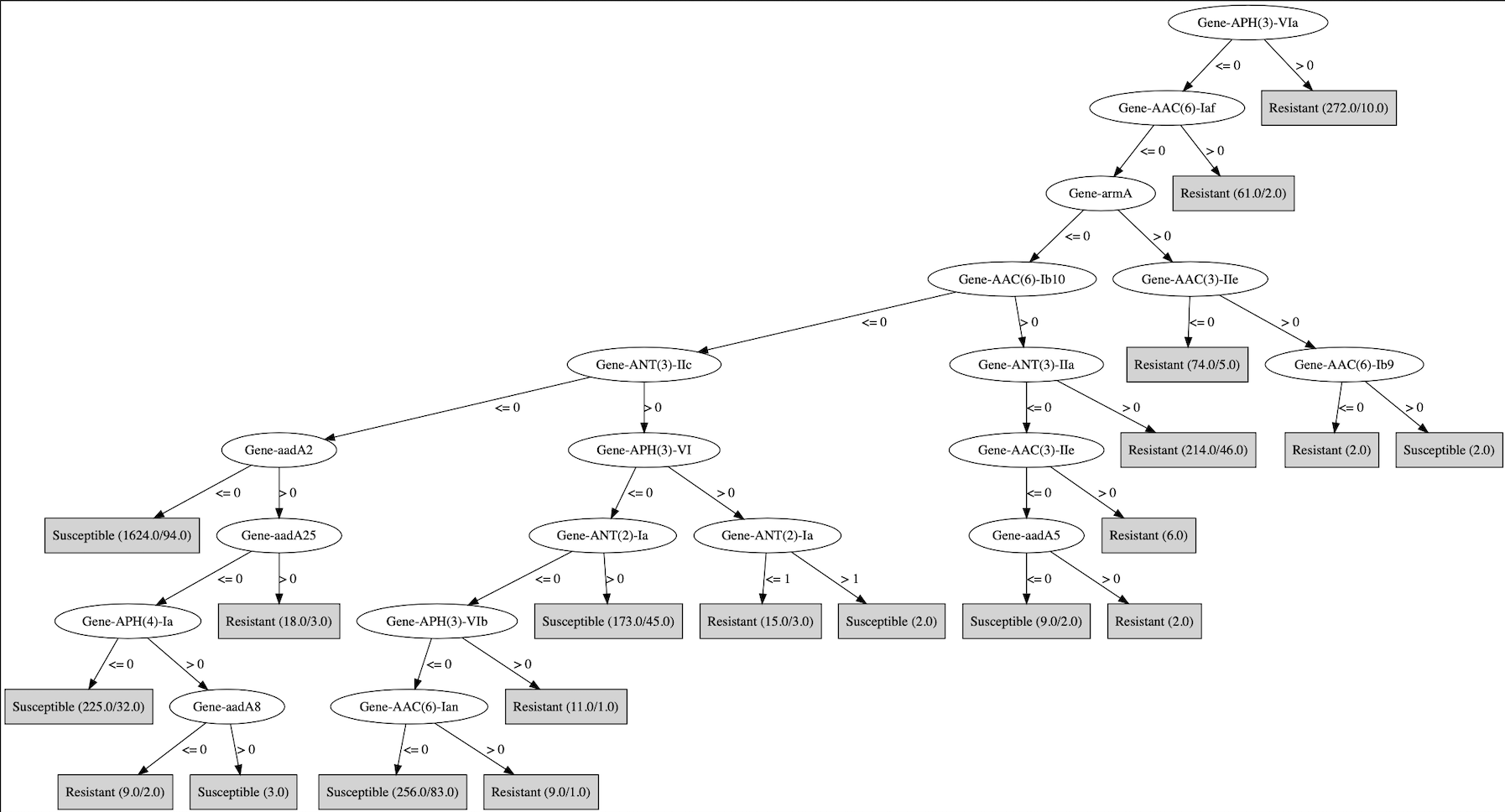

### PartB_Amoxicillin_RGI_specific.png

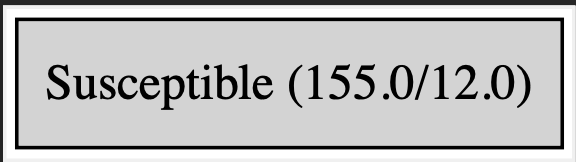

### PartC_Ampicillin_RGI_specific.png

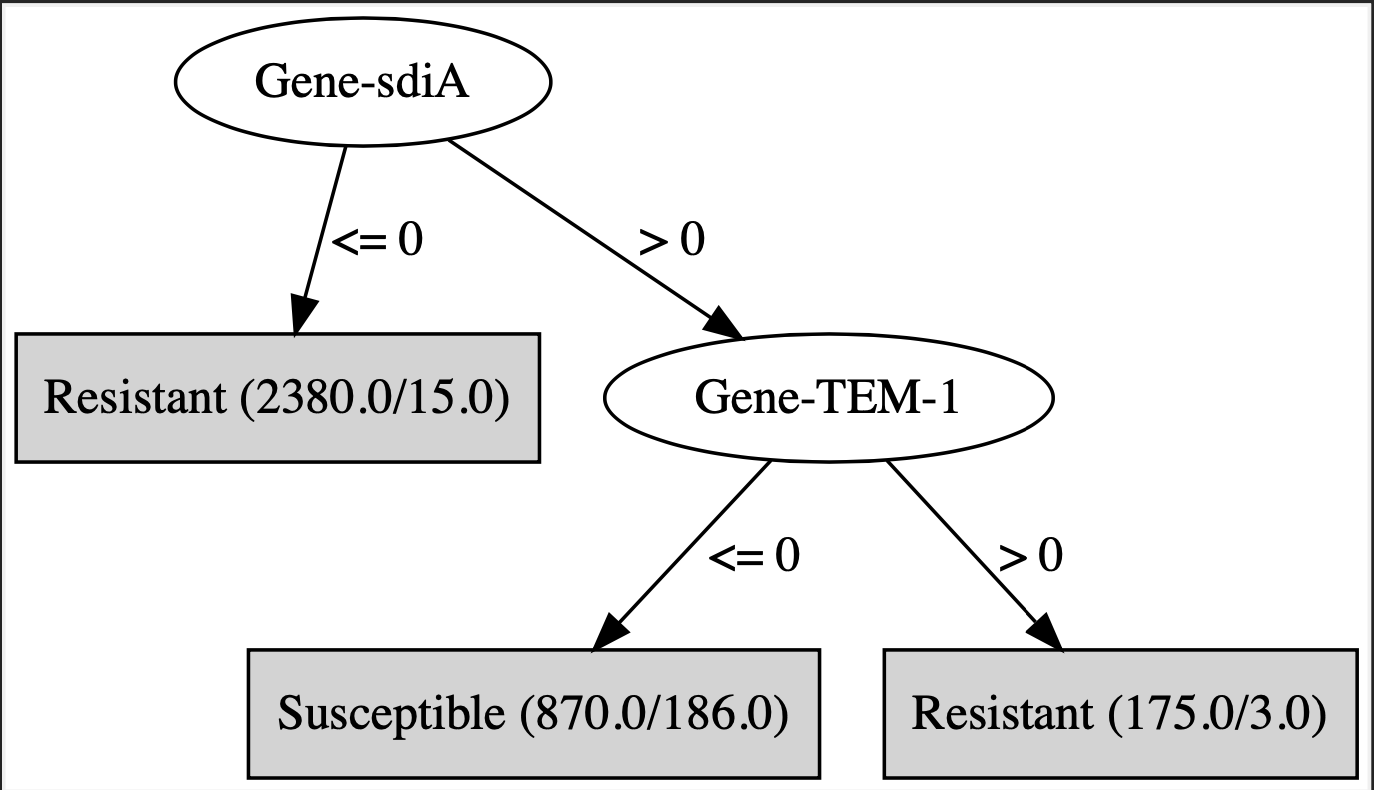

### PartD_Aztreonam_RGI_specific.png

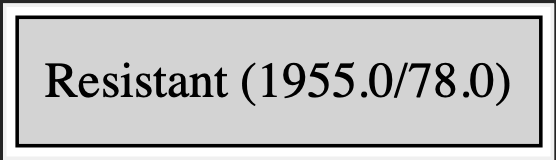

### PartE_Cefepime_RGI_specific.png

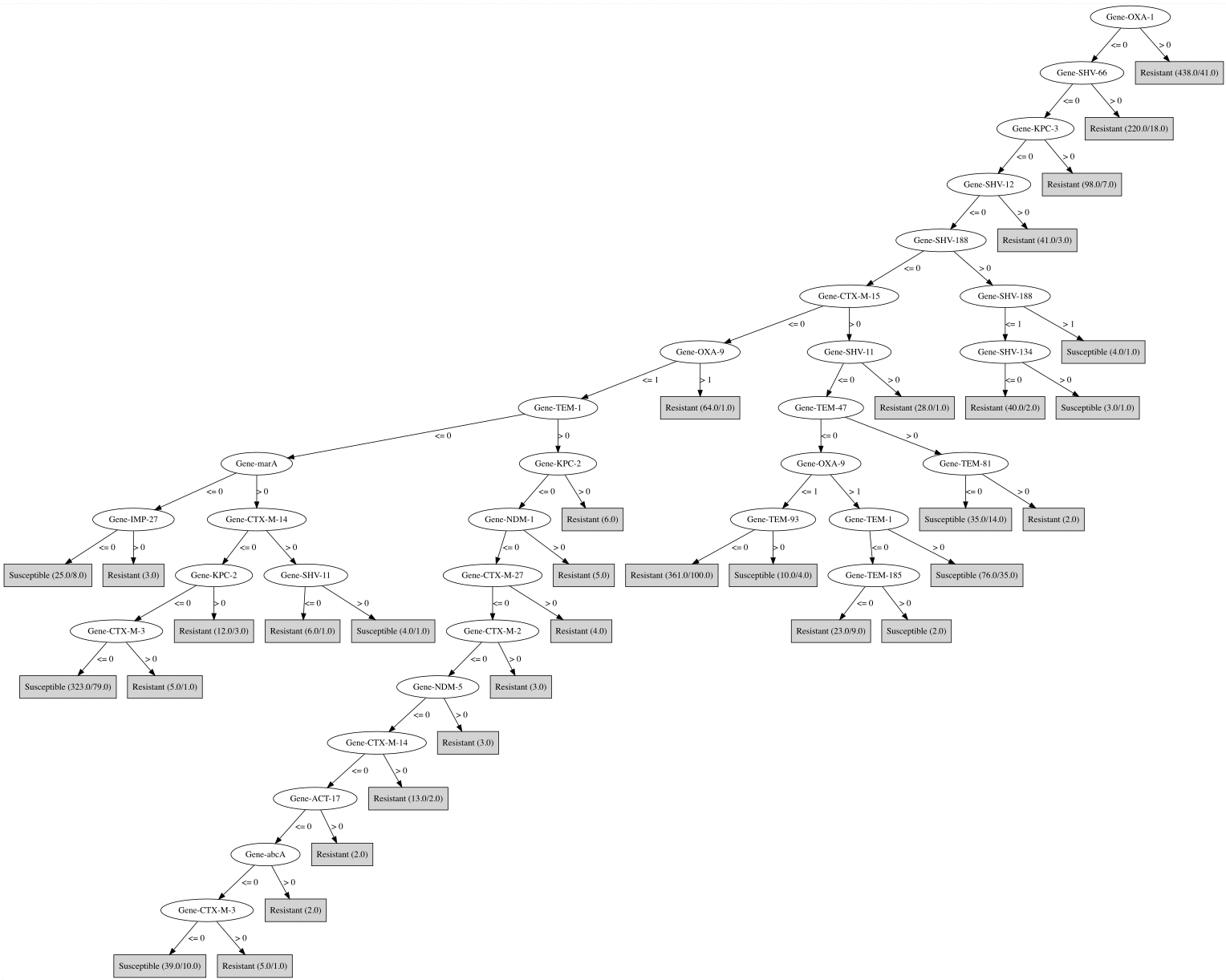

### PartF_Ceftriaxone_RGI_specific.png

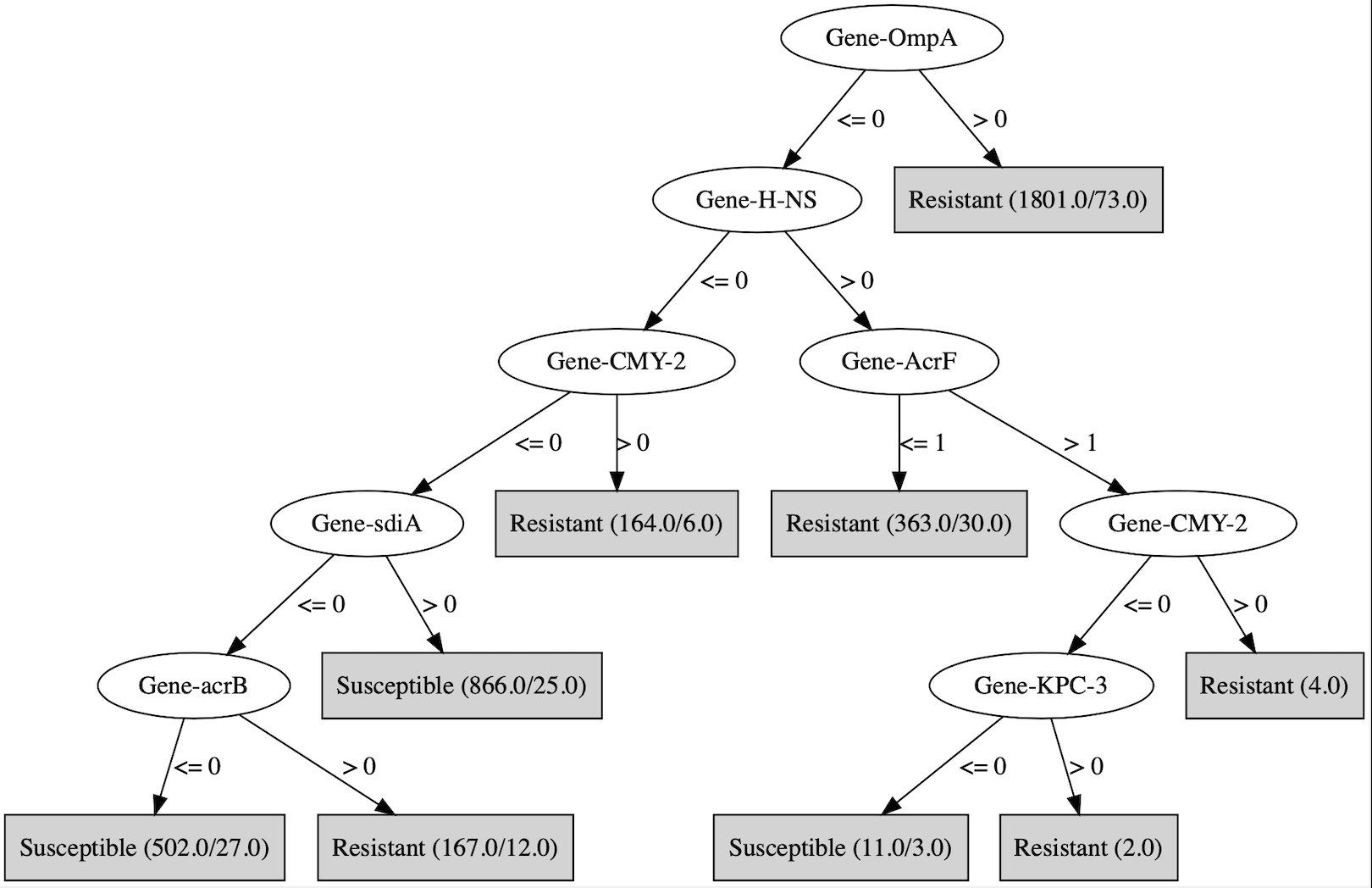

### PartG_Chloramphenicol_RGI_specific.png

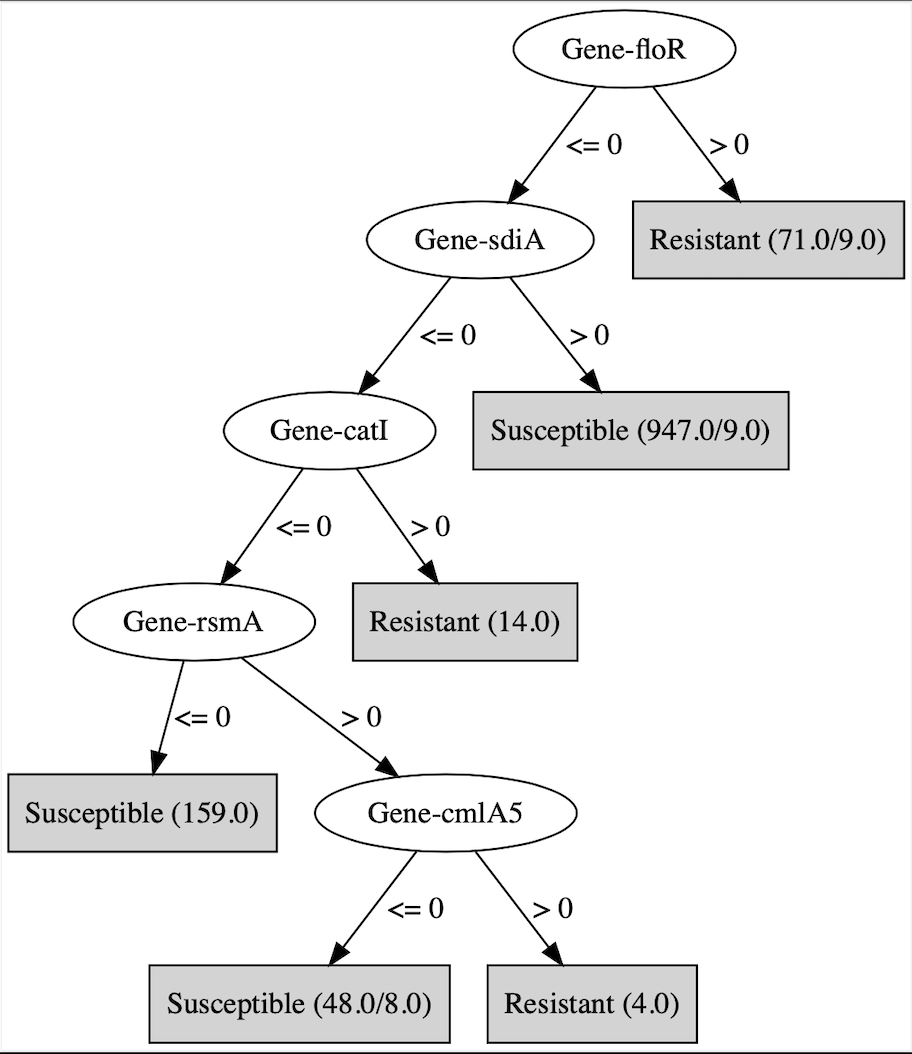

### PartH_Ciprofloxacin_RGI_specific.png

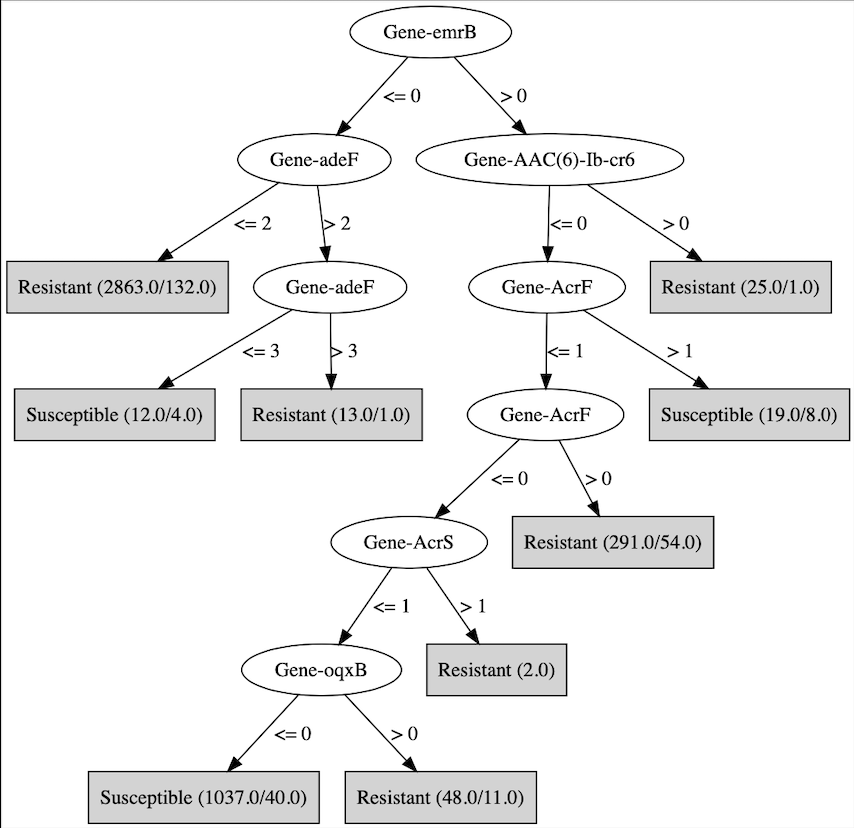

### PartI_Clindamycin_RGI_specific.png

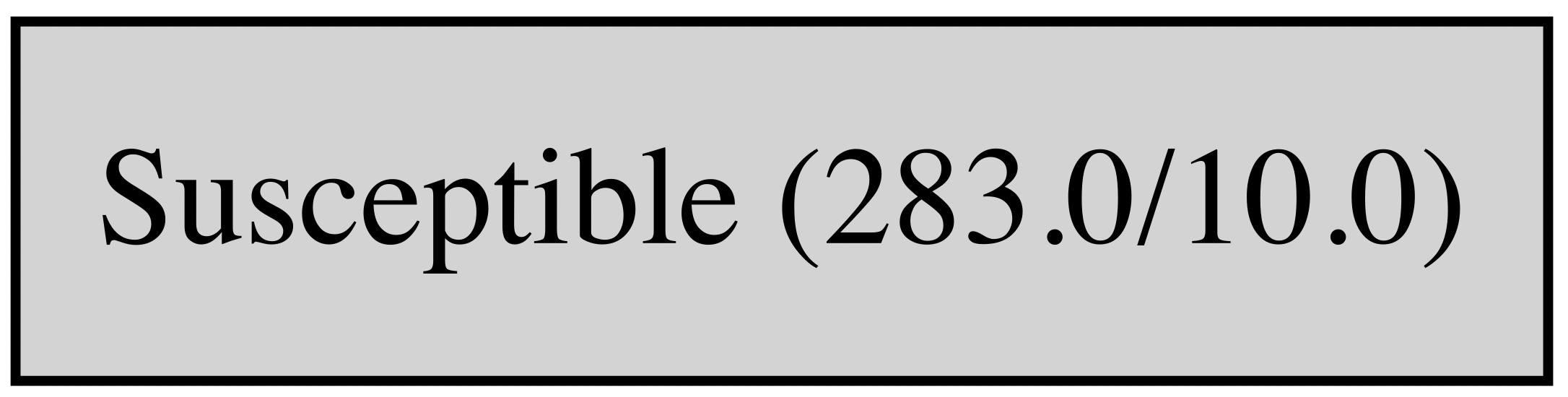

### PartJ_Colistin_RGI_specific.png

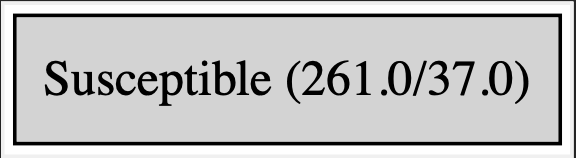

### PartK_Doripenem_RGI_specific.png

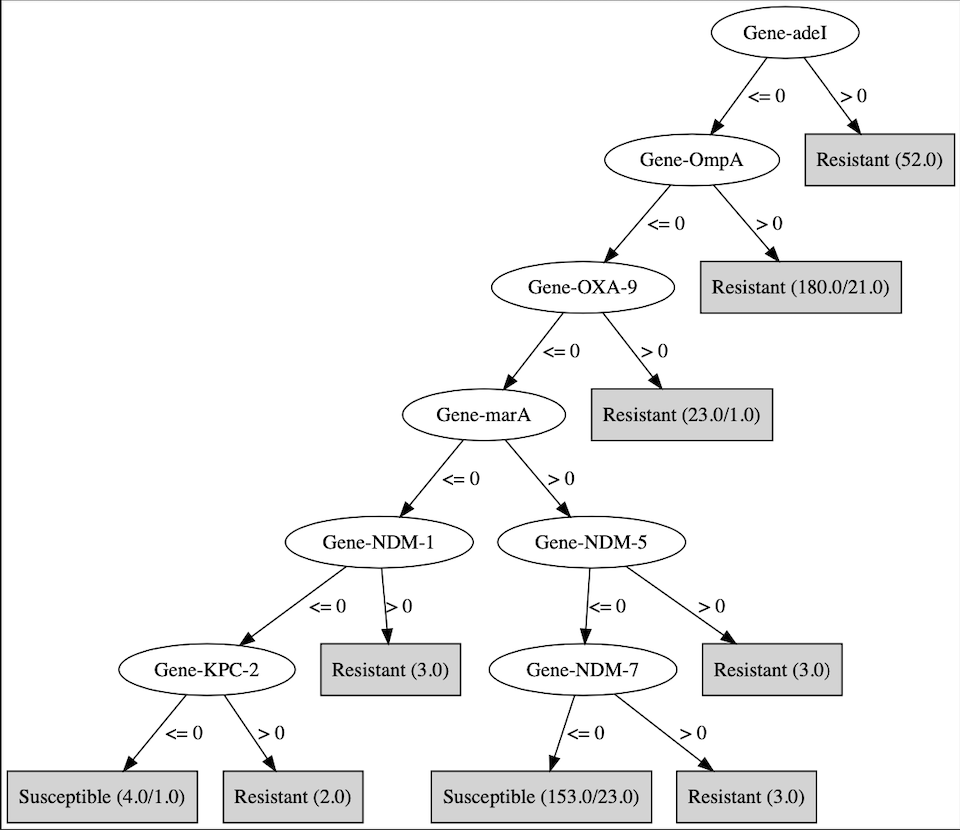

### PartL_Ertapenem_RGI_specific.png

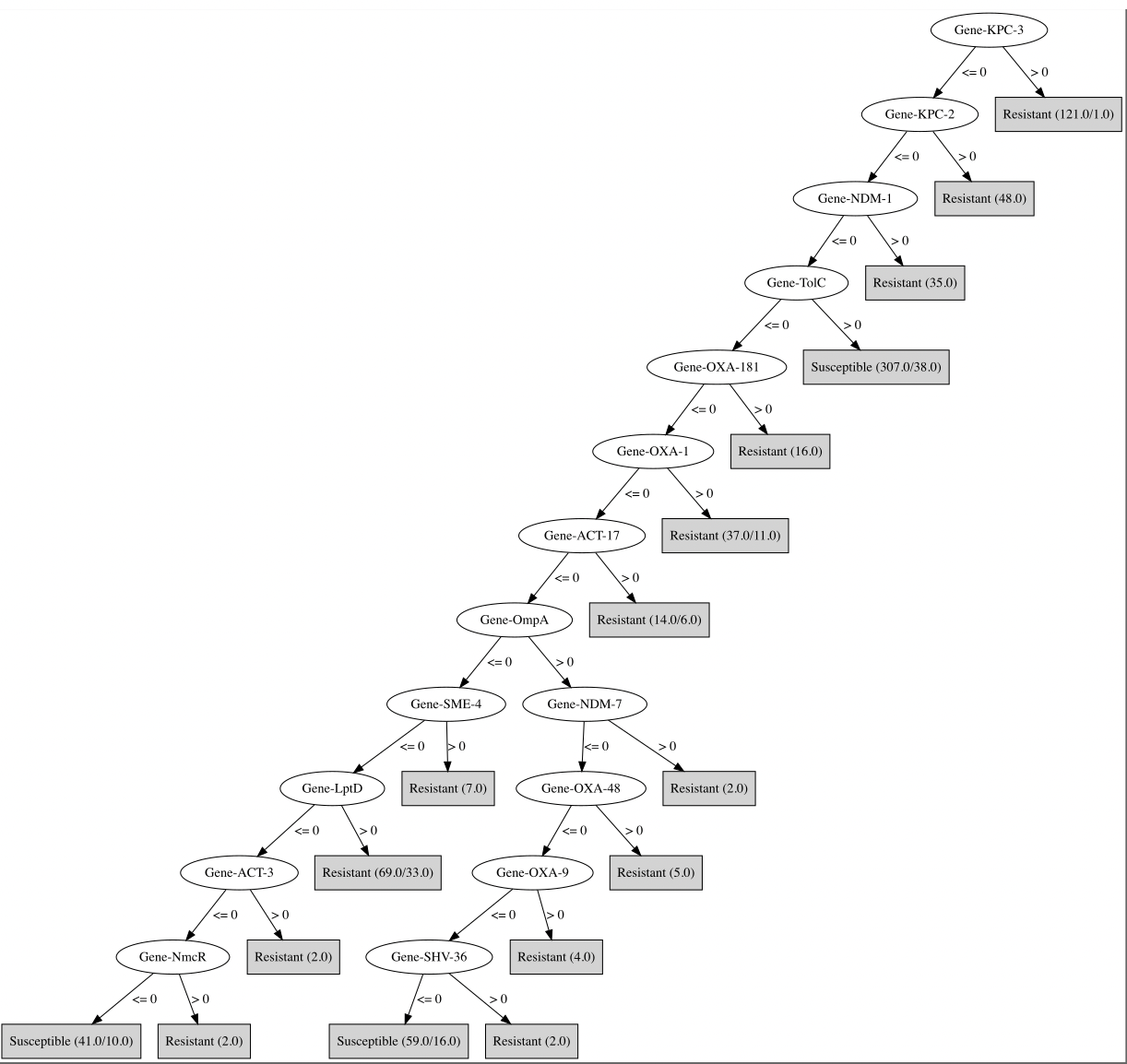

### PartM_Erythromycin_RGI_specific.png

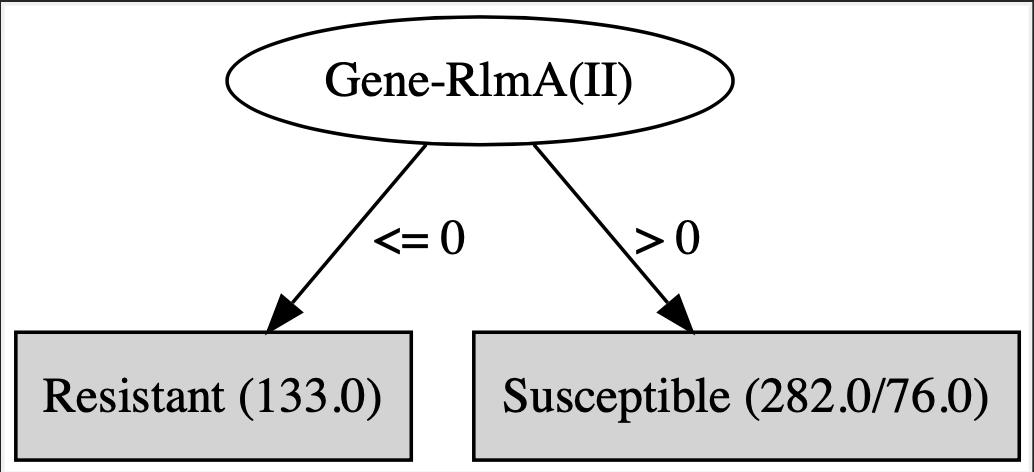

### PartN_Fosfomycin_RGI_specific.png

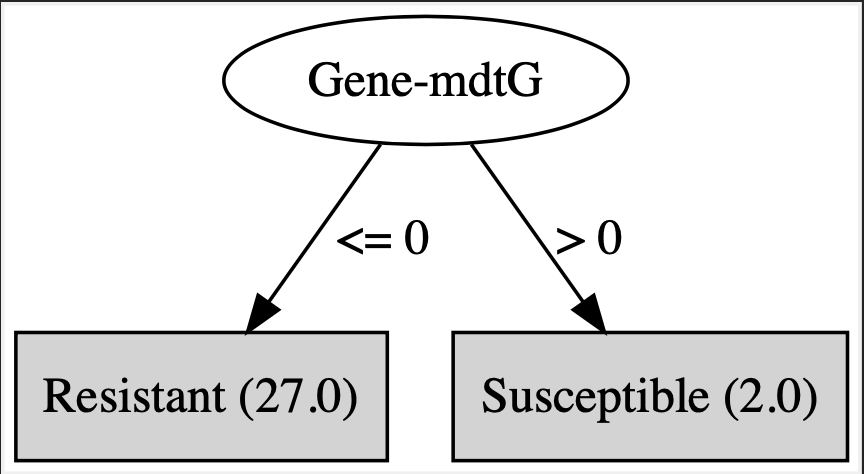

### PartO_Gentamicin_RGI_specific.png

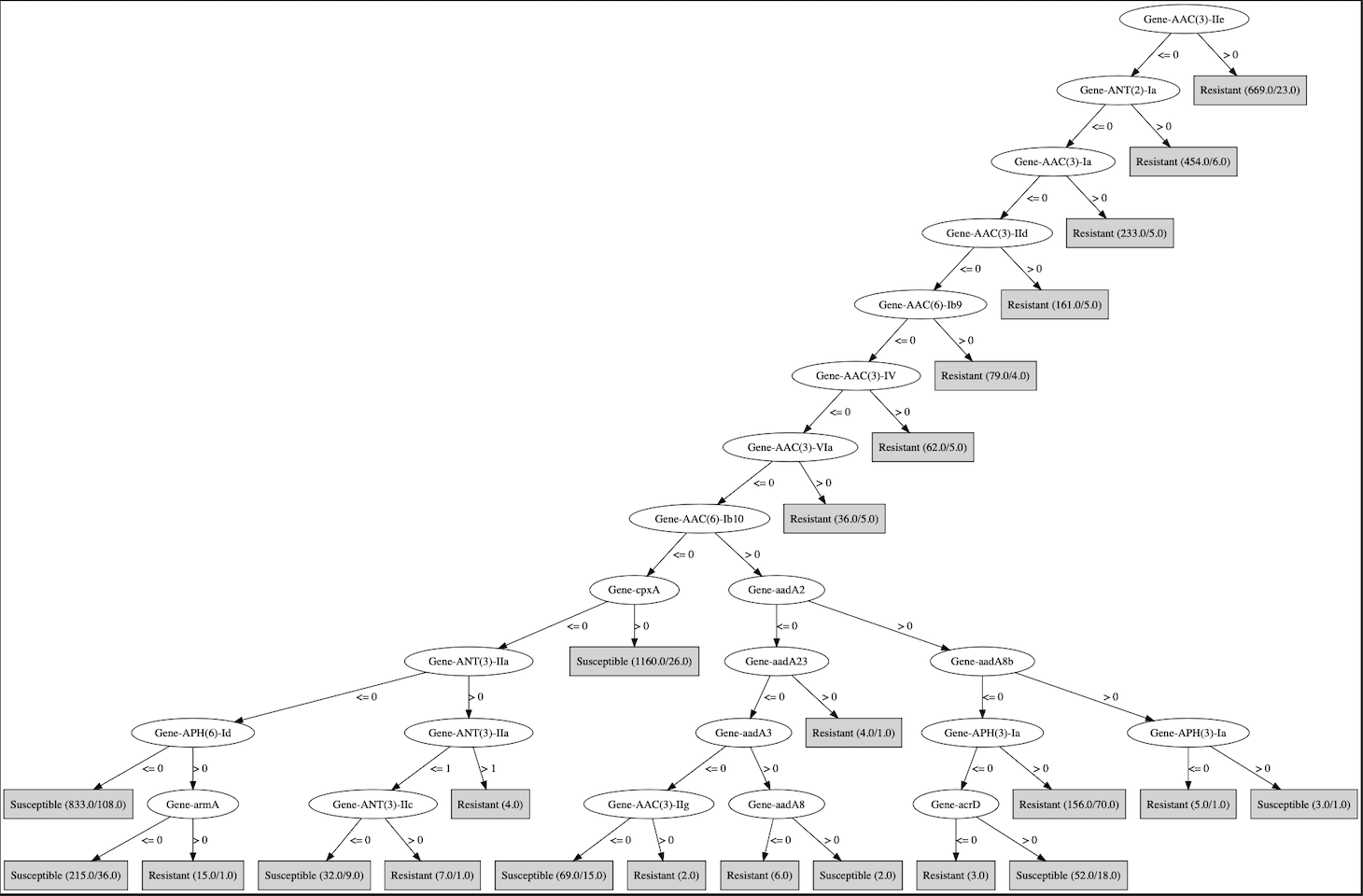

### PartP_Imipenem_RGI_specific.png

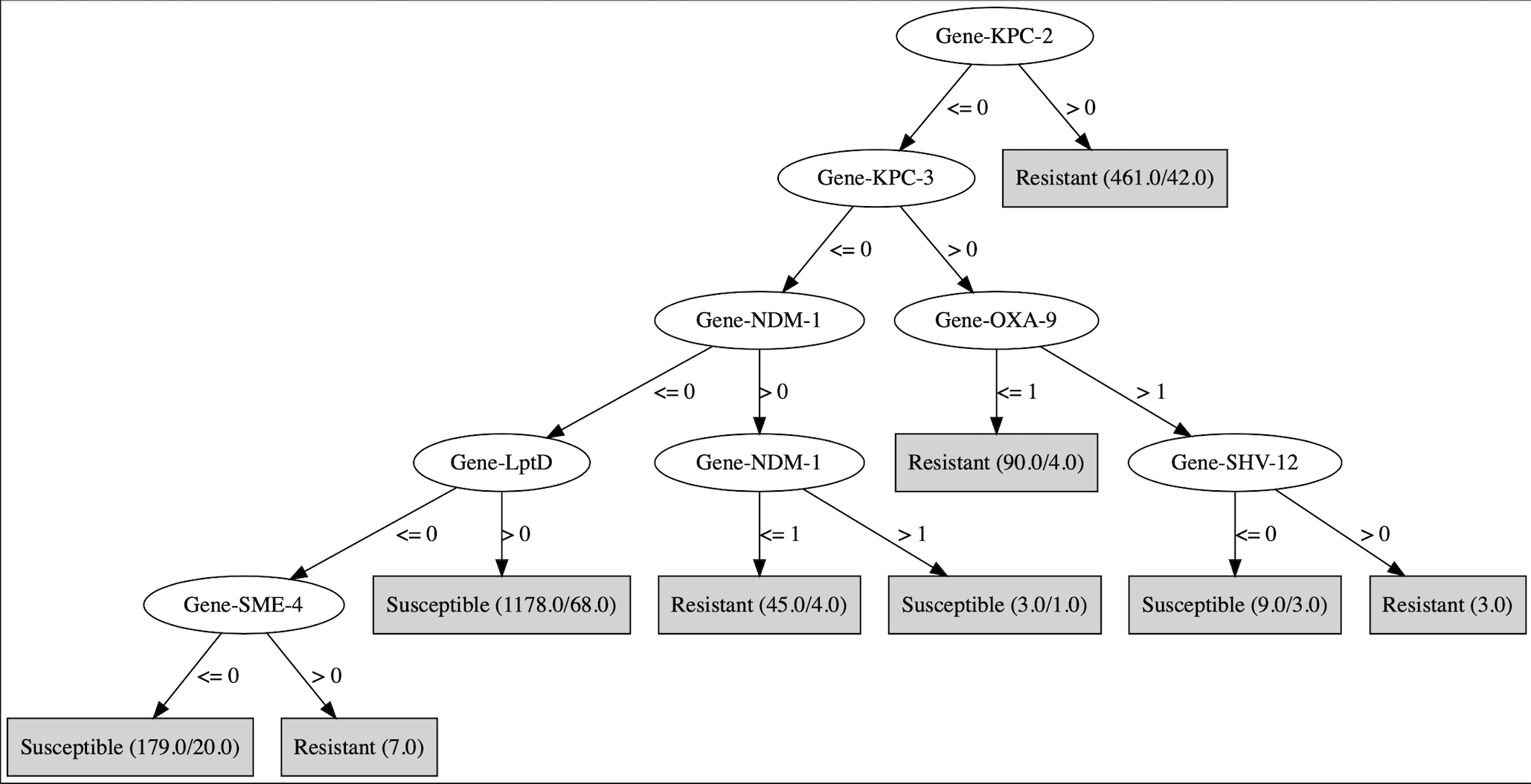

### PartQ_Levofloxacin_RGI_specific.png

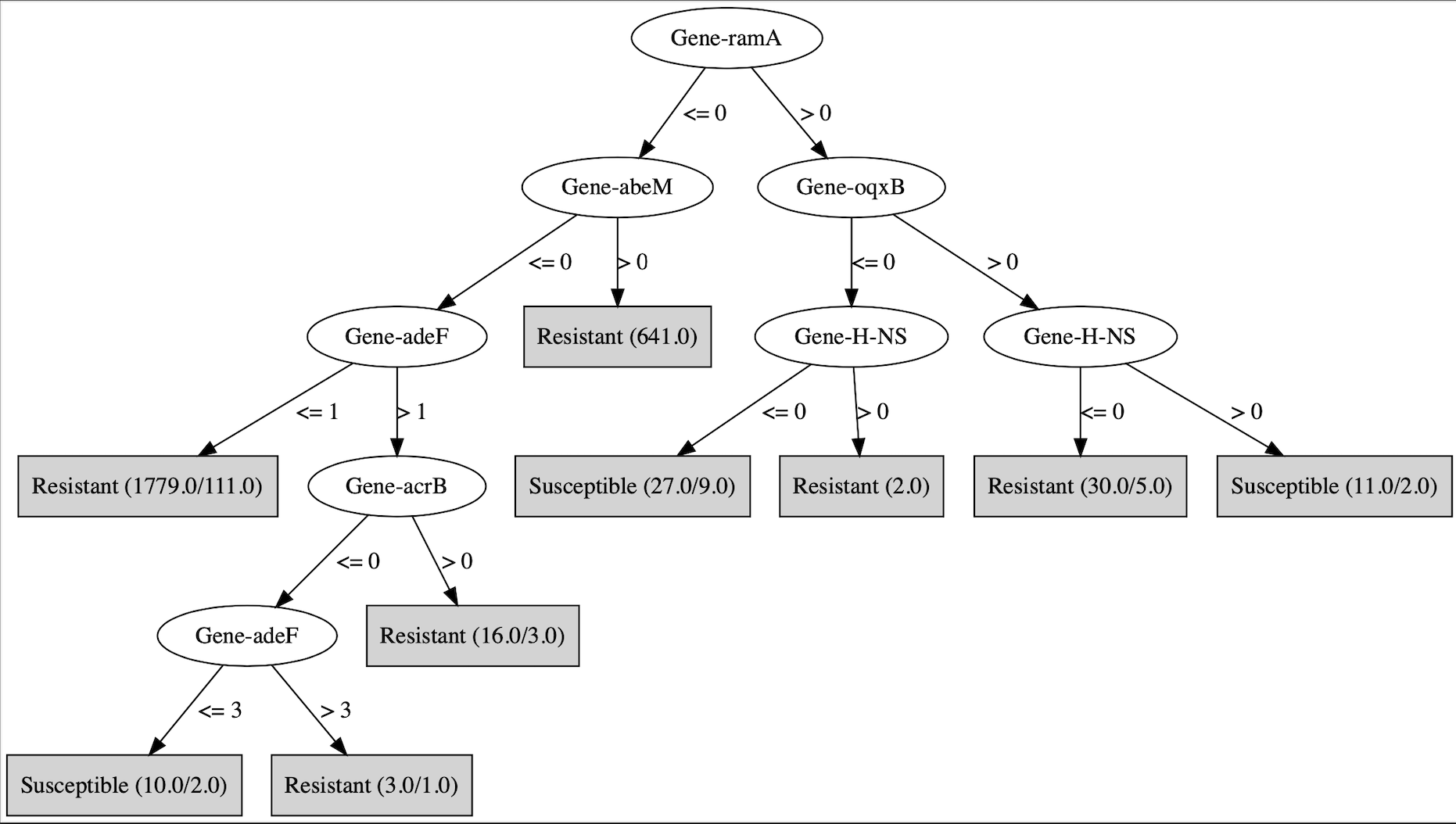

### PartR_Meropenem_RGI_specific.png

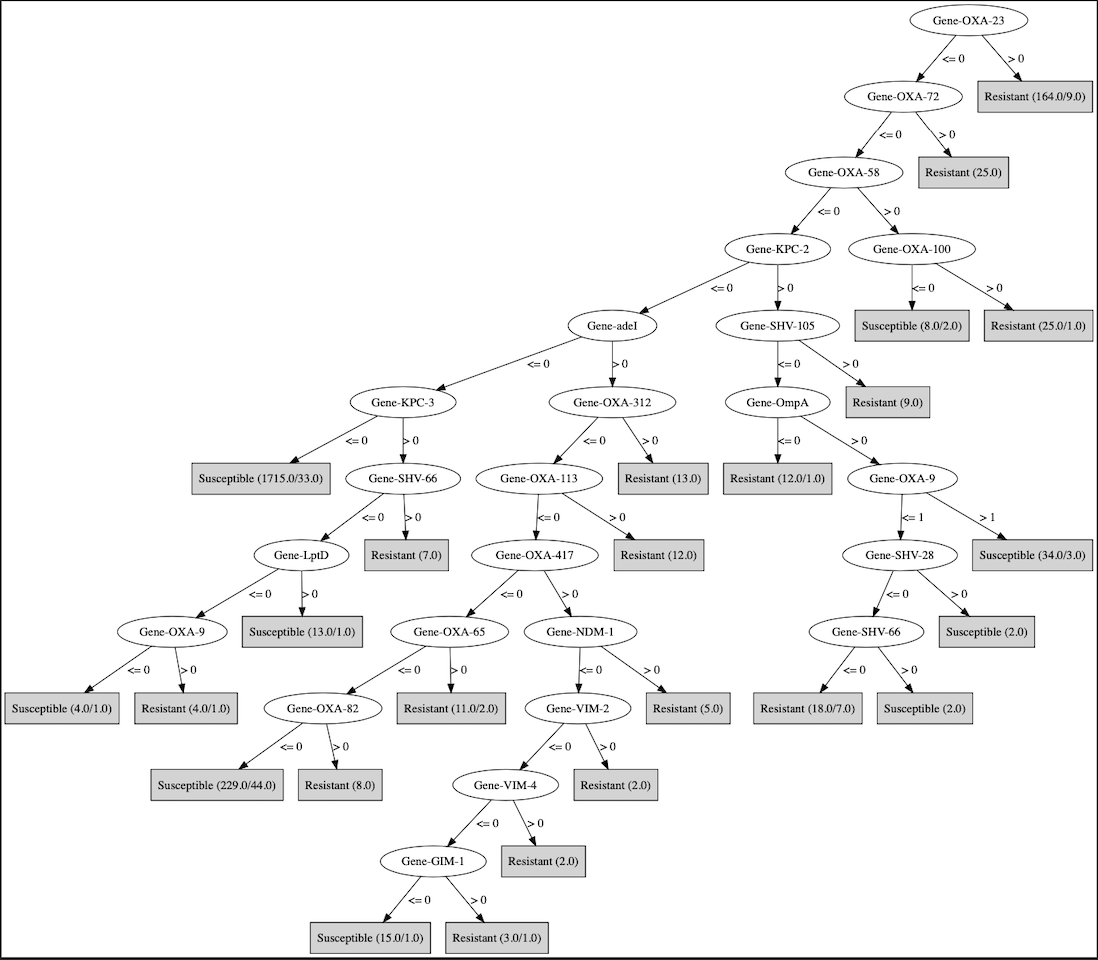

### PartS_Moxifloxacin_RGI_specific.png

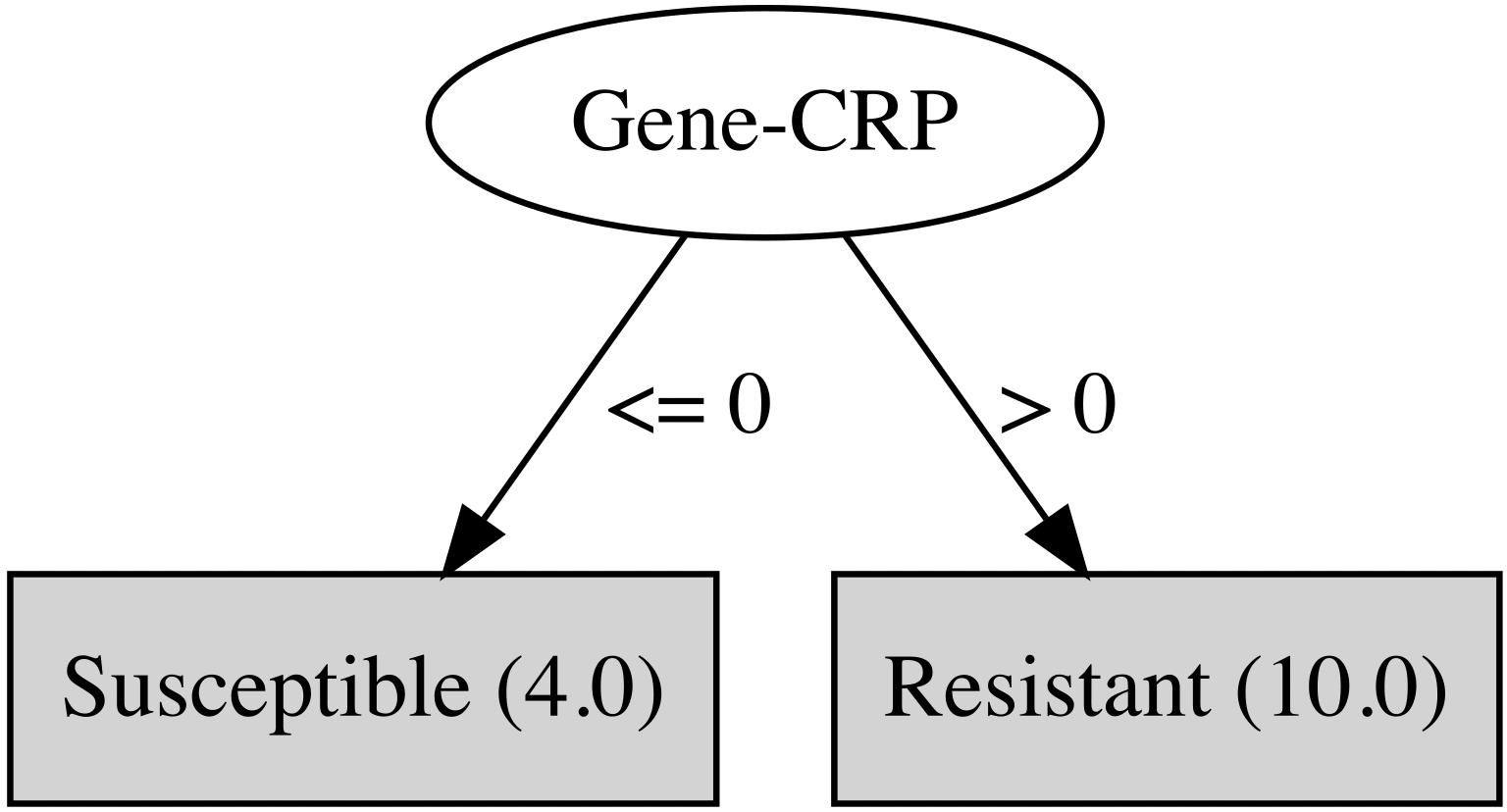

### PartT_Nitrofurantoin_RGI_specific.png

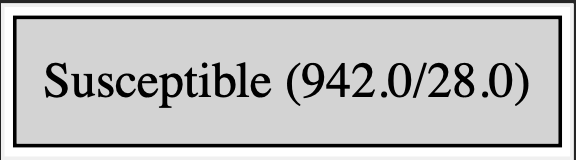

### PartU_Tetracycline_RGI_specific.png

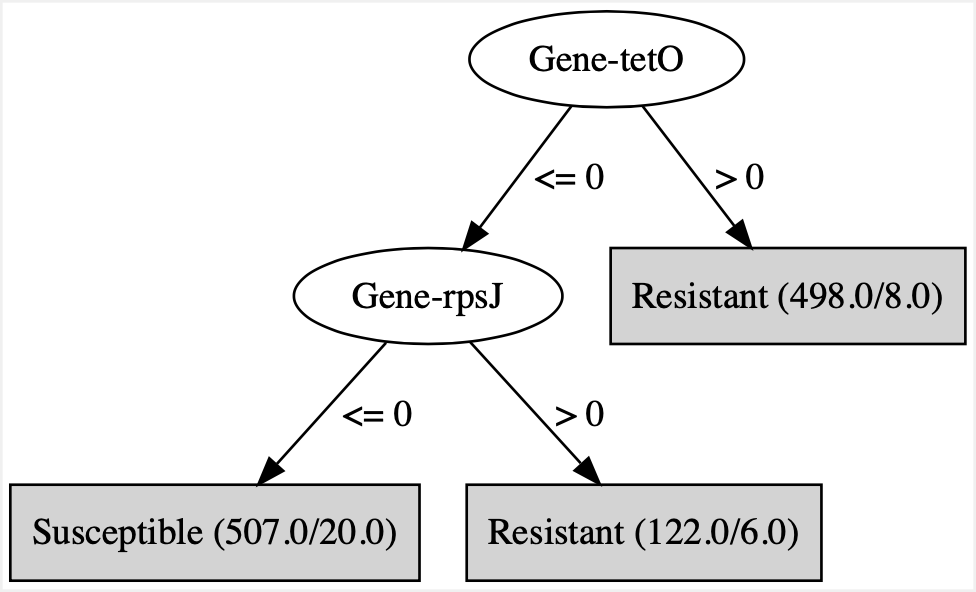

### PartV_Tigecycline_RGI_specific.png

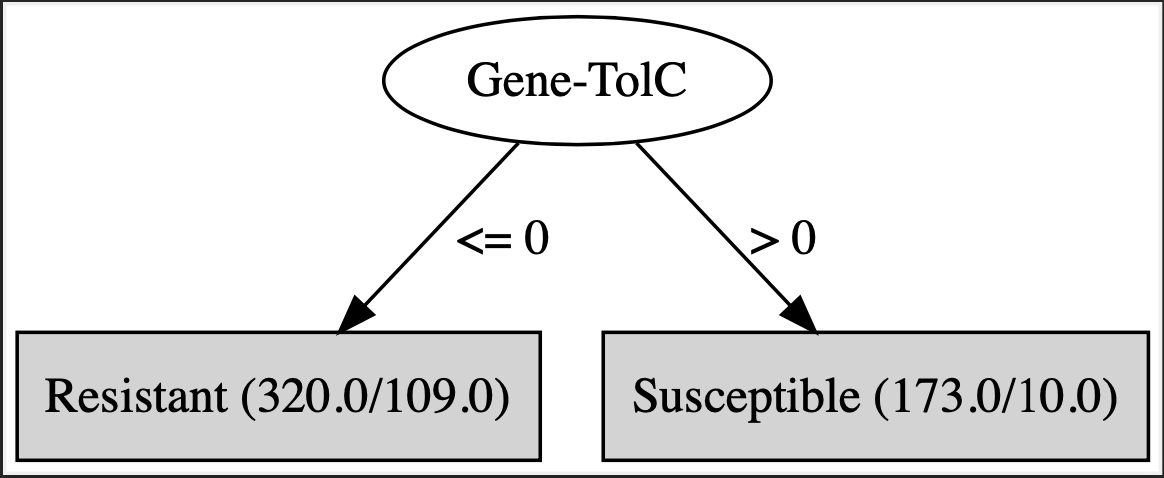

### PartW_Tobramycin_RGI_specific.png

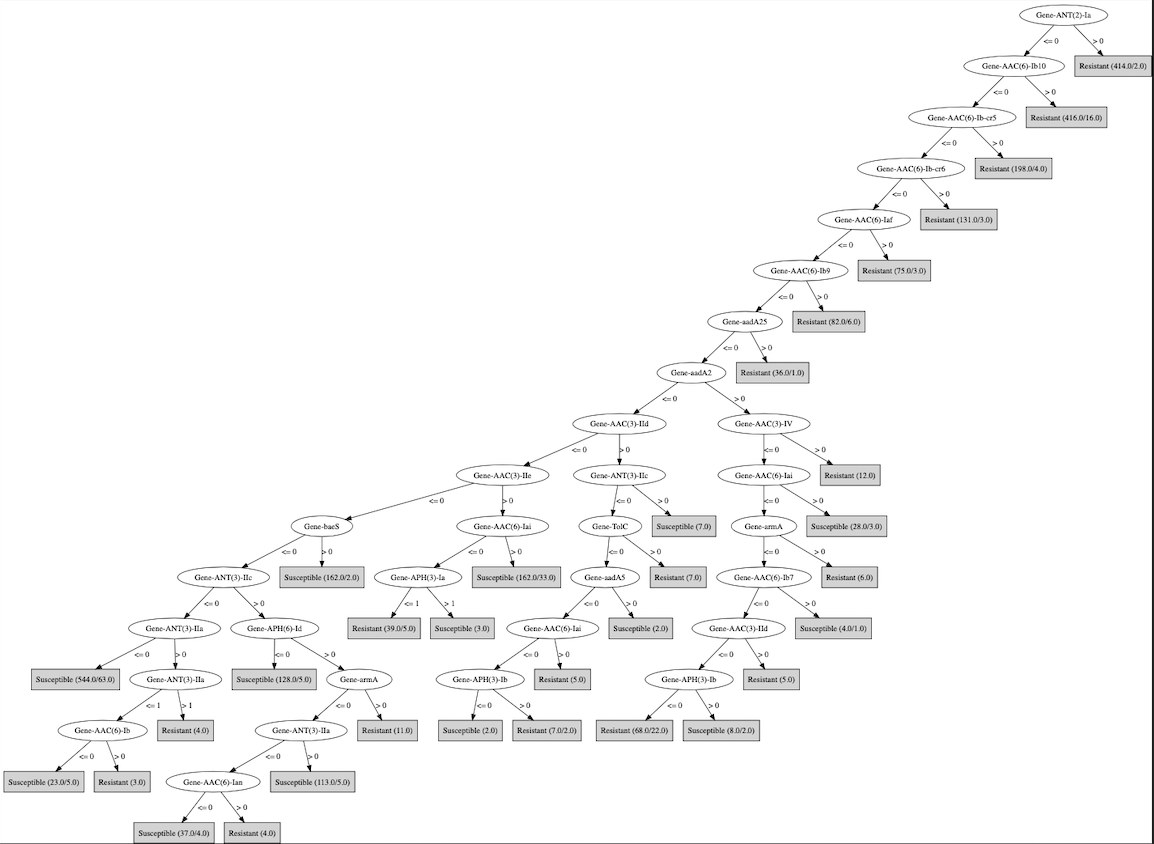

### Supplementary_Figure_1_Pheno_ratio.png

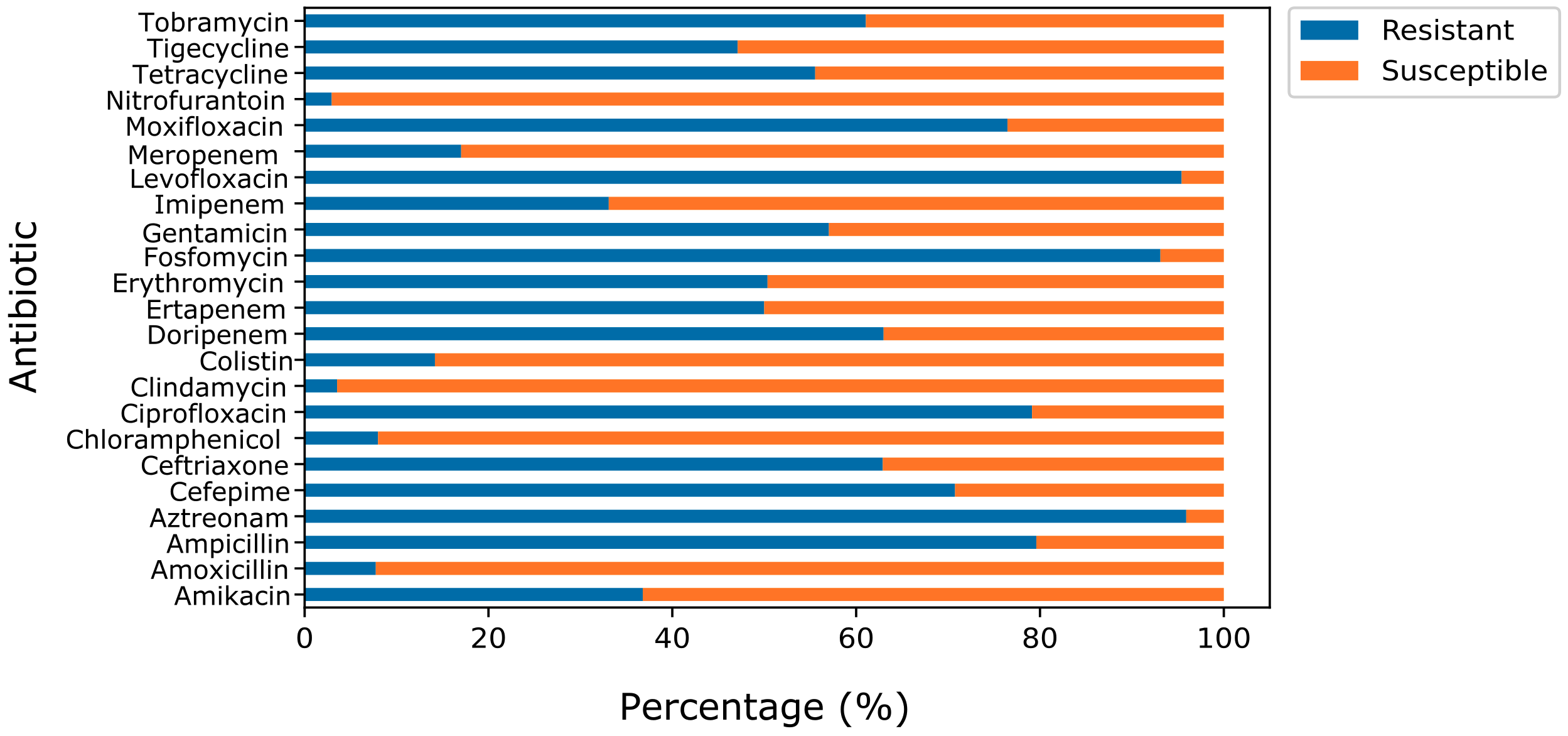

### Supplementary_Figure_2_RGI_schematic.png

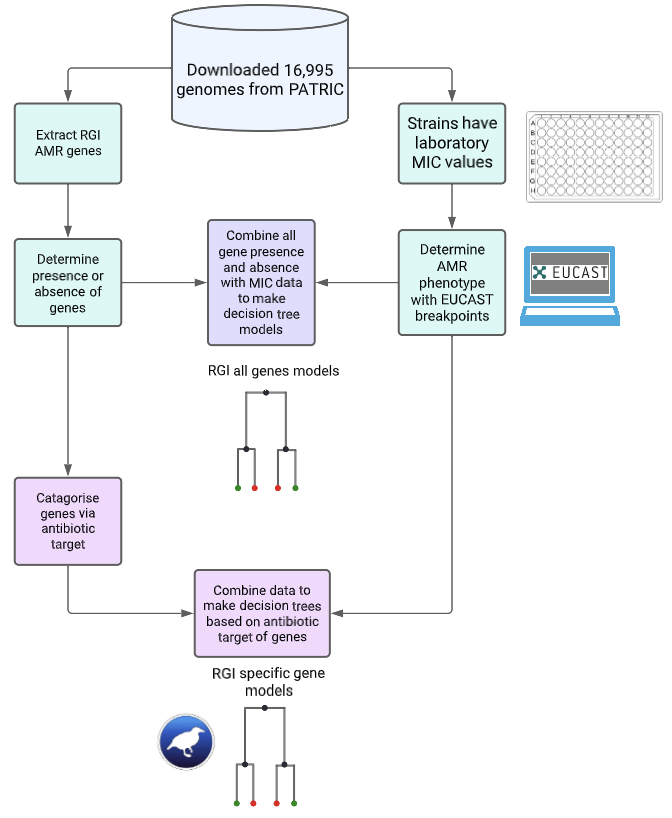

### Supplementary_Figure_6_Stacked_barcharts_cog.png

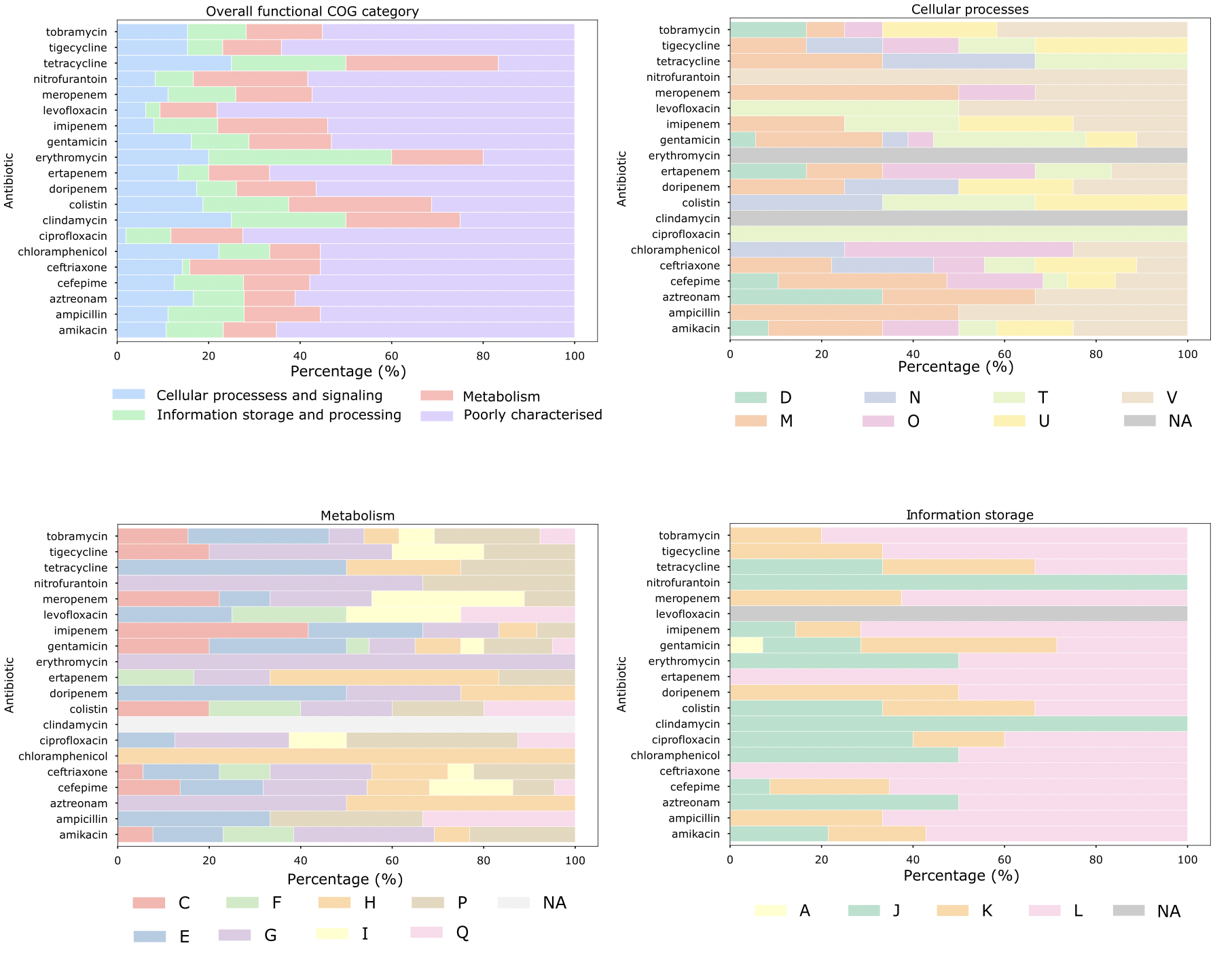

### Supplementary_Figure_7_Model_comparison_boxplot.pdf

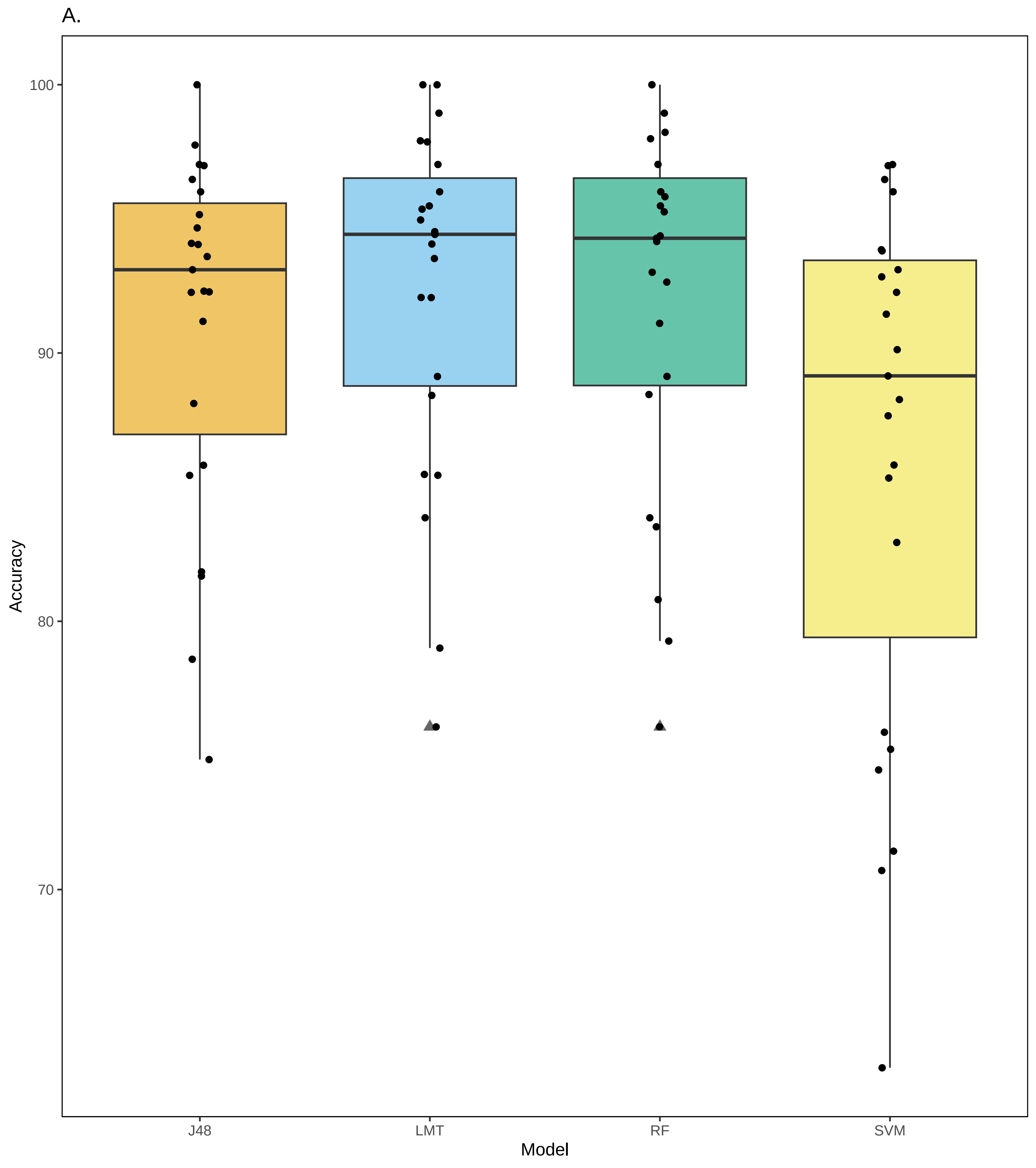

### Supplementary_Figure_10_Cytoscape_network_by_AB.png

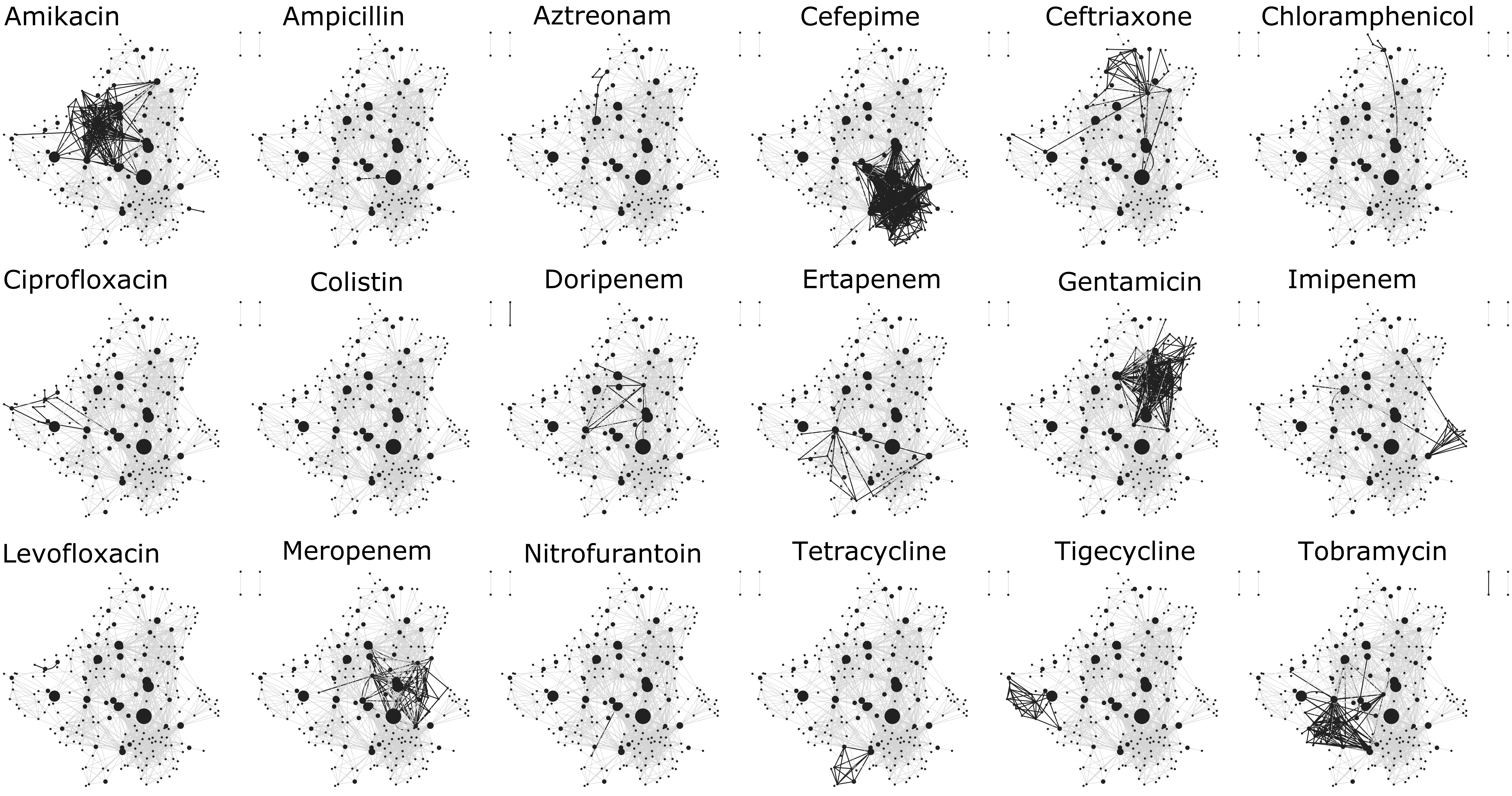

### Supplementary_Figure_11_Gene_network_by_drugclass.png

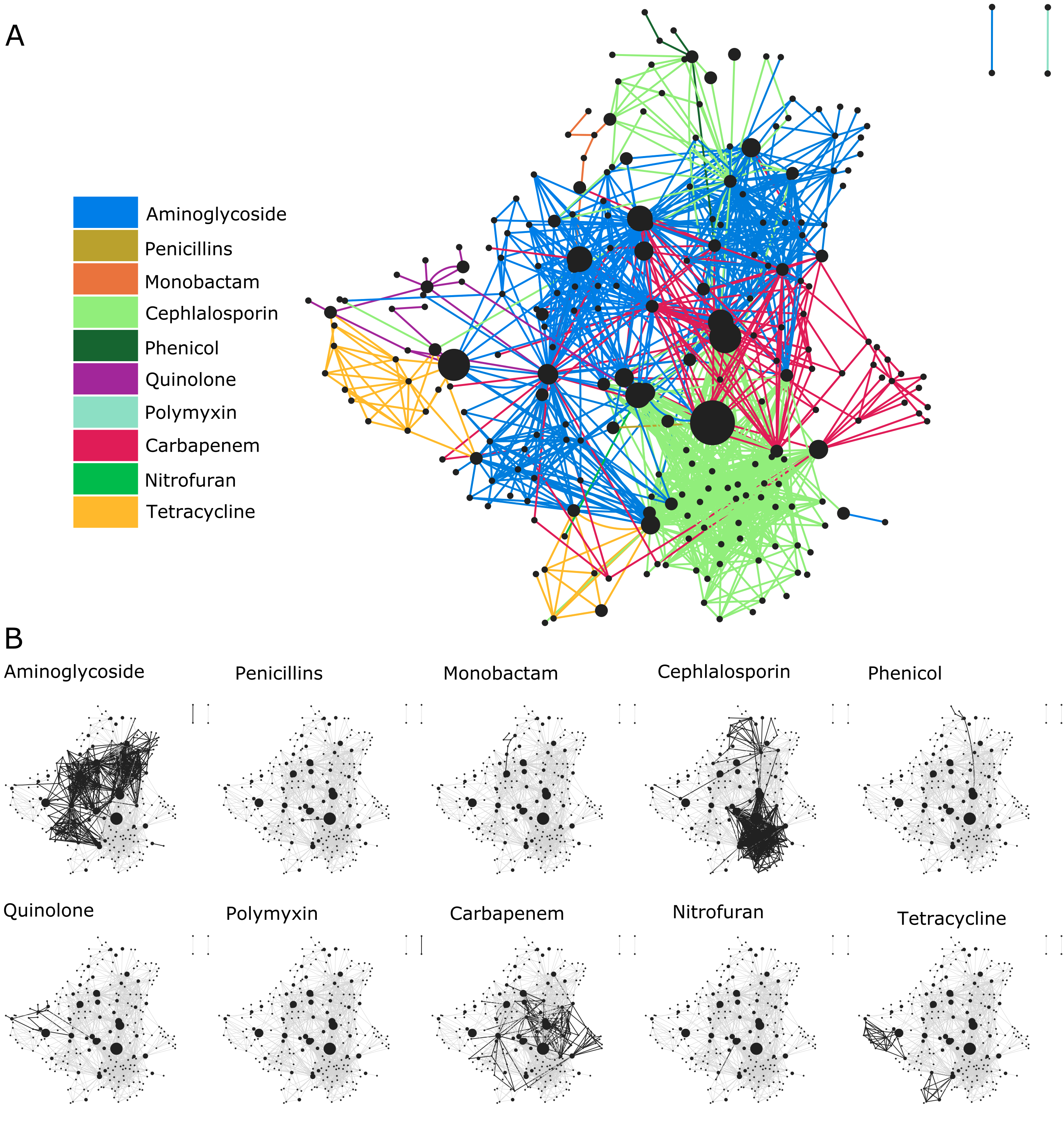
